# Integrative approaches to improve the informativeness of deep learning models for human complex diseases

**DOI:** 10.1101/2020.09.08.288563

**Authors:** Kushal K. Dey, Samuel S. Kim, Steven Gazal, Joseph Nasser, Jesse M. Engreitz, Alkes L. Price

## Abstract

Deep learning models have achieved great success in predicting genome-wide regulatory effects from DNA sequence, but recent work has reported that SNP annotations derived from these predictions contribute limited unique information for human complex disease. Here, we explore three integrative approaches to improve the disease informativeness of allelic-effect annotations (predicted difference between reference and variant alleles) constructed using several previously trained deep learning models: DeepSEA, Basenji and DeepBind (and a related machine learning model, deltaSVM). First, we employ gradient boosting to learn optimal combinations of deep learning annotations, using fine-mapped SNPs and matched control SNPs (on held-out chromosomes) for training. Second, we improve the specificity of these annotations by restricting them to SNPs implicated by (proximal and distal) SNP-to-gene (S2G) linking strategies, e.g. prioritizing SNPs involved in gene regulation. Third, we predict gene expression (and derive allelic-effect annotations) from deep learning annotations at SNPs implicated by S2G linking strategies — generalizing the previously proposed ExPecto approach, which incorporates deep learning annotations based on distance to TSS. We evaluated these approaches using stratified LD score regression, using functional data in blood and focusing on 11 autoimmune diseases and blood-related traits (average *N* =306K). We determined that the three approaches produced SNP annotations that were uniquely informative for these diseases/traits, despite the fact that linear combinations of the underlying DeepSEA, Basenji, DeepBind and deltaSVM blood annotations were not uniquely informative for these diseases/traits. Our results highlight the benefits of integrating SNP annotations produced by deep learning models with other types of data, including data linking SNPs to genes.

## Introduction

Deep learning models^1–8^ (and related machine learning models^9–11^) have shown considerable promise in predicting regulatory marks from DNA sequence, motivated by the well-documented role of non-coding variation in complex disease^12–18^. However, we recently showed that existing deep learning models provide limited unique information about complex disease when conditioned on a broad set of coding, conserved, regulatory and LD-related annotations^19^. Thus, further ideas are required in order for deep learning models to achieve their full potential in contributing to our understanding of complex disease.

Here, we explore three approaches for integrating different types of functional data to improve the disease informativeness of allelic-effect SNP annotations (predicted difference between reference and variant alleles) constructed using several previously trained deep learning models: DeepSEA^4^, Basenji^5^ and DeepBind^1^; for comparison purposes, we also consider a related machine learning model, deltaSVM^9^. First, we employ gradient boosting^20^ to learn optimal combinations of deep learning annotations, integrating these annotations with fine-mapped SNPs on held-out chromosomes from previous studies^21–23^. Second, we improve the specificity of deep learning/machine learning annotations by restricting them to SNPs linked to genes; we consider a broad set of proximal and distal SNP-to-gene (S2G) linking strategies, e.g. prioritizing SNPs involved in gene regulation^19, 24–32^. Third, we predict gene expression (and derive allelic-effect annotations) from deep learning annotations at SNPs implicated by S2G linking strategies, generalizing the previously proposed ExPecto approach^4^, which incorporates deep learning annotations based on distance to TSS. We consider either SNPs linked to all genes, or SNPs linked to genes in biologically important gene sets^19, 33^. We assessed the informativeness of the resulting annotations for disease heritability by applying stratified LD score regression (S-LDSC)^16^ to 11 autoimmune diseases and blood-related traits (average *N* =306K), conditional on a broad set of coding, conserved, regulatory and LD-related annotations from the baseline-LD model^34, 35^.

## Results

### Overview of Methods

We define an *annotation* as an assignment of a numeric value to each SNP with minor allele count ≥5 in a 1000 Genomes Project European reference panel^36^, as in our previous work^16^; we primarily focus on annotations with values between 0 and 1. Our annotations are derived from allelic-effect deep learning (or machine learning) annotations (predicted difference between reference and variant alleles of sequence-based predictions of functional annotations) from several recently developed models: DeepSEA^4^, Basenji^5^, DeepBind^1^ and deltaSVM^9^. DeepSEA employs a multi-class classification model to predict transcription factor and chromatin features by analyzing sequence data in a 1kb of human reference sequence around a SNP. Basenji employs a Poisson likelihood model to predict chromatin and CAGE profiles by analyzing 130kb of human reference sequence around each SNP using dilated convolutional layers. DeepBind fits a deep convolutional neural net model to sequences of varying length (14-101bp) to predict binding motifs of transcription factors and RNA-binding proteins. deltaSVM applies a gapped k-mer support vector machine (gkm-SVM^10, 11^) based classification method to sequences of length 10bp to predict profiles for transcription factors and chromatin features.

Our previous work^19^ focused on unsigned (absolute) allelic-effect annotations for DNase and three histone marks, H3K27ac, H3K4me1 and H3K4me3 (associated with active enhancers and promoters). Here, we integrate signed allelic-effect annotations for all features with other types of data - fine-mapped SNPs, SNPs linked to genes, and gene expression - to generate more disease-informative unsigned annotations. We have publicly released all new annotations analyzed in this study, along with open-source software for constructing the new annotations (see URLs).

First, we employ gradient boosting to integrate deep learning (or machine learning) annotations with fine-mapped SNPs (on held-out chromosomes) for blood-related traits from previous studies^21–23^ to generate boosted annotations representing an optimal combination of annotations. We use the XGBoost gradient boosting model^20^, and we train the gradient boosting model on even (respectively odd) chromosomes in order to construct annotations on odd (respectively even) chromosomes that are not used for training (to avoid overfitting); all parameter settings follow our previous work on AnnotBoost^37^, which has different goals.As input features, we use all of the deep learning (or machine learning) annotations from the pre-trained DeepSEA, Basenji, DeepBind and deltaSVM models, respectively. For comparison purposes, we also consider a simpler logistic regression model. Second, we improve the specificity of these annotations by restricting them to SNPs linked to genes using 10 (proximal and distal) SNP-to-gene (S2G) strategies^24–32, 38^ (Table 1). Third, we predict gene expression (and derive allelic-effect annotations) from deep learning annotations at SNPs implicated by S2G linking strategies, generalizing the previously proposed ExPecto approach^4^, which incorporates deep learning annotations based on distance to TSS.

**Table 1.** List of 10 S2G strategies: For each S2G strategy, we provide a brief description, indicate whether the S2G strategy prioritizes distal or proximal SNPs relative to the gene, and report its size (% of SNPs linked to genes). S2G strategies are listed in order of increasing size. Further details are provided in the Methods section.

| S2G strategy | Description | Distal/<br>Proximal | Size (%) |
| --- | --- | --- | --- |
| ABC | Inter-genic SNPs with distal enhancer-gene connections, assessed by Activity-By-Contact<sup>24,25</sup> across blood cell types. | Distal | 1.4 |
| TSS | SNPs in predicted Transcription start sites<sup>30,31</sup> overlapping Ensembl gene±5kb window. | Proximal | 1.6 |
| Coding | SNPs in coding regions | Proximal | 1.6 |
| ATAC | SNPs in ATAC-seq peaks >50% correlated to mouse expression across blood cell-types<sup>26</sup> (mapped to human). | Distal | 1.6 |
| eQTL | SNPs with fine-mapped causal posterior probability<sup>27</sup> (CPP) >0.001 in GTEx whole blood. | Distal<br>+Proximal | 2.4 |
| Roadmap | SNPs in predicted enhancer-gene links, assessed using Roadmap Epigenomics data<sup>28,32</sup>. | Distal | 3.2 |
| Promoter | SNPs in promoter regions. | Proximal | 4.3 |
| PC-HiC | Distal SNPs with Promoter Capture HiC (PC-HiC)<sup>29</sup> connections to promoter regions in blood cell-types. | Distal | 27 |
| 5kb | SNPs in ±5kb window around gene body. | Proximal | 53 |
| 100kb | SNPs in ±100kb window around gene body. | Distal | 81 |

We assessed the informativeness of the resulting annotations for disease heritability by applying stratified LD score regression (S-LDSC)^16^ to 11 independent blood-related diseases and traits (5 autoimmune diseases and 6 blood cell traits; average *N* =306K, Table S1) and meta-analyzing S-LDSC results across traits; we restricted our analyses to blood-related traits due to our focus on functional data in blood. We conservatively conditioned all analyses on a “baseline-LD-deep model” defined by 86 coding, conserved, regulatory and LD-related annotations from the baseline-LD model (v2.1)^34, 35^ and 14 additional jointly significant annotations from ref.^19^: 1 non-tissue-specific allelic-effect Basenji annotation, 3 Roadmap annotations, 5 ChromHMM annotations, and 5 other annotations (100 annotations total) (Table S2 and Table S3).

We used two metrics to evaluate the informativeness of individual annotations for disease heritability: enrichment and standardized effect size (*τ\**). Enrichment is defined as the proportion of heritability explained by SNPs in an annotation divided by the proportion of SNPs in the annotation^16^, and generalizes to annotations with values between 0 and 1^27^. Standardized effect size (*τ\**) is defined as the proportionate change in per-SNP heritability associated with a 1 standard deviation increase in the value of the annotation, conditional on other annotations included in the model^34^. Unlike enrichment, *τ\** quantifies effects that are unique to the focal annotation, thus, we use *τ\** as our primary metric. In our “marginal” analyses, we estimated *τ\** for each focal annotation conditional on the baseline-LD-deep annotations. In our “joint” analyses, we merged baseline-LD-deep annotations with focal annotations that were marginally significant after Bonferroni correction and performed forward stepwise elimination to iteratively remove focal annotations that had conditionally non-significant *τ^*^*values after Bonferroni correction, as in ref.^34^. Finally, in addition to the S-LDSC metrics enrichment and *τ\** (which evaluate *individual annotations*), we independently evaluated the combined joint model arising from our analyses using log*l*_SS_^39^, an approximate likelihood metric that evaluates a *heritability model* defined by a set of functional annotations, without running S-LDSC.

### DeepBoost annotations restricted to SNPs implicated by functionally informed S2G linking strategies are uniquely informative for autoimmune disease heritability

We developed a gradient boosting approach, DeepBoost, to learn optimal combinations of deep learning annotations (sequence-based predictions of functional annotations), using fine-mapped SNPs on held-out chromosomes for blood-related diseases/traits^21–23^ and matched control SNPs for training (Figure S1 and Methods). The input deep learning/machine learning annotations consisted of either 2,002 DeepSEA allelic-effect annotations^4^, or 4,229 Basenji allelic-effect annotations^5^, or 927 DeepBind allelic-effect annotations^1^ (based on TF and RBP motifs), or 1,329 deltaSVM allelic-effect annotations^9^ (trained on DHS and TF, which were reported to be the most informative features in recent work^40^). The DeepSEA, Basenji and deltaSVM models were based on tissue/cell-type-specific features spanning 127 tissues and cell types from Roadmap^41^; the DeepBind model was trained on non-tissue-specific features. We defined *published* allelic-effect annotations for each of these models as the maximum of the absolute allelic effects across relevant blood cell type features (or across all features for DeepBind, which is non-tissue-specific) (Methods).

The fine-mapped SNPs consisted of 8,741 fine-mapped autoimmune disease SNPs^21^ with causal probability *>* 0.0275, the threshold used by ref.^21^ (in secondary analyses, we also considered other sets of fine-mapped SNPs^22, 23^). DeepBoost uses decision trees to distinguish fine-mapped SNPs from matched control SNPs (with similar MAF and LD structure and local GC content) using an optimal combination of deep learning annotations; the DeepBoost model is trained using the XGBoost gradient boosting software^20^ (see URLs). DeepBoost attained an AUROC of up to 0.67 in distinguishing fine-mapped SNPs from control SNPs (highest = 0.67 for Basenji, second highest = 0.62 for DeepSEA), an encouraging result given the fundamental difficulty of this task (Table S4). The boosted allelic-effect annotations derived from DeepBoost (DeepSEAΔ- boosted, BasenjiΔ-boosted, DeepBindΔ-boosted, deltaSVMΔ-published; we use Δ to denote allelic-effect annotations) were only mildly correlated with published allelic-effect annotations as defined above (average *r*=0.16; Methods) (Figure S2). We also observed mild correlations between boosted annotations produced by the 4 models (average *r*=0.12; maximum of 0.32 between DeepSEAΔ-boosted and BasenjiΔ-boosted) (Figure S2). We determined that using logistic regression instead of XGBoost attained only slightly lower AUROC for each of the 4 models (average AUROC=0.612 for gradient boosting vs. 0.595 for logistic regression, difference=0.017; similar difference in secondary analyses of other fine-mapped SNP sets) (Table S4, Table S5). Given that a random classifier obtains an AUROC of 0.5, this can be viewed as a +17.9% improvement for gradient boosting (0.112*/*0.095 = 1.179); we believe this improvement is sufficient to justify the choice of gradient boosting in preference to logistic regression in our primary analyses; we also consider annotations constructed using logistic regression in our secondary analyses.

We broadly investigated which features of the DeepSEA, Basenji, DeepBind and deltaSVM models contributed the most to corresponding boosted annotations by applying Shapley Additive Explanation (SHAP)^42^, a widely used tool for pinpointing biological features underlying machine learning models^37, 43, 44^. For each model analyzed, we aggregated SHAP values across SNPs and primarily focused on the top 20 features, following ref.^37^ (see Data Availability for visualization of top 100 features); we caution that aggregating values across SNPs does not account for the wide variation in SHAP values in our analyses, and that the large number of features makes it difficult to delineate which features the observed improvements derive from. For DeepSEAΔ- boosted, top features included TF features in GM12878 and K562, two immune-related cell lines (Figure S3); for BasenjiΔ-boosted, top features included activating histone marks (H3K27ac and H3K4me3) in immune cell types (Figure S4); for DeepBindΔ- boosted, top features included the TF features *TBP*, *HOXA13* and *SP1* (Figure S5); for deltaSVMΔ-boosted, top features largely consisted of TF features in the immune cell type K562 (Figure S6). We also investigated which features of the DeepSEA, Basenji, DeepBind and deltaSVM models contributed the most to corresponding annotations constructed using logistic regression (instead of gradient boosting). We observed partial overlap with the top features from gradient boosting, including immune cell type features for DeepSEA and Basenji and the *HOXA13* TF feature for DeepBind (Table S6); *HOXA13* regulates genes associated with immune response, gap junction/cell adhesion, and pregnancy^45^. Finally, we investigated the impact of using only features from 27 blood cell types as input to our gradient boosting method (524 DeepSEA features or 479 Basenji features or 91 deltaSVM features; not applicable to DeepBind, which is non-tissue-specific). We determined that this attained only slightly lower AUROC than using all features (Table S7).

We assessed the informativeness for disease heritability of allelic-effect annotations constructed using DeepSEA, Basenji, DeepBind and deltaSVM. In our marginal analysis of disease heritability (across 11 autoimmune diseases and blood-related traits) using S-LDSC conditional on the baseline-LD-deep model, 2 of 4 published annotations (DeepSEAΔ-published, BasenjiΔ-published)and 1 of 4 boosted annotations (BasenjiΔ- boosted) were significantly enriched for heritability (after Bonferroni correction for 174 annotations tested; see Methods), with larger enrichments for the boosted annotations (Figure 1A, left panel, Figure 1C, left panel and Table S8); values of standardized enrichment (defined as enrichment scaled by the standard deviation of the annotation) are reported in Figure S7 and Table S9. However, none of these annotations attained Bonferroni-significant *τ\** values (although the BasenjiΔ-boosted annotation was FDR-significant) (Figure 1B, left panel, Figure 1D, left panel and Table S8). We constructed analogs of the DeepSEAΔ-boosted, BasenjiΔ-boosted, DeepBindΔ-boosted and deltaSVMΔ-boosted annotations using three other sets of fine-mapped SNPs: 4,312 fine-mapped inflammatory bowel disease SNPs^22^, 1,429 functionally fine-mapped SNPs for 14 blood-related UK Biobank traits^23, 46^, or the union of all 14,482 fine-mapped SNPs. The resulting annotations produced less disease signal than those constructed using the 8,741 fine-mapped autoimmune disease SNPs^21^ (Table S8).

**Figure 1.**
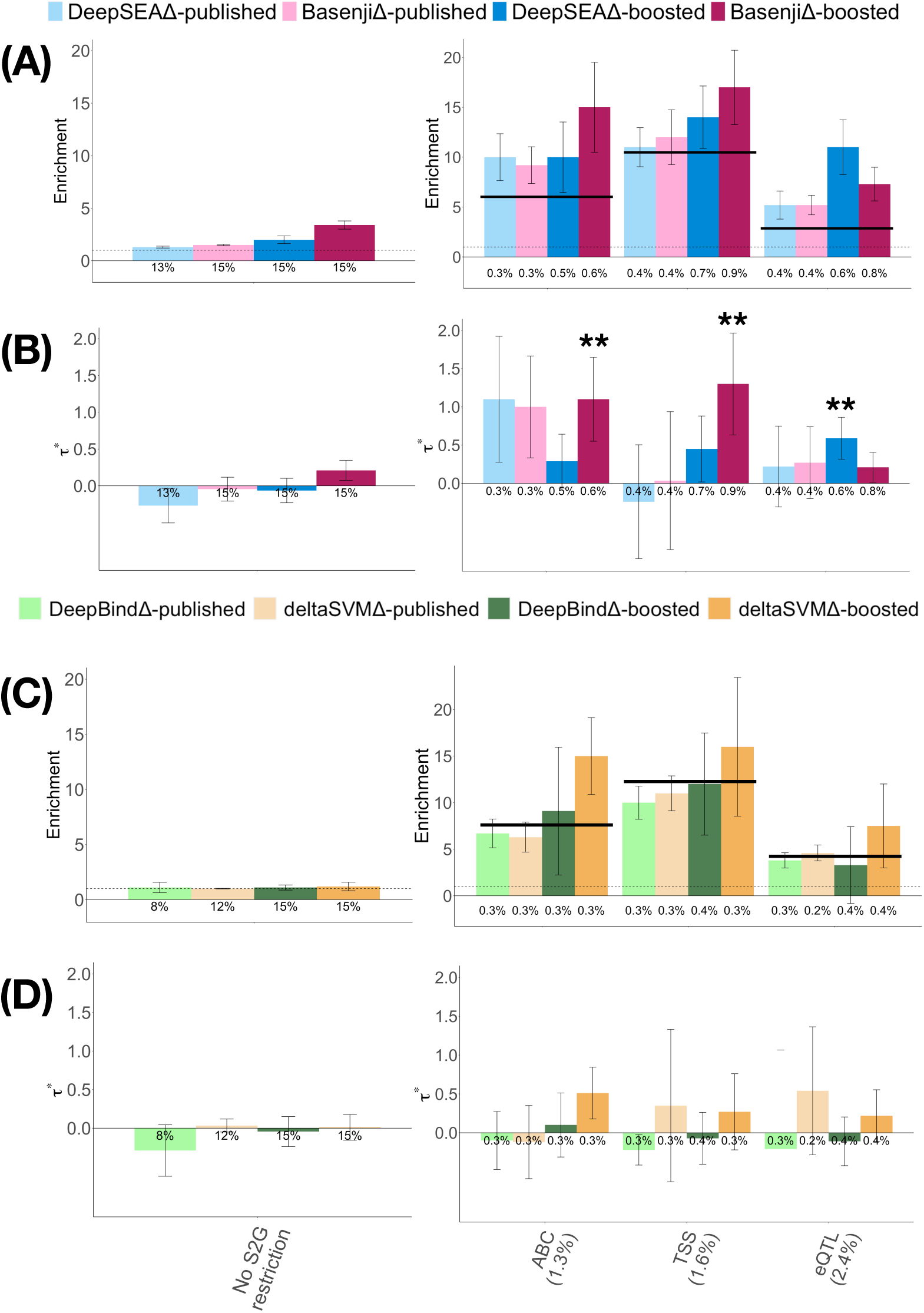
Disease informativeness of published and boosted allelic-effect deep learning annotations restricted to SNPs implicated by functionally informed S2G strategies: (A, Left panel) Heritability enrichment of published and boosted annotations based on the DeepSEA and Basenji models, conditional on the baseline-LD-deep model. Dashed horizontal line denotes no enrichment. (B, Left panel) Standardized effect size (*τ\**) of published and boosted DeepSEA and Basenji annotations, conditional on the baseline-LD-deep model. (A, Right panel) Heritability enrichment of published-restricted and boosted-restricted DeepSEA and Basenji annotations, conditional on the baseline-LD-deep-S2G model. Dashed horizontal line denotes no enrichment, solid horizontal lines denote enrichments of underlying S2G annotations. (B, Right panel) Standardized effect size (*τ\**) of published-restricted and boosted-restricted DeepSEA and Basenji annotations, conditional on the baseline-LD-deep-S2G model. (C, Left panel) Heritability enrichment of published and boosted annotations based on the DeepBind and deltaSVM models, conditional on the baseline-LD-deep model. Dashed horizontal line denotes no enrichment. (D, Left panel) Standardized effect size (*τ\**) of published and boosted DeepBind and deltaSVM annotations, conditional on the baseline-LD-deep model. (C, Right panel) Heritability enrichment of published-restricted and boosted-restricted DeepBind and deltaSVM annotations, conditional on the baseline-LD-deep-S2G model. Dashed horizontal line denotes no enrichment, solid horizontal lines denote enrichments of underlying S2G annotations. (D, Right panel) Standardized effect size (*τ\**) of published-restricted and boosted-restricted DeepBind and deltaSVM annotations, conditional on the baseline-LD-deep-S2G model. Results are meta-analyzed across 11 blood-related traits. The percentage under each bar denotes the size of the annotation (defined as average annotation value; equal to proportion of SNPs for binary annotations). ** denotes *P <* 0.05*/*174. Error bars denote 95% confidence intervals. Numerical results, including results for all 10 S2G strategies analyzed, are reported in Table S8 and Table S11.

We sought to improve the specificity of these annotations by restricting them to SNPs implicated by SNP-to-gene (S2G) linking strategies, e.g. prioritizing SNPs that may play a role in gene regulation; we define an S2G strategy as an assignment of 0, 1 or more linked genes to each SNP. We considered 10 S2G strategies capturing both proximal and distal gene regulation in blood, as in our previous work^38^ (see Methods and Table 1), and constructed 10 corresponding binary S2G annotations defined by SNPs linked to the set of all genes; the S2G annotations were only mildly positively correlated with each other (average *r* = 0.09; Figure S8). We defined *restricted* allelic-effect annotations as a simple product of allelic-effect annotations and S2G annotations. Due to correlations between allelic-effect annotations and S2G annotations (average *r* = 0.16; Figure S9), the size of a restricted allelic-effect annotation (defined as average annotation value; equal to proportion of SNPs for binary annotations) was generally larger than the product of the respective sizes of the underlying allelic-effect and S2G annotations (as would be expected if the two constituent annotations were independent); for example, the BasenjiΔ-boosted allelic effect annotation has size 15% and the ABC S2G annotation has size 1.4%, but the BasenjiΔ-boosted × ABC boosted-restricted allelic effect annotation has size 0.63%, which is lager than 15% × 1.4% = 0.21%. We evaluated 80 restricted allelic-effect annotations (8 allelic-effect annotations (4 published + 4 boosted) x 10 S2G annotations). We analyzed the restricted allelic-effect annotations conditional on a “baseline-LD-deep-S2G model” defined by 100 baseline-LD-deep annotations and 7 new S2G annotations from Table 1 that were not already included in the baseline-LD model (107 annotations total) (Table S2 and Table S10), to ensure that heritability enrichments that are entirely due to S2G annotations would not produce conditional signals.

In our marginal analysis of disease heritability using S-LDSC conditional on the baseline-LD-deep-S2G model, 48 of 80 annotations were significantly enriched for heritability (after Bonferroni correction for 174 annotations tested; see Methods), with larger enrichments for smaller annotations (Figure 1A right panel, Figure 1C right panel and Table S11); values of standardized enrichment were more similar across annotations (Table S12). Although published and boosted allelic-effect annotations were of similar size, the enrichments for boosted-restricted annotations across S2G strategies were higher on average (1.3x for DeepSEA, 1.5x for Basenji, 1.1x for DeepBind, 1.4x for deltaSVM) than the enrichments for published-restricted annotations. 3 of the boosted-restricted annotations (DeepSEAΔ-boosted × eQTL), BasenjiΔ-boosted × ABC and BasenjiΔ- boosted × TSS; no DeepBind or deltaSVM annotations) attained Bonferroni-significant *τ\** values (Figure 1B right panel, Figure 1D right panel and Table S11). (In comparison, when we conditioned only on the baseline-LD-deep model, 20 of the 80 annotations attained Bonferroni-significant *τ\** values (Table S13).

We jointly analyzed the 3 marginally significant annotations from the marginal analyses from Figure 1B, right panel by performing forward stepwise elimination to iteratively remove annotations that had conditionally non-significant *τ^∗^*values after Bonferroni correction. All 3 annotations were jointly significant in the resulting joint model, with joint effect sizes very similar to the conditional effect sizes from Figure 1B, right panel (Figure S10 and Table S14). All three annotations had joint *τ* >* 0.5; annotations with *τ^∗^>* 0.5 are unusual, and considered to be important^47^.

We investigated whether the boosted-restricted annotations would detect gene set-specific signals by further restricting them to SNPs linked to two biologically important gene sets: genes intolerant to loss-of-function (LoF) mutations^33^ (pLI) and genes with high PPI network connectivity to Enhancer-driven genes in blood^38^ (PPI-enhancer). We defined *gene set-specific* boosted-restricted annotations by replacing the S2G annotations (containing SNPs linked to all genes) with annotations containing SNPs linked to genes in the input gene set (Methods); we primarily focused on boosted-restricted annotations (instead of published-restricted annotations) because these were the restricted annotations that produced significant conditional signals in Figure 1B, right panel. We evaluated 80 gene set-specific boosted-restricted annotations (2 gene sets (pLI, PPI-enhancer) x 4 boosted allelic-effect annotations (BasenjiΔ-boosted, DeepSEAΔ-boosted, DeepBindΔ- boosted, deltaSVMΔ-boosted) x 10 S2G strategies). We analyzed the gene set-specific boosted-restricted annotations conditional on a “baseline-LD-deep-S2G-geneset” model defined by 107 baseline-LD-deep-S2G annotations and 8 jointly significant gene set-specific S2G annotations (Table S15 and Table S2), to ensure that heritability enrichments that are entirely due to the gene set-specific S2G annotations would not produce conditional signals.

In our marginal analysis of disease heritability using S-LDSC conditional on the baseline-LD-deep-S2G-geneset model, 41 of the 80 gene set-specific boosted-restricted annotations were significantly enriched for heritability (after Bonferroni correction for 174 annotations tested; see Methods), with larger enrichment for smaller annotations (Figure 2A and Table S16); values of standardized enrichment were more similar across annotations (Figure S11 and Table S17). 13 of the 80 annotations (3 DeepSEAΔ- boosted (PPI-enhancer), 3 BasenjiΔ-boosted (PPI-enhancer), 2 DeepBindΔ-boosted (PPI-enhancer), 3 deltaSVMΔ-boosted (PPI-enhancer), 1 DeepBindΔ-boosted (pLI) and 1 deltaSVMΔ-boosted (pLI)) attained conditionally Bonferroni-significant *τ^*^*values (Figure 2B and Table S16). We jointly analyzed these 13 annotations by performing forward stepwise elimination. The resulting joint model contained 2 jointly significant annotations, BasenjiΔ-boosted (PPI-enhancer) × ABC and BasenjiΔ-boosted (PPI- enhancer) Roadmap (Figure 2C and Table S18); both annotations had joint *τ* >* 0.5. Both annotations remained jointly significant, with very similar *τ^∗^*values, when further conditioned on the 3 jointly significant boosted-restricted annotations from Figure 1B, right panel (including the underlying BasenjiΔ-boosted *×* ABC annotation and the underlying BasenjiΔ-boosted *×* Roadmap annotation) (Table S18).

**Figure 2.**
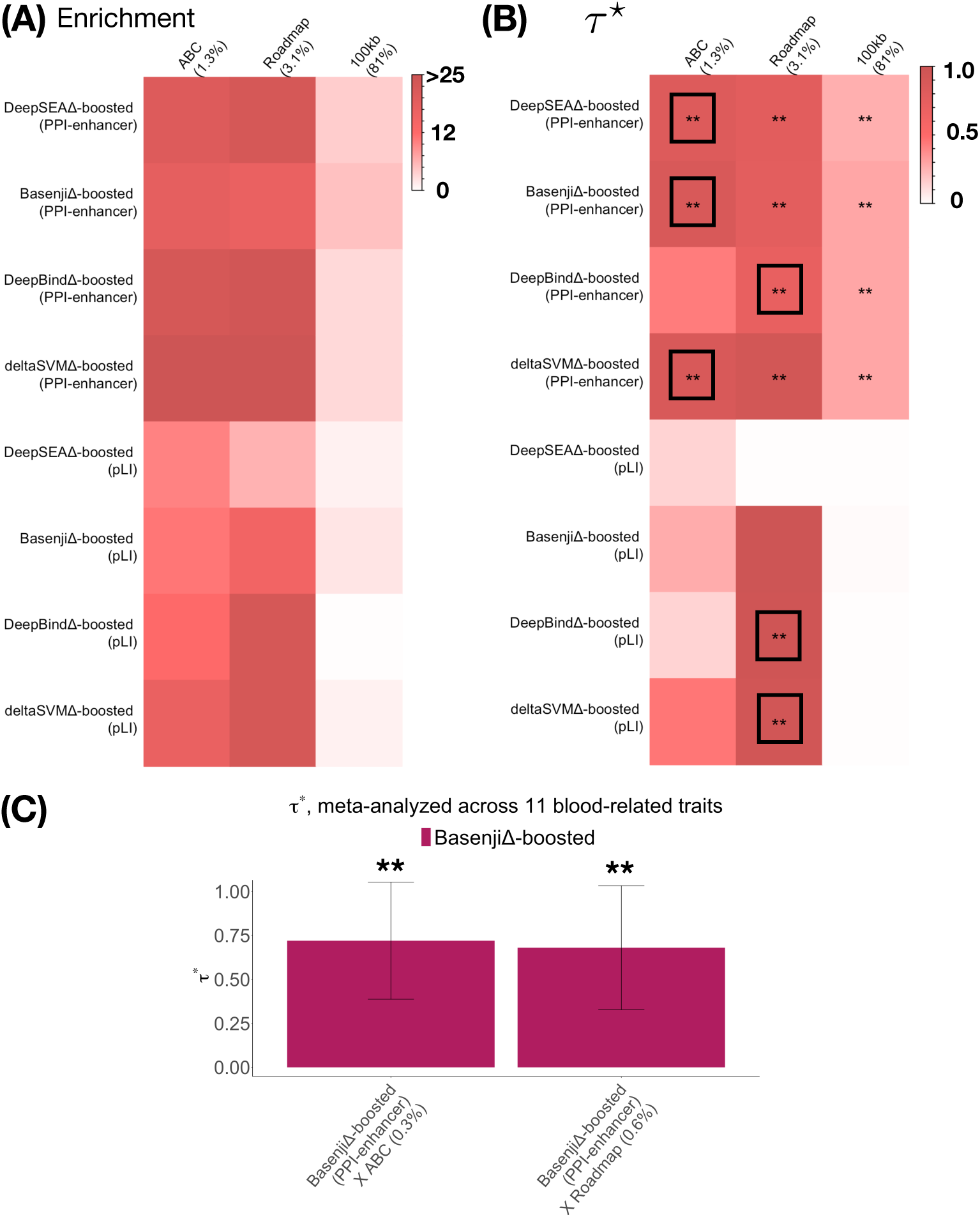
Disease informativeness of gene set-specific boosted-restricted annotations: (A) Heritability enrichment of gene set-specific boosted-restricted annotations based on the DeepSEA, Basenji, DeepBind and deltaSVM models, conditional on the baseline-LD-deep- S2G-geneset model. (B) Standardized effect size (*τ\**) of gene set-specific boosted-restricted DeepSEA, Basenji, DeepBind and deltaSVM annotations, conditional on the baseline-LD-deep- S2G-geneset model. (C) Standardized effect size (*τ\**) of the two jointly significant annotations, conditional on the baseline-LD-deep-S2G-geneset model plus both annotations. Results are meta-analyzed across 11 blood-related traits. *τ\** values less than 0 are displayed as 0 for visualization purposes. In panel C, the percentage under each bar denotes the size of the annotation (defined as average annotation value; equal to proportion of SNPs for binary annotations). ** denotes *P <* 0.05*/*174. Error bars denote 95% confidence intervals. In panel B, the black box in each row denotes the S2G strategy with highest *τ\**. Numerical results, including results for all 10 S2G strategies analyzed, are reported in Table S16 and Table S18.

We performed 3 secondary analyses. First, we repeated the analysis of restricted annotations using local GC-content (proportion of G and C nucleotides in a 1000bp window around each SNP) in addition to the S2G strategies, conditioning on the baseline-LD-deep-S2G model and the unweighted local GC-content annotation. The *τ^∗^*values for all 3 jointly significant restricted annotations from Figure 1B, right panel were nearly unchanged and remained Bonferroni-significant (Table S19); this implies that the unique disease signal in our restricted annotations cannot be explained by local GC-content. Second, we assessed the informativeness for disease heritability of allelic-effect annotations constructed using logistic regression (instead of gradient boosting). We determined that these annotations were less informative for disease heritability; in particular, only 1 of 3 annotations from Figure 1B, right panel (and no other annotations) attained conditionally Bonferroni-significant *τ^∗^*values (Table S20 and Table S21). Third, we repeated the analysis of gene set-specific restricted annotations using published-restricted annotations instead of boosted-restricted annotations. Marginal results were comparable to Figure 2B (14 annotations with Bonferroni-significant *τ^∗^*values; Table S22), but none of the gene set-specific published-restricted annotations annotations were significant conditional on the 2 jointly significant gene set-specific boosted-restricted from Figure 2C (Table S23).

We conclude that boosted deep learning allelic-effect annotations restricted to SNPs implicated by functionally informed S2G linking strategies are uniquely informative for autoimmune diseases and blood-related traits. All annotations that were uniquely informative for disease in our joint analyses were based on the DeepSEA and Basenji models, and we thus restrict our remaining analyses to these two deep learning models.

### Sequence-based deep learning predictions of gene expression informed by S2G linking strategies are uniquely informative for autoimmune disease heritability

We developed a new approach, Imperio, to predict gene expression from DNA sequence by using S2G strategies to prioritize deep learning annotations (sequence-based deep learning predictions of functional annotations) as features (Figure S12 and Methods). Imperio generalizes the ExPecto approach^4^, which prioritizes deep learning annotations as features based on distance to TSS. Specifically, Imperio uses regularized linear regression to fit optimal combinations of features predicting gene expression across 22,020 genes based on 2,002 DeepSEA or 4,229 Basenji deep learning annotations restricted to relatively common SNPs (MAF *>* 1%) linked to the target gene by 5 S2G strategies that are suitably large in size and generalizable to tissues beyond blood (5kb, 100kb, TSS, ABC and Roadmap Enhancer; Table 1) (2,002 × 5 or 4,229 × 5 features); the feature weights are independent of the target gene but dependent on the deep learning annotation and the S2G strategy (see Methods). In contrast, ExPecto fits optimal combinations of features based on 2,002 DeepSEA annotations restricted to 10 different functions of distance to TSS (using exponential decay), for a total of 2,002 × 10 features. We restricted our Imperio analyses to the DeepSEA and Basenji models, as all annotations that were uniquely informative for disease in our above joint analyses were based on these models. We focused on predicting gene expression in blood, due to the larger amount of data currently available for ABC and Roadmap Enhancer in blood cell types (however, our approach is generalizable to other tissues). We evaluated the accuracy of Imperio in predicting gene expression *across genes* on chromosome 8, which was withheld from Imperio training data (analogous to ref.^4^). We determined that Imperio attained similar predictive accuracy as ExPecto (Spearman correlation *ρ* = 0.76 (Basenji) and *ρ* = 0.72 (DeepSEA) with log RPKM expression, vs. *ρ* = 0.79 for ExPecto; Figure S13). The expression predictions were highly correlated between the Imperio and ExPecto models (average *ρ* = 0.82) (Figure S14), but the resulting allelic-effect annotations were less correlated, such that Imperio may contribute unique information (see below). The top significant features driving the Imperio model fit included Transcription Factor (TF) features for DeepSEA and CAGE features for Basenji (Table S24). When we compared the 5 Imperio models utilizing a single S2G strategy, TSS outperformed the other S2G strategies, but the resulting model fit (Spearman correlation *ρ* = 0.69 (Basenji) and *ρ* = 0.71 (DeepSEA) with log RPKM expression) was substantially worse than the model fit of the Imperio model utilizing all 5 S2G strategies (Table S25).

We used the Imperio allelic effects (signed predicted difference in expression between reference and variant alleles) to predict GTEx blood gene expression *across individuals* for each gene (see Methods). For each gene, we compared the Imperio prediction *r*^2^ to the total cis-SNP heritability of that gene, which represents an upper bound on the prediction *r*^2^ that can be attained using DNA sequence (because Imperio uses a (constrained) linear model to compute predictions; see Methods). Averaging across all 22,020 genes, Imperio predictions captured up to 82% of the total cis-SNP heritability on average (82% for Imperio-Basenji and 79% for Imperio-DeepSEA, vs. 75% for ExPecto; this analysis was not considered in ref.^4^) (Table S26). The Imperio prediction *r*^2^ closely tracked cis-SNP heritability (*ρ* = 0.83 for Imperio-DeepSEA, *ρ* = 0.84 for Imperio-Basenji across genes, vs. *ρ* = 0.81 for ExPecto) (Figure S15). Because disease heritability pertains to variation *across individuals*, the higher accuracy of Imperio in predicting gene expression variation across individuals may be expected to lead to annotations that are more informative for disease heritability.

We used the gene expression predictions from Imperio (DeepSEA and Basenji models) and ExPecto^4^ (DeepSEA model) to construct expression allelic-effect annotations (absolute value of the predicted difference in expression between reference and variant alleles) by summing allelic effects across genes linked by S2G strategies to the annotated SNP (see Methods). The Imperio training data excluded chromosome 8 (analogous to ref.^4^; see above), but did not exclude the target chromosomes on which allelic-effect annotations were constructed. However, this does not constitute overfitting, because the Imperio model was trained using reference sequence only. The Imperio-DeepSEA and Imperio-Basenji annotations were moderately correlated with each other (*r* = 0.54) and with ExPecto-DeepSEA (average *r* = 0.48) (Figure S16), such that each may contribute unique information. Furthermore, Imperio-DeepSEA and Imperio-Basenji annotations showed only mild correlation (average *r*=0.11) with boosted-restricted allelic effect annotations from previous section (Table S27). We analyzed the Imperio-DeepSEA, Imperio-Basenji and ExPecto-DeepSEA allelic-effect annotations conditional on the baseline-LD-deep-S2G-geneset model (see above; Table S2 and Table S15), for consistency with analyses of gene set-specific allelic-effect annotations (see below).

In our marginal analysis of disease heritability using S-LDSC conditional on the baseline-LD-deep-S2G-geneset model, all 3 annotations were significantly enriched for disease heritability (after Bonferroni correction for 174 annotations tested; see Methods), with larger enrichments for smaller annotations annotations (Figure 3A and Table S28); values of standardized enrichment were more similar across annotations (Table S29). One annotation, Imperio-Basenji, attained a Bonferroni-significant *τ^*^*value (Figure 3B and Table S28); the *τ\** value was very close to 0.5. This implies that Imperio-Basenji provides unique information about autoimmune diseases and blood-related traits. We note that the improvement of Imperio-Basenji vs. Expecto-DeepSEA derives both from the use of S2G strategies in Imperio (Imperio-DeepSEA vs. Expecto-DeepSEA) and the use of the Basenji model (Imperio-Basenji vs. Imperio-DeepSEA); however, statistical uncertainty precludes a precise quantification of the relative importance of these two factors.

**Figure 3.**
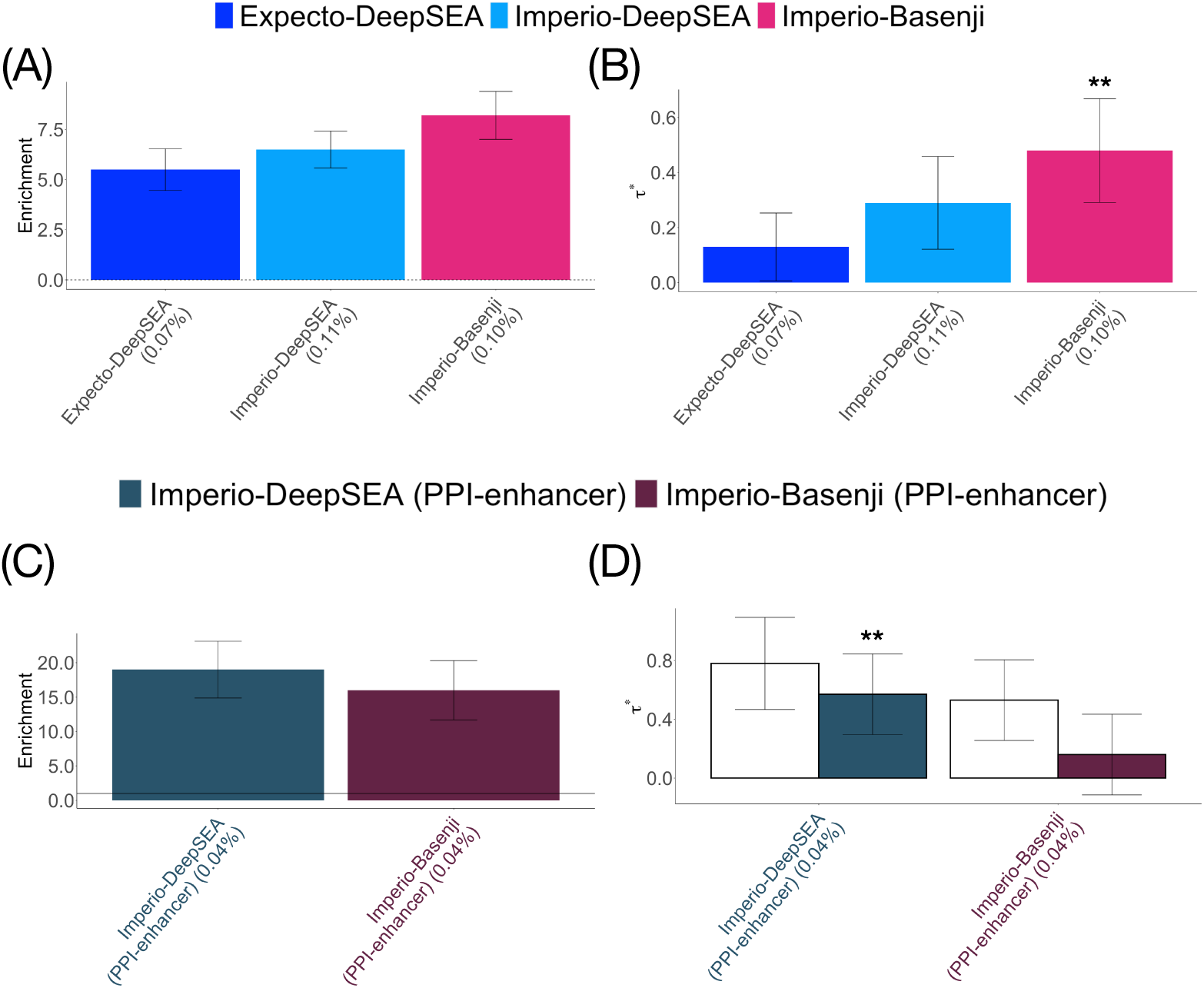
Disease informativeness of allelic-effect annotations based on predictions of gene expression from DNA sequence using S2G linking strategies to prioritize deep learning annotations as features: (A) Heritability enrichment of Imperio allelic-effect annotations, conditional on the baseline-LD-deep-S2G-geneset model. Dashed horizontal line denotes no enrichment. (B) Standardized effect size (*τ\**) of Imperio allelic-effect annotations, conditional on the baseline-LD-deep-S2G-geneset model. (C) Heritability enrichment of gene set-specific Imperio allelic-effect annotations, conditional on the baseline-LD-deep-S2G-geneset model. Dashed horizontal line denotes no enrichment. (D) Standardized effect size (*τ\**) of gene set-specific Imperio allelic-effect annotations, conditional on the baseline-LD-deep-S2G-geneset model. Results are meta-analyzed across 11 blood-related traits. The percentage under each bar denotes the size of the annotation (defined as average annotation value; equal to proportion of SNPs for binary annotations). ** denotes *P <* 0.05*/*174. Error bars denote 95% confidence intervals. Numerical results, including results for both pLI and PPI-enhancer gene sets, are reported in Table S28, Table S30 and Table S32.

We investigated whether the Imperio approach would detect gene-set specific signals by restricting Imperio to two biologically important gene sets: pLI^33^ and PPI-enhancer^38^ (see above). We defined *gene set-specific* allelic-effect annotations by restricting both the fitting of feature weights and the gene expression predictions to genes in the input gene set. Pairwise correlations between the 4 gene set-specific allelic-effect annotations ([Imperio-DeepSEA or Imperio-Basenji] x [pLI or PPI-enhancer]) (and the 3 non-gene set-specific allelic-effect annotations) are reported in Figure S16. We analyzed the gene set-specific allelic-effect annotations conditional on the baseline-LD-deep-S2G-geneset model (see above; Table S2 and Table S15).

In our marginal analysis of disease heritability using S-LDSC conditional on the baseline-LD-deep-S2G-geneset model, all 4 annotations were significantly enriched for disease heritability (after Bonferroni correction for 174 annotations tested; see Methods), with larger enrichments for smaller annotations annotations (Figure 3C and Table S30); values of standardized enrichment were more similar across annotations (Table S31). Two annotations, Imperio-DeepSEA (PPI-enhancer) and Imperio-Basenji (PPI-enhancer), attained Bonferroni-significant *τ\** values (Figure 3D and Table S30). In a joint analysis of both annotations, only Imperio-DeepSEA (PPI-enhancer) remained significant (Figure 3D and Table S32); the *τ\** value was larger than 0.5. Imperio-DeepSEA (PPI-enhancer) remained significant (with *τ* >* 0.5) when further conditioned on the Imperio-Basenji annotation from Figure 3B (Table S33).

We performed 5 secondary analyses. First, we fit an Imperio+ExPecto model using both Imperio (DeepSEA or Basenji) and ExPecto features. The Imperio+ExPecto allelic-effect annotations did not produce a significant disease signal conditional on the baseline-LD-deep-S2G-geneset model plus the Imperio-Basenji annotation from Figure 3B (Table S34). Second, we investigated a partially restricted gene set-specific Imperio approach by restricting either (a) the fitting of feature weights *or* (b) the gene expression predictions (but not both) to genes in the input gene set. None of the partially restricted gene set-specific annotations produced a significant disease signal conditional on the baseline-LD-deep-S2G-geneset model plus the two significant annotations from Figure 3B,D (Table S35 and Table S36). Third, we assessed whether the disease informativeness of Imperio could be explained by annotations defined by the number of genes linked to each SNP by each S2G strategy (see Methods). However, none of these annotations produced a significant disease signal conditional on the baseline-LD-deep-S2G-geneset model, either for all genes (Table S37) or when restricted to PPI-enhancer genes (Table S38). Fourth, we modified Imperio by constructing allelic-effect annotations using the *maximum* across genes proximal to the annotated SNPs, instead of the sum (see Methods). None of the modified annotations produced a significant disease signal conditional on the baseline-LD-deep-S2G-geneset model plus the two significant annotations from Figure 3B,D (Table S39). Fifth, we compared the Imperio annotations to MaxCPP-blood (Maximum across genes of fine-mapped eQTL Causal Posterior Probability) annotation^27^ constructed using GTEx whole blood gene expression data^48^. The MaxCPP-blood annotation was only weakly correlated with Imperio annotations (average *r* = 0.09) and did not produce a significant disease signal conditioned on the baseline-LD-deep-S2G- geneset model (Table S40), consistent with the fact that a related MaxCPP annotation based on a meta-analysis across tissues^27^ is already included in the baseline-LD model.

We conclude that allelic-effect annotations based on predictions of gene expression from DNA sequence using S2G linking strategies to prioritize deep learning annotations as features are uniquely informative for autoimmune diseases and blood-related traits.

### Combined joint model

We constructed a combined joint model containing annotations from the above analyses that were jointly significant, contributing unique information conditional on all other annotations. In detail, we merged the baseline-LD-deep-S2G-geneset model with 3 DeepBoost boosted-restricted allelic-effect annotations from Figure 1, 2 gene-set specific DeepBoost annotations from Figure 2, 1 Imperio gene expression prediction allelic-effect annotation from Figure 3B, and 1 gene-set specific Imperio annotation from Figure 3D, and performed forward stepwise elimination to iteratively remove annotations that had conditionally non-significant *τ\** values after Bonferroni correction. The resulting combined joint model contained 3 new annotations, including 1 DeepBoost annotation (BasenjiΔ-boosted × TSS) and the 2 Imperio annotations (Imperio-Basenji and Imperio-DeepSEA (PPI-enhancer)) (Figure 4 and Table S41). 2 of these annotations attained *τ^∗^>* 0.5: BasenjiΔ-boosted *×* TSS (1.1 *±* 0.29) and Imperio-DeepSEA (PPI-enhancer) (0.67 *±* 0.15); as noted above, annotations with *τ^∗^>* 0.5 are unusual, and considered to be important^47^. The combined *τ^∗^*^19, 49^ of the 3 annotations was high (1.7 *±* 0.3).

**Figure 4.**
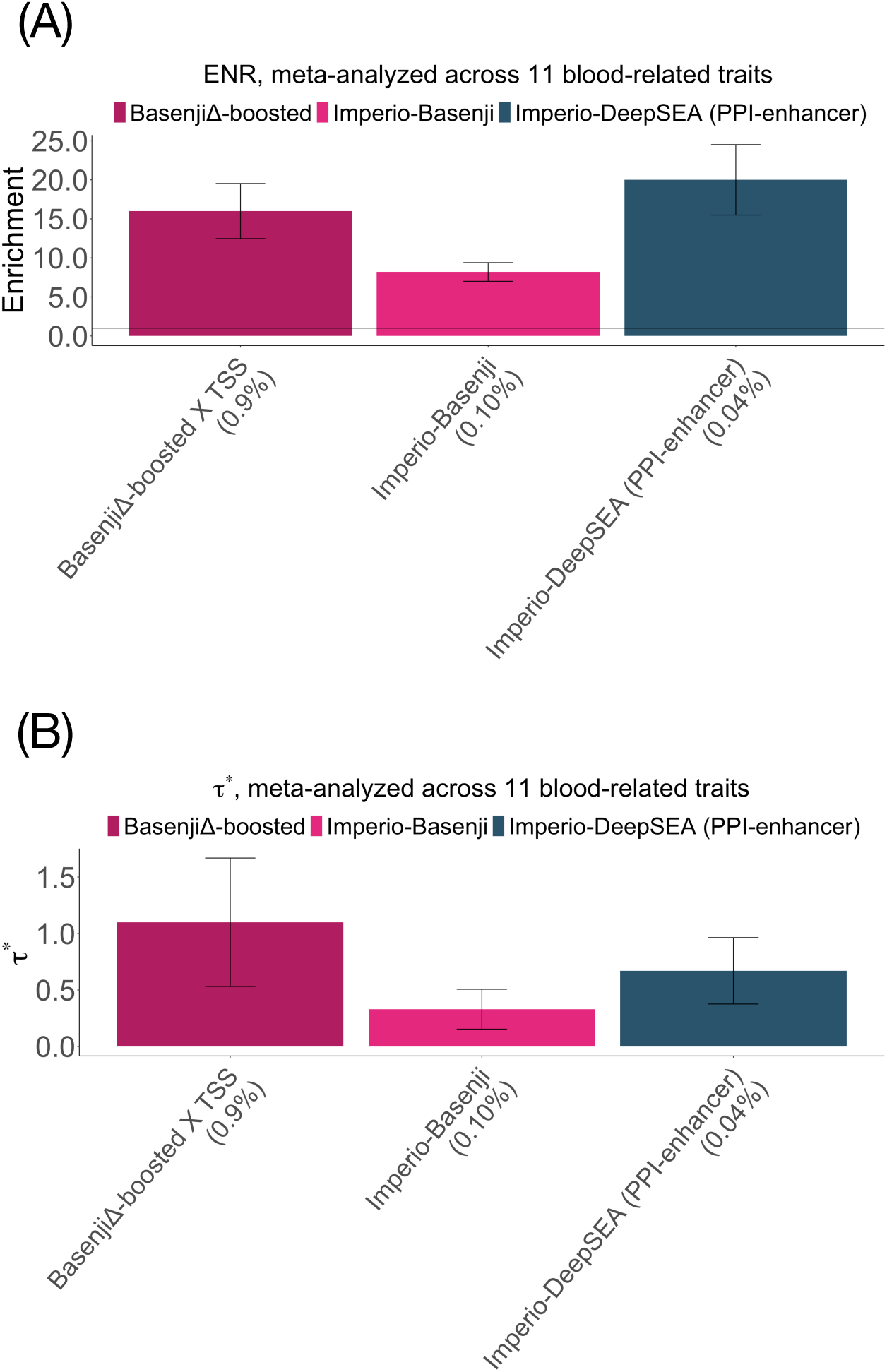
Combined joint model: (A) Heritability enrichment of 3 jointly significant annotations, conditional on the baseline-LD-deep-S2G-geneset model. (B) Standardized effect size (*τ\**) conditional on the baseline-LD-deep-S2G-geneset model plus the 3 jointly significant annotations. Results are meta-analyzed across 11 blood-related traits. The percentage under each bar denotes the size of the annotation (defined as average annotation value; equal to proportion of SNPs for binary annotations). Error bars denote 95% confidence intervals. Numerical results are reported in Table S41.

We independently evaluated the combined joint model of Figure 4 (and other models) by computing log*l*_SS_^39^, (an approximate likelihood metric that evaluates a heritability model defined by a set of functional annotations) relative to a model with no functional annotations (Δlog*l*_SS_), averaged across a subset of 6 blood-related traits (1 autoimmune disease and 5 blood cell traits) from the UK Biobank^46^ (Table S1). The combined joint model attained a +20.3% larger Δlog*l*_SS_ than the baseline-LD model (Table S42); +2.5% of the improvement derived from the 3 new annotations from Figure 4. The combined joint model also attained a +14.2% larger Δlog*l*_SS_ than the baseline-LD model (+2.2% of the improvement derived from the 3 new annotations from Figure 4) in a separate analysis of 24 non-blood-related traits from the UK Biobank (see Table S43 for list of traits) that had lower absolute log*l*_SS_ values (Table S42), implying that the value of the annotations introduced in this paper is not restricted to autoimmune diseases and blood-related traits.

We conclude that two types of allelic-effect annotations informed by S2G strategies—DeepBoost boosted-restricted annotations and Imperio gene expression prediction annotations—are jointly informative for autoimmune diseases and blood-related traits.

## Discussion

We have evaluated the contribution to autoimmune disease of SNP annotations constructed by integrating 4 sequence-based models - 3 deep learning approaches (DeepSEA, Basenji and DeepBind) and 1 machine learning approach (deltaSVM), with different types of functional data, including fine-mapped SNPs, SNP-to-gene linking strategies, gene expression data, and biologically important gene sets, using our DeepBoost and Imperio frameworks. We determined that boosted deep allelic-effect annotations restricted to SNPs implicated by functionally informed S2G linking strategies are uniquely informative for disease. We also determined that allelic-effect annotations based on prediction of gene expression from DNA sequence that were informed by S2G linking strategies are uniquely informative for disease, outperforming allelic-effect annotations from ExPecto^4^. We further determined that both DeepBoost and Imperio allelic-effect annotations were jointly informative for disease, resulting in an improved heritability model. All annotations that were uniquely informative for disease in our joint analyses using DeepBoost were based on the DeepSEA and Basenji models (and we thus restricted our Imperio analyses to these models). However, the DeepBind and deltaSVM models have performed well under other metrics: deltaSVM performed as well or better than DeepSEA in analyses of MPRA data^40^, and DeepSEA, DeepBind and deltaSVM performed similarly well in analyses of allele-specific transcription factor binding^50^.

Our work has several downstream implications. First, the DeepBoost and Imperio frameworks can be applied to other models beyond DeepSEA, Basenji, DeepBind and deltaSVM, and we anticipate that future deep learning models will benefit from these frameworks. Second, the accuracy of the Imperio framework in capturing cis-SNP heritability in blood suggests that it may be valuable to integrate Imperio gene expression predictions in other settings, such as transcriptome-wide association studies (TWAS)^51–53^ or mediated expression score regression (MESC)^54^. Third, our findings have immediate potential for improving functionally informed fine-mapping^23, 55–57^ (including experimental follow-up^58^), polygenic localization^23^, and polygenic risk prediction^59, 60^.

Our work has several limitations, representing important directions for future research. First, we focused our analyses on functional data in blood, and on blood-related diseases/traits; this choice was motivated by (i) the better representation of some S2G strategies, such as ABC and Roadmap Enhancer, in blood cell types than in other tissues, and (ii) the particularly large functional enrichments observed in autoimmune diseases and blood-related traits^16, 19, 27, 34^. However, it will be of interest to apply the DeepBoost and Imperio frameworks to other tissues and corresponding diseases/traits, once richer functional data becomes available. Second, we investigated the 10 S2G strategies separately, instead of constructing a single optimal combined strategy. A comprehensive evaluation of S2G strategies, and a method to combine them, will be provided elsewhere^61^. Third, our S-LDSC analyses are inherently focused on common variants, but deep learning models have also shown promise in prioritizing rare pathogenic variants^4, 8, 62^. The value of deep learning models for prioritizing rare pathogenic variants has been questioned in a recent analysis focusing on Human Gene Mutation Database (HGMD) variants^63^, meriting further investigation. Fourth, we focused here on deep learning models trained using human data, but models trained using data from other species may also be informative for human disease^26, 64^. Fifth, the forward stepwise elimination procedure that we use to identify jointly significant annotations^34^ is a heuristic procedure whose choice of prioritized annotations may be close to arbitrary in the case of highly correlated annotations. Nonetheless, our framework does impose rigorous criteria for conditional informativeness. Sixth, the large number of features (up to 4,229 features; Basenji model) makes it difficult to delineate which features the observed improvements derive from; this limitation is not unique to our work, as previous studies using deep learning models included a similarly large number of features^1, 2, 4, 5^.

Despite all these limitations, our findings improve the informativeness of deep learning models for autoimmune diseases and blood-related traits, and enhance our understanding of the sequence-mediated regulatory processes impacting these diseases/traits.

## Methods

### Genomic annotations and the baseline-LD model

We define a functional annotation as an assignment of a numeric value to each SNP with minor allele count ≥ 5 in a predefined reference panel (e.g., 1000 Genomes Project^36^; see URLs). Annotations can be either binary or continuous-valued (Methods). Our focus is on continuous-valued annotations (with values between 0 and 1) that are obtained by integrating deep learning models with functional data, including fine-mapped SNPs, SNP-to-gene linking strategies, gene expression data, and biologically important gene sets. Annotations that correspond to known or predicted function are referred to as functional annotations. The baseline-LD model (v.2.1) contains 86 functional annotations (see URLs). These annotations include binary coding, conserved, and regulatory annotations (e.g., promoter, enhancer, histone marks, TFBS) and continuous-valued linkage disequilibrium (LD)-related annotations.

### DeepSEA, Basenji, DeepBind and deltaSVM functional annotations

Deep learning/machine learning annotations were derived using three pre-trained Convolutional Neural Net (CNN) models: Basenji^5^, DeepSEA^2, 4^ (architecture from ref.^4^) and DeepBind^1^; and a Support Vector Machine (SVM) based machine learning model: deltaSVM^9^ (see URLs). Basenji is a Poisson likelihood model trained on original count data from 4, 229 cell-type specific histone mark, chromatin accessibility and FANTOM5 CAGE^65, 66^ annotations. Basenji uses dilated convolutional layers that allow scanning much larger contiguous sequence around a variant (≈ 130kb). DeepSEA is a classification based model trained on binary peak call data from 2, 002 cell-type specific TFBS, histone mark and chromatin accessibility annotations from the ENCODE^67^ and Roadmap Epigenomics^41^ projects with a sequence length of 1kb. DeepBind is a convolutional neural net model trained on 927 non-tissue-specific features based on 538 distinct transcription factors and 194 distinct RNA binding proteins. We restricted the deltaSVM model to 1,329 pre-trained sequence-based gapped k-mer support vector machine (gkm-SVM^10, 11^) features comprising of 699 ENCODE3 TFs, 317 DHS promoters and 313 DHS enhancers from Roadmap^9^ (see URLs), as these features were previously shown to be most informative in ref.^40^. For each SNP with minor allele count ≥ 5 in 1000 Genomes, we applied the pre-trained DeepSEA and Basenji models to the surrounding DNA sequence to compute both the prediction (at reference allele) and the predicted difference in probability between the reference and the alternate alleles. We call these the *variant-level annotations* and *allelic-effect annotations* respectively; this naming convention has been used previously^19^. The allelic-effect annotations are more interesting from a biological perspective as they are specific to a sequence-based predictive model like these deep learning models. We define a “‘published” allelic-effect annotation for each model by aggregating allelic effects across features. We defined DeepSEAΔ- published and BasenjiΔ-published annotations as the maximum absolute allelic effect across DNase, H3K27ac, H3K4me1 and H3K4me3 epigenomic marks in 27 blood cell types from Roadmap Epigenomics data^19, 41^. Similarly, we defined deltaSVMΔ-published as the maximum absolute effect across all features in the 27 blood cell types. Since DeepBind is non-tissue-specific, we defined DeepBoostΔ-published as the maximum absolute effect across all features considered..

### Boosted deep learning annotations using DeepBoost

DeepSEAΔ-published and BasenjiΔ-published represent a simple maximum of allelic-effect annotations across tissues and chromatin features. Here we introduce a gradient boosting approach to combine allelic-effect annotations across tissues and chromatin features. In detail, we train a classification model using decision trees, where each node in a tree splits SNPs into 2 classes (fine-mapped and control) using deep learning allelic-effect annotations from DeepSEA and Basenji models. The features in this classification model comprise of either allelic-effect annotations for 2,002 DeepSEA features or allelic-effect annotations for 4,229 Basenji features. We choose the control SNPs from non-finemapped SNPs matched for MAF, LD, local GC-content and the number of repeats distribution. MAF is based on the same reference panel (European samples from 100 Genomes Phase 3^36^), and LD is estimated by applying S-LDSC on all SNPs annotation (‘base’). The number of control SNPs were chosen equal to the number of fine-mapped SNPs. We used fine-mapped SNPs data related to blood traits from three sources^21–23^.

We used the Extreme gradient boosting (XGBoost) method implemented in the XGBoost software^20, 68^ with following model parameters: the number of estimators (200, 250, 300), depth of the tree (25, 30, 35), learning rate (0.05), gamma (minimum loss reduction required before additional partitioning on a leaf node; 10), minimum child weight (6, 8 ,10), and subsample (0.6, 0.8, 1); we optimized parameters by tuning hyper-parameters (a randomized search) with five-fold cross-validation. Two important parameters to avoid over-fitting are gamma and learning rate; we chose these values consistent with previous studies^69^, as in our previous work on AnnotBoost framework^37^.

The gradient boosting predictor is based on T additive estimators (T=200, 250, 300) and it minimizes the loss objective function *ℒ^t^* at iteration *t*.

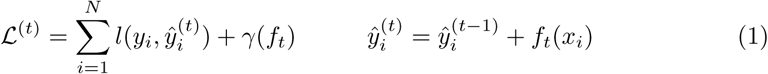

*f_t_* is an independent tree structure and *γ*(*f_t_*) is the complexity parameter. The final prediction from the gradient boosting model therefore is

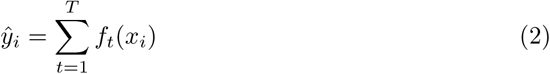

In order to avoid winner’s curse and overfitting, we use fine-mapped SNPs on odd (respectively even) chromosomes as training data to make predictions for even (respectively odd) chromosomes, as in our previous work on AnnotBoost^37^; thus, boosted annotations on a given chromosome are not informed by fine-mapped SNPs on that chromosome. We report the average AUROC of odd and even chromosome classifiers. The boosted annotations produced as output of the classifier are probabilistic in nature because of the logistic loss. We generate 4 boosted annotations, DeepSEAΔ-boosted, BasenjiΔ-boosted, DeepBindΔ-boosted and deltaSVMΔ-boosted, for each of 4 sets of fine-mapped SNPs, comprising of 8,741 fine-mapped autoimmune disease SNPs^21^, 4,312 fine-mapped inflammatory bowel disease SNPs^22^, 1,429 functionally fine-mapped SNPs for 14 blood-related UK Biobank traits^23, 46^, or the union of these 14,482 fine-mapped SNPs.

### Boosted-restricted deep learning annotations using S2G strategies

We define a SNP-to-gene (S2G) linking strategy as an assignment of 0, 1 or more linked genes to each SNP with minor allele count ≥ 5 in a 1000 Genomes Project European reference panel^36^. We intersect the 8 allelic-effect annotations from the previous subsections (DeepSEAΔ-published, BasenjiΔ-published, DeepBindΔ-published, deltaSVMΔ-published, DeepSEAΔ-boosted, BasenjiΔ-boosted, DeepBindΔ-boosted and deltaSVMΔ-boosted) with 10 S2G strategies used in ref.^38^ to generate 80 restricted allelic-effect annotations.

We explored 10 SNP-to-gene linking strategies in blood (Table 1). The proximal strategies included gene body ± 5kb; gene body ± 100kb; predicted TSS (by Segway^30, 31^); coding SNPs; and promoter SNPs (as defined by UCSC^70, 71^). The distal strategies included regions predicted to be distally linked to the gene by Activity-by-Contact (ABC) score^24, 25^ *>* 0.015 as suggested in ref.^24^ (see below); regions predicted to be enhancer-gene links based on Roadmap Epigenomics data (Roadmap)^28, 32, 41^; regions in ATAC-seq peaks that are highly correlated (*>* 50% as recommended in ref.^26^) to expression of a gene in mouse immune cell-types (ATAC)^26^; regions distally connected through promoter-capture Hi-C links (PC-HiC)^29^; and SNPs with fine-mapped causal posterior probability (CPP)^27^ *>* 0.001 (we chose this threshold to ensure that the SNP annotations generated after combining the gene scores with the eQTL S2G strategy were of reasonable size (0.2% of SNPs or larger) for all gene scores analyzed) in GTEx whole blood. (This is the threshold used throughout the analyses in our parallel study providing a comprehensive evaluation of S2G strategies^61^, which was initiated prior to the current study and has different goals.)

The boosted-restricted allelic-effect annotations were further restricted to SNPs linked to genes in two biologically important gene sets - pLI^33^ and PPI-enhancer^38^.

- **PPI-enhancer**: A binary gene score denoting genes in top 10% in terms of closeness centrality measure to the disease informative enhancer-regulated gene scores as defined in ref.^38^. To get the closeness centrality metric, we first perform a Random Walk with Restart (RWR) algorithm^72^ on the STRING protein-protein interaction (PPI network^73, 74^(see URLs) with seed nodes defined by genes in top 10% of the 4 enhancer-regulated gene scores defined in ref.^38^ with jointly significant disease informativeness (ABC-G^24, 25^, ATAC-distal^26^, EDS-binary^75^ and SEG-GTEx^76^). The closeness centrality score was defined as the average network connectivity of the protein products from each gene based on the RWR method.
- **pLI** : A probabilistic gene score with each gene graded by the probability of intolerance to loss of function mutations^33^.

We generate an additional 80 annotations by combining the 2 gene scores (pLI, PPI-enhancer) with 40 restricted boosted allelic-effect annotations for DeepSEA, Basenji, DeepBind and deltaSVM models and 10 S2G strategies.

### Imperio deep learning annotations using gene expression predictions informed by S2G strategies

We propose a new framework, Imperio, that for predicting gene expression from DNA sequence by using S2G strategies to prioritize deep learning annotations (sequence-based deep learning predictions of functional annotations) as features. This approach is analogous to the recent ExPecto framework^4^, but focuses on sequences around common variants linked to genes—either proximally or distally via enhancers, as in the Roadmap and ABC distal S2G strategies. We selected these two distal S2G strategies because they outperformed other distal strategies in blood in our previous work^38^. We integrate both DeepSEA and Basenji models with S2G strategies to predict gene expression. We consider a reduced set of 5 classes of S2G strategies: 5kb, 100kb, TSS, ABC and Roadmap Enhancer. We fit a elastic net regularized linear regression model to log RPKM expression data for gene *g*, *Y_g_* .

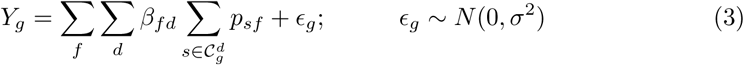

where *f* represents the chromatin mark features for the deep learning model (2,002 for the DeepSEA model and 4,229 for the Basenji model), *s* represents SNPs that are at least 1Kb apart ensuring relatively weaker correlation in their variant-effect or allelic-effect annotations, *d* represents a SNP-to-gene linking strategy, and 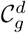 represents the set of all SNPs linked to gene *g* by the S2G strategy *d*. The 5 types of 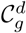 are:

- 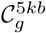: SNPs in a window of 5kb around gene *g*
- 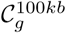: SNPs in a window of 100kb around gene *g*
- 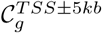: SNPs in a window of of *±*5*kb* around the TSS of gene *g*.
- 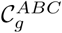: SNPs in regions linked to gene *g* by aggregation of Hi-C and enhancer marks data in 56 blood cell-types with a Acitivity-by-Contact (ABC) score of *>* 0.03.
- 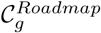: SNPs in Roadmap Enhancers linked to gene *g* in 27 blood cell-types.

*β_fd_*represents the model coefficient capturing the effect of each chromatin feature *f* and each S2G strategy *d* on the gene expression. *p_sf_* represents the variant-level prediction for chromatin feature *f* around SNP *i*. ϵ*_g_* represents white noise in the regression model. The model in Equation 3 is fitted by using Extreme gradient boosting (XGBoost) method. Following the training procedure in ExPecto, all genes except the ones in chromosome chr8 were used for training. The predictive performance of this approch is assessed on the holdout chromosome chr8.

We define the signed Imperio effect of each SNP as the sequence mediated effect on expression of a variant *s* and S2G strategy *d*.

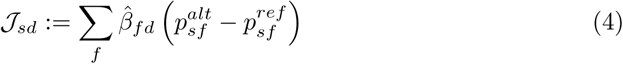

*𝒥_sd_* is the per-allele estimated change in expression caused by SNP *s* for any gene it is linked to through S2G strategy *d*. *𝒥_sd_* is treated as the atom for any Imperio based annotations we investigate.

The total absolute change in expression of gene *g* caused by SNP *s* and strategy *d* is given as follows.

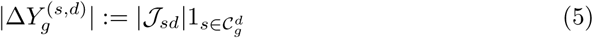

The total sequence mediated absolute predicted change by SNP *s* and S2G strategy *d* across all genes *g* is given by

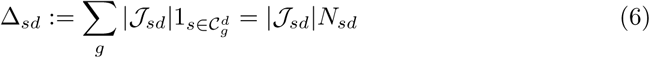

where *N_s_d* is the number of genes linked to SNP *s* by S2G strategy *d*.

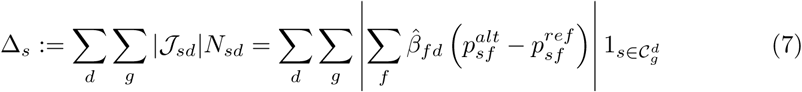

We adjust for the minor allele frequency (MAF) *p_s_* for each SNP *s* to adjust for per-allele effect sizes, as per ref^77^.

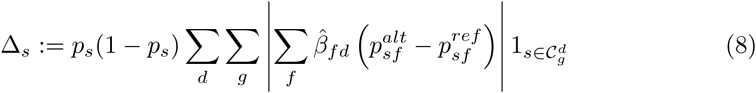

These Δ*_s_* scores were normalized to convert them to a probabilistic scale.

For a supplementary analysis, we also consider annotations that do not include the information of the number of genes linked to a SNP (*ξ_s_*).

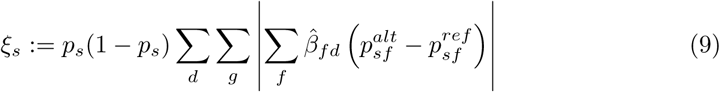

We analyze 3 annotations, 2 Imperio annotations, Imperio-DeepSEA and Imperio-Basenji, and ExPecto-DeepSEA.

### Predicting gene expression *across individuals* using Imperio

We use the Imperio effect of each SNP *s* in S2G strategy *d*, *𝒥_sd_* from Equation 4 (for either DeepSEA or the Basenji model) to define a gene specific Imperio score for each individual *n* and S2G strategy *d* as follows.

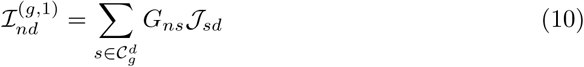

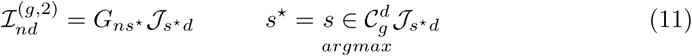

where *G_ns_* represents the number of risk alleles for individual *n* and the commonly varying SNP *s*.

Next we perform a regression on the normalized gene expression log RPKM data for individual *n* and gene *g*, *Y_ng_* with predictors given by 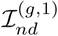 and 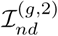.

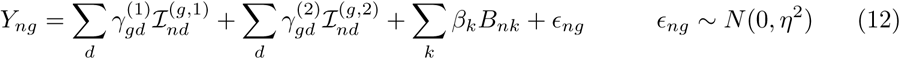

*B* denotes the covariates that are adjusted for in the model. We consider a total of 68 covariates including 5 principal components across samples, platform, gender and PCR amplification. In cases where there is only one SNP *s* in 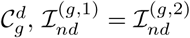 and only one of these predictors is used. This model provides an insight on relative contribution of different S2G strategies in explaining the inter-individual gene expression variation. The inter-individual Imperio model in Equation 12 is a linear model in risk alleles (*G_ns_*), similar to a gene expression cis-heritability model but with constrained parameters; thus, the cis-heritability represents an upper bound on the prediction *r*^2^ from the Imperio inter-individual prediction model.

We compute 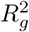, the proportion of variance explained by the predictor variables 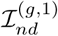 and 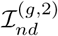 for all S2G strategies *d* and for gene *g*.

### Gene set-specific Imperio deep learning annotations

The Imperio model coefficients *β_fd_*in the previous section are fitted across all genes. However, different genes may have distinct sequence-mediated regulatory characteristics. Additionally, not all genes in blood are equally important. Therefore, we propose a gene-set specific Imperio model, where we perform the training of the model in Equation 3 over all genes *g* in a particular gene set 𝒢. We consider two gene sets, pLI^33^ and PPI-enhancer^38^ (see above).

The sequence-mediated expression effect of a variant *s* corresponding to gene set 𝒢 is given by

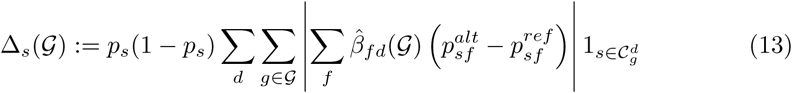

where 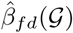 are the estimated model coefficients of *β_fd_*in Equation 3 fitted for genes in gene set 𝒢.

We analyze 4 annotations, combining Imperio models for DeepSEA and Basenji models with the PPI-enhancer and pLI gene sets.

We further define intermediate Imperio annotations by restricting either (a) the fitting of feature weights *or* (b) the gene expression predictions (but not both) to genes in the input gene set.

We define Imperio-sub-1 annotations generated by using all genes for fitting the model and gene sets for computing the expression allelic effects.

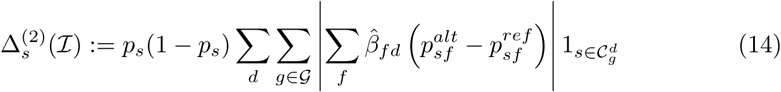

We define Imperio-sub-2 annotations generated by using genes in a geneset for fitting the model and all genes for computing the expression allelic effects

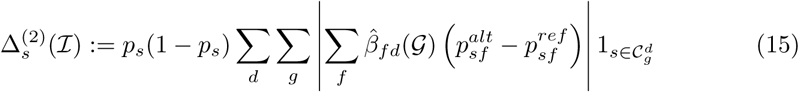

### Activity-by-Contact S2G strategy

The Activity-by-Contact (ABC)^24, 25^ (https://github.com/broadinstitute/ABC-Enhancer-Gene-Prediction) S2G strategy is determined by a predictive model for enhancer-gene connections in each cell type, based on measurements of chromatin accessibility (ATAC-seq or DNase-seq) and histone modifications (H3K27ac ChIP-seq), as previously described^24, 25^. We provide a brief summary of this approach, following ref.^24, 61^ (which contains further details). In a given cell type, the ABC model reports an “ABC score” for each element-gene pair, where the element is within 5 Mb of the TSS of the gene.

For each cell type, we:

- Called peaks on the chromatin accessibility data using MACS2 with a lenient p-value cutoff of 0.1.
- Counted chromatin accessibility reads in each peak and retained the top 150,000 peaks with the most read counts. We then resized each of these peaks to be 500bp centered on the peak summit. To this list we added 500bp regions centered on all gene TSS’s and removed any peaks overlapping blacklisted regions^78, 79^ (https://sites.google.com/site/anshulkundaje/projects/blacklists). Any resulting overlapping peaks were merged. We call the resulting peak set candidate elements.
- Calculated element Activity as the geometric mean of quantile normalized chromatin accessibility and H3K27ac ChIP-seq counts in each candidate element region.
- Calculated element-promoter Contact using the average Hi-C signal across 10 human Hi-C datasets as described below.
- Computed the ABC Score for each element-gene pair as the product of Activity and Contact, normalized by the product of Activity and Contact for all other elements within 5 Mb of that gene.

To generate a genome-wide averaged Hi-C dataset, we downloaded KR normalized Hi-C matrices for 10 human cell types (GM12878, NHEK, HMEC, RPE1, THP1, IMR90, HUVEC, HCT116, K562, KBM7). This Hi-C matrix (5 Kb) resolution is available here: ftp://ftp.broadinstitute.org/outgoing/lincRNA/average_hic/average_hic.v2.191020.tar.gz^25, 80^. For each cell type we performed the following steps.

- Transformed the Hi-C matrix for each chromosome to be doubly stochastic.
- We then replaced the entries on the diagonal of the Hi-C matrix with the maximum of its four neighboring bins.
- We then replaced all entries of the Hi-C matrix with a value of NaN or corresponding to Knight–Ruiz matrix balancing (KR) normalization factors ¡ 0.25 with the expected contact under the power-law distribution in the cell type.
- We then scaled the Hi-C signal for each cell type using the power-law distribution in that cell type as previously described.
- We then computed the “average” Hi-C matrix as the arithmetic mean of the 10 cell-type specific Hi-C matrices.

In each cell type, we assign enhancers only to genes whose promoters are “active” (i.e., where the gene is expressed and that promoter drives its expression). We defined active promoters as those in the top 60% of Activity (geometric mean of chromatin accessibility and H3K27ac ChIP-seq counts). We used the following set of TSSs (one per gene symbol) for ABC predictions: https://github.com/broadinstitute/ABC-Enhancer-Gene-Prediction/blob/v0.2.1/reference/RefSeqCurated.170308.bed. CollapsedGeneBounds.bed. We note that this approach does not account for cases where genes have multiple TSSs either in the same cell type or in different cell types.

For intersecting ABC predictions with variants, we took the predictions from the ABC Model and applied the following additional processing steps: (i) We considered all distal element-gene connections with an ABC score ≥ 0.015, and all distal or proximal promoter-gene connections with an ABC score ≥ 0.1. (ii) We shrunk the ~500-bp regions by 150-bp on either side, resulting in a ~200-bp region centered on the summit of the accessibility peak. This is because, while the larger region is important for counting reads in H3K27ac ChIP-seq, which occur on flanking nucleosomes, most of the DNA sequences important for enhancer function are likely located in the central nucleosome-free region. (iii) We included enhancer-gene connections spanning up to 2 Mb.

### Number of new annotations analyzed

For the Bonferroni correction, we corrected for 174 new annotations analyzed in our primary analyses (8 + 80 + 80 + 2 + 4 = 174). This choice is appropriate given the large number of potential secondary analyses, and is consistent with previous work^19^:

1. 8 genome-wide allelic-effect annotations: 4 published (DeepSEAΔ-published, BasenjiΔ-published, DeepBindΔ-published and deltaSVMΔ-published) and 4 boosted (DeepSEAΔ-boosted, BasenjiΔ-boosted, DeepBindΔ-boosted and deltaSVMΔ-boosted) annotations constructed using fine-mapped SNPs from ref.^21^ (vs. matched control SNPs). [Figure 1]
2. 80 restricted deep learning allelic-effect annotations corresponding to 4 published annotations (DeepSEAΔ-published, BasenjiΔ-published, DeepBindΔ-published and deltaSVMΔ-published) and 4 boosted annotations (DeepSEAΔ-boosted, BasenjiΔ- boosted, DeepBindΔ-boosted and deltaSVMΔ-boosted), restricted using 10 S2G strategies from Table 1. [Figure 1]
3. 80 gene set-specific restricted deep learning allelic-effect annotations, integrating DeepSEAΔ-boosted, BasenjiΔ-boosted, DeepBindΔ-boosted and deltaSVMΔ- boosted annotations, restricted using 10 S2G strategies from Table 1 with SNPs linked to genes specific to 2 gene scores (pLI and PPI-enhancer). [Figure 2]
4. 2 Imperio annotations (Imperio-DeepSEA, Imperio-Basenji) (we also analyzed 1 ExPecto-DeepSEA annotation from ref.^4^, but this is not a new annotation). [Figure 3]
5. 4 gene-set specific Imperio annotations combining Imperio-DeepSEA and Imperio-Basenji models with genes from 2 gene sets (pLI and PPI-enhancer). [Figure 3]

### Stratified LD score regression

Stratified LD score regression (S-LDSC) is a method that assesses the contribution of a genomic annotation to disease and complex trait heritability^16, 34^. S-LDSC assumes that the per-SNP heritability or variance of effect size (of standardized genotype on trait) of each SNP is equal to the linear contribution of each annotation

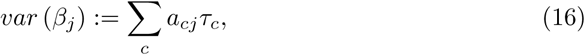

where *a_cj_* is the value of annotation *c* for SNP *j*, where *a_cj_* may be binary (0/1), continuous or probabilistic, and *τ_c_* is the contribution of annotation *c* to per-SNP heritability conditioned on other annotations. S-LDSC estimates the *τ_c_* for each annotation using the following equation

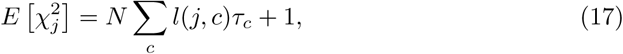

where 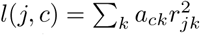 is the *stratified LD score* of SNP *j* with respect to annotation *c* and *r_jk_* is the genotypic correlation between SNPs *j* and *k* computed using data from 1000 Genomes Project^36^ (see URLs); N is the GWAS sample size.

We assess the informativeness of an annotation *c* using two metrics. The first metric is enrichment (*E*), defined as follows (for binary and probabilistic annotations only):

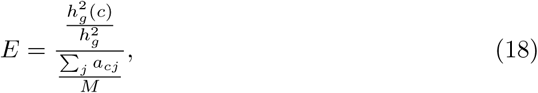

where 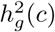 is the heritability explained by the SNPs in annotation *c*, weighted by the annotation values.

The second metric is standardized effect size (*τ\**) defined as follows (for binary, probabilistic, and continuous-valued annotations):

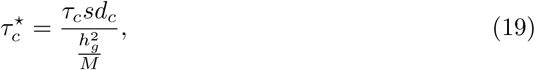

where *sd_c_* is the standard error of annotation *c*, 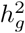 the total SNP heritability and *M* is the total number of SNPs on which this heritability is computed (equal to 5, 961, 159 in our analyses). 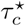 represents the proportionate change in per-SNP heritability associated to a 1 standard deviation increase in the value of the annotation.

### Combined ***τ\****

We use the combined *tau\** metric of ref.^19^, quantifying the conditional informativeness of a heritability model (combined *τ^∗^*, generalizing the combined *τ\** metric of ref.^49^ to more than two annotations. In detail, given a joint model defined by *M* annotations (conditional on a published set of annotations such as the baseline-LD model), we define

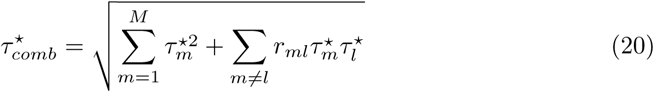

Here *r_ml_* is the pairwise correlation of the annotations *m* and *l*, and 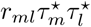 is expected to be positive since two positively correlated annotations typically have the same direction of effect (resp. two negatively correlated annotations typically have opposite directions of effect). We calculate standard errors for 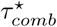 using a genomic block-jackknife with 200 blocks.

### Evaluating heritability model fit using SumHer log***l_SS_***

Given a heritability model (e.g. the baseline-LD model or the combined joint model of Figure 4), we define the Δlog*l*_SS_ of that heritability model as the log*l*_SS_ of that heritability model minus the log*l*_SS_ of a model with no functional annotations (baseline- LD-nofunct; 17 LD and MAF annotations from the baseline-LD model^34^), where log*l*_SS_^39^ is an approximate likelihood metric that has been shown to be consistent with the exact likelihood from restricted maximum likelihood (REML). We compute p-values for Δlog*l*_SS_ using the asymptotic distribution of the Likelihood Ratio Test (LRT) statistic: *−*2 log*l*_SS_ follows a *χ*^2^ distribution with degrees of freedom equal to the number of annotations in the focal model, so that *−*2Δlog*l*_SS_ follows a *χ*^2^ distribution with degrees of freedom equal to the difference in number of annotations between the focal model and the baseline-LD-nofunct model. We used UK10K as the LD reference panel and analyzed 4,631,901 HRC (haplotype reference panel^81^) well-imputed SNPs with MAF ≥ 0.01 and INFO≥ 0.99 in the reference panel; We removed SNPs in the MHC region, SNPs explaining *>* 1% of phenotypic variance and SNPs in LD with these SNPs.

### Data Availability

All DeepBooost and Imperio annotations are available at https://alkesgroup.broadinstitute.org/LDSCORE/DeepLearning/Dey_DeepBoost_Imperio/. The deep learning allelic effect SNP level annotations for DeepSEA, Basenji, DeepBind and deltaSVM models are available at https://alkesgroup.broadinstitute.org/LDSCORE/DeepLearning/. This work used summary statistics from the UK Biobank study (http://www.ukbiobank.ac.uk/). The summary statistics for UK Biobank is available online (https://data.broadinstitute.org/alkesgroup/UKBB/). The 1000 Genomes Project Phase 3 data are available at ftp://ftp.1000genomes.ebi.ac.uk/vol1/ftp/release/20130502. The baseline-LD annotations are available at https://data.broadinstitute.org/alkesgroup/LDSCORE/. The SHAP visualization of top 100 features for each model are at https://alkesgroup.broadinstitute.org/LDSCORE/DeepLearning/Dey_DeepBoost_Imperio/ExtDataFigures.

### Code Availability

The codes for generating DeepBoost and Imperio annotations are available in the Github repository https://github.com/kkdey/Imperio. This work primarily uses the S-LDSC software (https://github.com/bulik/ldsc). We used publicly available software for DeepSEA (https://github.com/FunctionLab/ExPecto), Basenji (https://github.com/calico/basenji), DeepBind (http://tools.genes.toronto.edu/deepbind/) and deltaSVM (https://www.beerlab.org/deltasvm/) to generate annotations for these respective models.

## Acknowledgments

We thank David Kelley, Steven Gazal, Alexander Gusev and Armin Schoech for helpful discussions. This research was funded by NIH grants U01 HG009379, R01 MH101244, R37 MH107649, R01 MH115676 and R01 MH109978. S.S.Kim was supported by NIH award F31HG010818. This research was conducted using the UK Biobank Resource under application 16549.

## Supplementary Tables

**Table S1.** List of all blood-related traits: List of 11 blood-related traits (6 autoimmune diseases and 5 blood cell traits) analyzed in this paper.

| Trait | Source | N |
| --- | --- | --- |
| Auto Immune Traits (Sure) | UKBiobank<sup>46</sup> | 459324 |
| Crohn's Disease | Jostins et al., 2012 Nature<sup>82</sup> | 20883 |
| Rheumatoid Arthritis | Okada et al., 2014 Nature<sup>83</sup> | 37681 |
| Ulcerative Colitis | Jostins et al., 2012 Nature<sup>82</sup> | 27432 |
| Lupus | Bentham et al., 2015<sup>84</sup> | 14267 |
| Celiac | Dubois et al., 2010<sup>85</sup> | 15283 |
| Platelet Count | UKBiobank<sup>46</sup> | 444382 |
| Red Blood Cell Count | UKBiobank<sup>46</sup> | 445174 |
| Red Blood Cell Distribution Width | UKBiobank<sup>46</sup> | 442700 |
| Eosinophil Count | UKBiobank<sup>46</sup> | 439938 |
| White Blood Cell Count | UKBiobank<sup>46</sup> | 444502 |

**Table S2.**
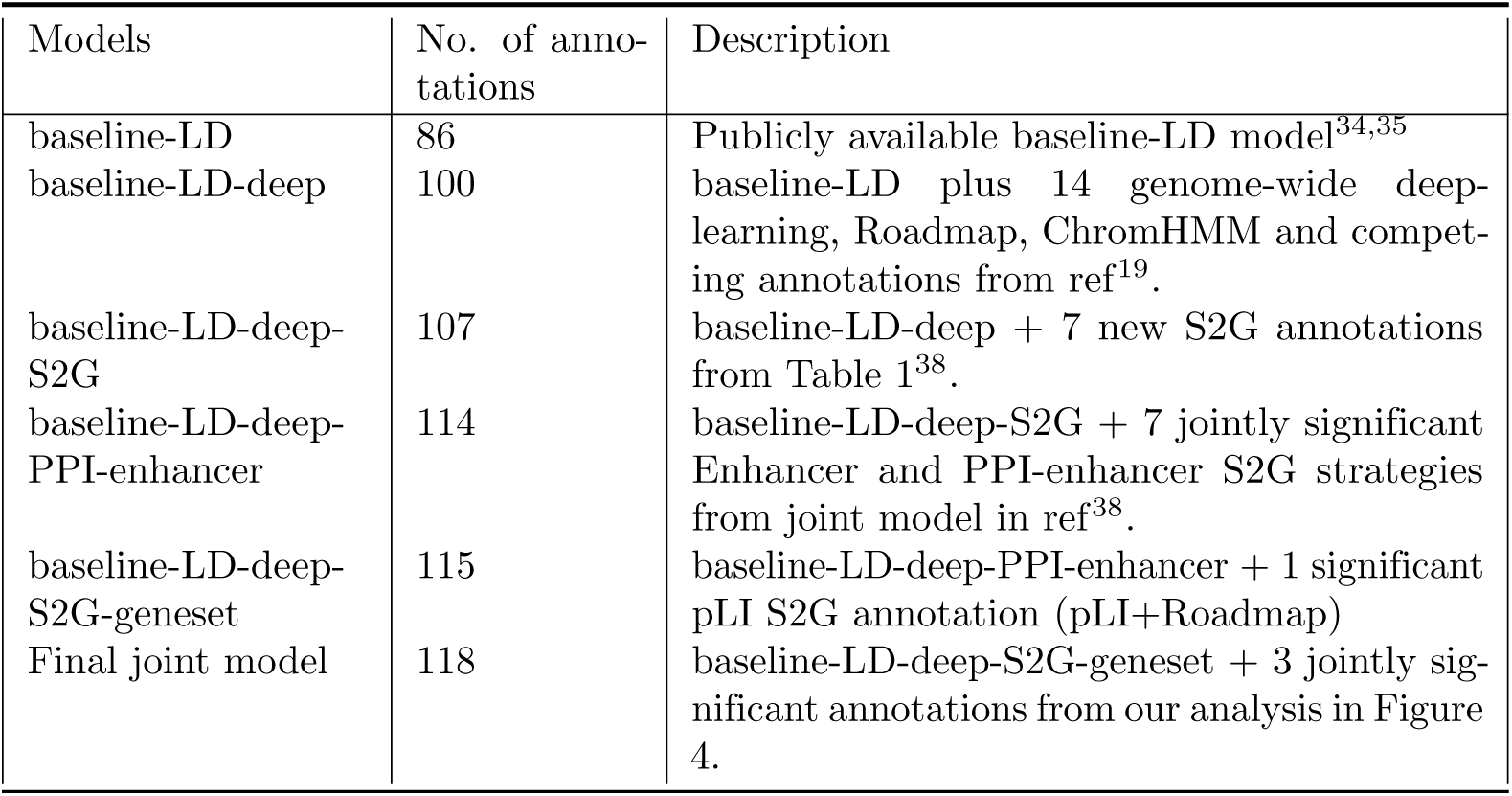
List of baseline models used in this paper: We report the 6 baseline models or joint models discussed in this paper, along with number of annotations and a brief description.

| Models | No. of annotations | Description |
| --- | --- | --- |
| baseline-LD | 86 | Publicly available baseline-LD model<sup>34,35</sup> |
| baseline-LD-deep | 100 | baseline-LD plus 14 genome-wide deep-learning, Roadmap, ChromHMM and competing annotations from ref<sup>19</sup>. |
| baseline-LD-deep-S2G | 107 | baseline-LD-deep + 7 new S2G annotations from Table 1<sup>38</sup>. |
| baseline-LD-deep-PPI-enhancer | 114 | baseline-LD-deep-S2G + 7 jointly significant Enhancer and PPI-enhancer S2G strategies from joint model in ref<sup>38</sup>. |
| baseline-LD-deep-S2G-geneset | 115 | baseline-LD-deep-PPI-enhancer + 1 significant pLI S2G annotation (pLI+Roadmap) |
| Final joint model | 118 | baseline-LD-deep-S2G-geneset + 3 jointly significant annotations from our analysis in Figure 4. |

**Table S3.**
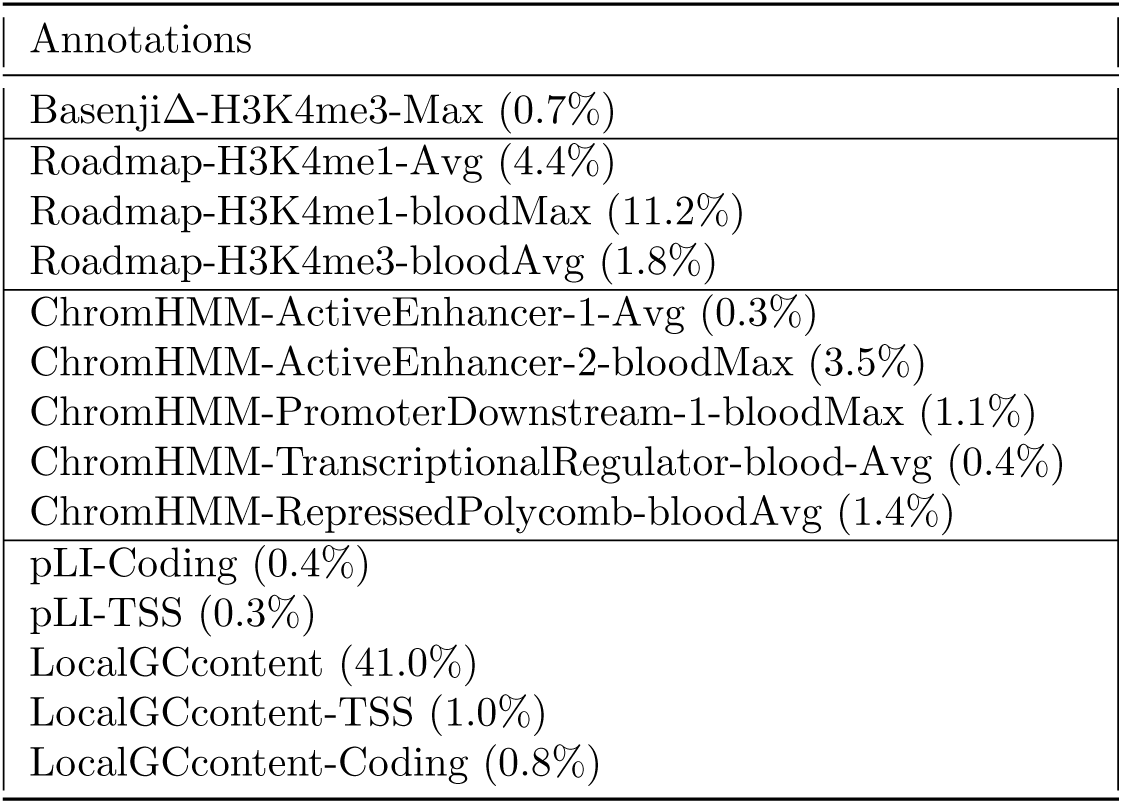
Additional annotations of baseline-LD-deep model: List of 14 jointly significant annotations from ref.^19^ added to the baseline-LD model to create the baseline- LD-deep model. They include 1 non-tissue-specific Basenji allelic-effect annotation, 3 Roadmap annotations, 5 ChromHMM annotations and 5 other annotations.

| Annotations |
| --- |
| BasenjiΔ-H3K4me3-Max (0.7%) |
| Roadmap-H3K4me1-Avg (4.4%) |
| Roadmap-H3K4me1-bloodMax (11.2%) |
| Roadmap-H3K4me3-bloodAvg (1.8%) |
| ChromHMM-ActiveEnhancer-1-Avg (0.3%) |
| ChromHMM-ActiveEnhancer-2-bloodMax (3.5%) |
| ChromHMM-PromoterDownstream-1-bloodMax (1.1%) |
| ChromHMM-TranscriptionalRegulator-blood-Avg (0.4%) |
| ChromHMM-RepressedPolycomb-bloodAvg (1.4%) |
| pLI-Coding (0.4%) |
| pLI-TSS (0.3%) |
| LocalGCcontent (41.0%) |
| LocalGCcontent-TSS (1.0%) |
| LocalGCcontent-Coding (0.8%) |

**Table S4.** AUROC attained by DeepBoost. We report the AUROC for a gradient boosting model distinguishing fine-mapped SNPs from matched control SNPs using allelic-effect annotations from the DeepSEA, Basenji, DeepBind and deltaSVM models as features. We consider four sets of fine-mapped SNPs - 8,741 fine-mapped autoimmune disease SNPs^21^ (Farh), 4,312 fine-mapped inflammatory bowel disease SNPs^22^ (Huang), 1,429 functionally fine-mapped SNPs for 14 blood-related UK Biobank traits^23, 46^ (Weissbrod), or union of all 14,482 fine-mapped SNPs (Union).

| Model | Fine-mapped SNPs | AUROC |
| --- | --- | --- |
| DeepSEA | Farh | 0.62 |
|  | Huang | 0.57 |
|  | Weissbrod | 0.72 |
|  | Union | 0.64 |
| Basenji | Farh | 0.67 |
|  | Huang | 0.68 |
|  | Weissbrod | 0.73 |
|  | Union | 0.69 |
| DeepBind | Farh | 0.58 |
|  | Huang | 0.54 |
|  | Weissbrod | 0.61 |
|  | Union | 0.58 |
| deltaSVM | Farh | 0.58 |
|  | Huang | 0.56 |
|  | Weissbrod | 0.55 |
|  | Union | 0.56 |

**Table S5.** AUROC attained by logistic classification instead of XGBoost. We report the AUROC for a logistic regression model distinguishing fine-mapped SNPs from matched control SNPs using allelic-effect annotations from the DeepSEA, Basenji, DeepBind and deltaSVM models as features. We consider four sets of fine-mapped SNPs - 8,741 fine-mapped autoimmune disease SNPs^21^ (Farh), 4,312 fine-mapped inflammatory bowel disease SNPs^22^ (Huang), 1,429 functionally fine-mapped SNPs for 14 blood-related UK Biobank traits^23, 46^ (Weissbrod), or union of all 14,482 fine-mapped SNPs (Union).

| Model | Fine-mapped SNPs | AUROC |
| --- | --- | --- |
| DeepSEA | Farh | 0.60 |
|  | Huang | 0.56 |
|  | Weissbrod | 0.69 |
|  | Union | 0.62 |
| Basenji | Farh | 0.65 |
|  | Huang | 0.67 |
|  | Weissbrod | 0.69 |
|  | Union | 0.67 |
| DeepBind | Farh | 0.57 |
|  | Huang | 0.54 |
|  | Weissbrod | 0.60 |
|  | Union | 0.58 |
| deltaSVM | Farh | 0.56 |
|  | Huang | 0.55 |
|  | Weissbrod | 0.56 |
|  | Union | 0.56 |

**Table S6.** Top features underlying the logistic regression model. We report the top 5 features for the logistic regression model (instead of the gradient boosting model) fitted on 8,741 fine-mapped autoimmune disease SNPs^21^ (Farh) corresponding to allelic-effect annotations from four different models: DeepSEA, Basenji, DeepBind and deltaSVM.

| Model | Top features |
| --- | --- |
| DeepSEA | Jurkat-DNase; Fetal Muscle Trunk H3K4me1; iPS-18 H3K4me1; CD19 Primary cells DNase; CD34 Cultured cells H3K4me1 |
| Basenji | GSM701490-DNASE:large intestine; HISTONE:H3K4me3 CD4 primary cells; HISTONE:H3K4me1 HepG2; HISTONE:H4K20me1 CD14-positive monocyte female |
| DeepBind | HOXA13 TF; Irx3 TF mouse; TEAD1 TF; NHLH1 TF; Hoxa11 TF mouse |
| deltaSVM | DHS RM 287 300 proms; TF E3 704 ZGPAT HepG2 ; DHS RM 203 300 proms; TF E3 563 E2F1 K562; TF E3 488 NFE2L2 HepG2 |

**Table S7.** AUROC attained by DeepBoost using 27 blood cell types only. We report the AUROC for a gradient boosting model distinguishing fine-mapped SNPs from matched control SNPs using using blood-specific allelic-effect from the DeepSEA, Basenji and deltaSVM models as features. (DeepBind model was not considered, as its features are non-tissue-specific.) We consider four sets of fine-mapped SNPs - 8,741 fine-mapped autoimmune disease SNPs^21^ (Farh), 4,312 fine-mapped inflammatory bowel disease SNPs^22^ (Huang), 1,429 functionally fine-mapped SNPs for 14 blood-related UK Biobank traits^23, 46^ (Weissbrod), or union of all 14,482 fine-mapped SNPs (Union).

| Model | Fine-mapped SNPs | AUROC |
| --- | --- | --- |
| DeepSEA | Farh | 0.60 |
|  | Huang | 0.56 |
|  | Weissbrod | 0.67 |
|  | Union | 0.62 |
| Basenji | Farh | 0.64 |
|  | Huang | 0.65 |
|  | Weissbrod | 0.70 |
|  | Union | 0.67 |
| deltaSVM | Farh | 0.54 |
|  | Huang | 0.53 |
|  | Weissbrod | 0.55 |
|  | Union | 0.54 |

**Table S8.** S-LDSC results for published and boosted allelic-effect annotations: Standardized Effect sizes (*τ\**) and Enrichment (E) of 4 published allelic-effect annotations for 4 sequence-based models, DeepSEA, Basenji, DeepBind and deltaSVM (DeepSEAΔ- published, BasenjiΔ-published, DeepBindΔ-published, deltaSVMΔ-published) and 16 boosted allelic-effect annotations for the same 4 deep learning models and 4 sets of finemapped SNPs for blood-related traits - 8,741 fine-mapped autoimmune disease SNPs^21^ (DeepSEAΔ-boosted, BasenjiΔ-boosted, DeepBindΔ-boosted, deltaSVMΔ- boosted), 4,312 fine-mapped inflammatory bowel disease SNPs^22^ (DeepSEAΔ-boosted- Huang, BasenjiΔ-boosted-Huang, DeepBindΔ-boosted-Huang, deltaSVMΔ-boosted- Huang), 1,429 functionally fine-mapped SNPs for 14 blood-related UK Biobank traits^23^ (DeepSEAΔ-boosted-Weissbrod, BasenjiΔ-boosted-Weissbrod, DeepBindΔ- boosted-Weissbrod, deltaSVMΔ-boosted-Weissbrod), or union of these fine-mapped SNPs (DeepSEAΔ-boosted-Union, BasenjiΔ-boosted-Union, DeepBindΔ-boosted-Union, deltaSVMΔ-boosted-Union). Results are conditioned on 100 baseline-LD-deep annotations. Reports are meta-analyzed across 11 blood and autoimmune traits.

| | $\tau^*$ | se($\tau^*$) | p($\tau^*$) | $E$ | se($E$) | p($E$) |
| --- | --- | --- | --- | --- | --- | --- |
| DeepSEAD-published | -0.27 | 0.12 | 0.02 | 1.3 | 0.046 | 9.5e-05 |
| BasenjiD-published | -0.046 | 0.083 | 0.58 | 1.5 | 0.028 | 2.2e-08 |
| DeepBindD-published | -0.29 | 0.17 | 0.066 | 1.1 | 0.24 | 0.69 |
| deltaSVMΔ-published | 0.034 | 0.043 | 0.42 | 1 | 0.012 | 0.086 |
| DeepSEAD-boosted | -0.065 | 0.085 | 0.44 | 2 | 0.19 | 0.00052 |
| DeepSEAD-boosted-Huang | 0.14 | 0.071 | 0.049 | 3 | 0.18 | 2.1e-09 |
| DeepSEAD-boosted-Weissbrod | -0.16 | 0.1 | 0.12 | 2.6 | 0.2 | 5e-05 |
| DeepSEAD-boosted-Union | -0.13 | 0.088 | 0.06 | 2.6 | 0.2 | 0.0001 |
| BasenjiD-boosted | 0.21 | 0.07 | 0.0032 | 3.4 | 0.2 | 3.8e-09 |
| BasenjiD-boosted-Huang | 0.14 | 0.071 | 0.049 | 3 | 0.18 | 2.1e-09 |
| BasenjiD-boosted-Weissbrod | -0.094 | 0.092 | 0.3 | 2.6 | 0.19 | 6e-05 |
| BasenjiD-boosted-Union | 0.13 | 0.066 | 0.02 | 3.3 | 0.19 | 6.7e-07 |
| DeepBindD-boosted | -0.044 | 0.099 | 0.66 | 1.1 | 0.12 | 0.10 |
| DeepBindD-boosted-Huang | 0.18 | 0.12 | 0.11 | 1.5 | 0.18 | 0.03 |
| DeepBindD-boosted-Weissbrod | 0.11 | 0.11 | 0.33 | 1.8 | 0.26 | 0.012 |
| DeepBindD-boosted-Union | 0.12 | 0.22 | 0.32 | 1.6 | 0.16 | 0.02 |
| deltaSVMΔ-boosted | 0.011 | 0.085 | 0.9 | 1.2 | 0.2 | 0.28 |
| deltaSVMΔ-boosted-Huang | 0.026 | 0.095 | 0.79 | 1.1 | 0.23 | 0.23 |
| deltaSVMΔ-boosted-Weissbrod | 0.12 | 0.1 | 0.25 | 1.7 | 0.24 | 0.12 |
| deltaSVMΔ-boosted-Union | 0.10 | 0.14 | 0.23 | 1.3 | 0.20 | 0.10 |

**Table S9.**
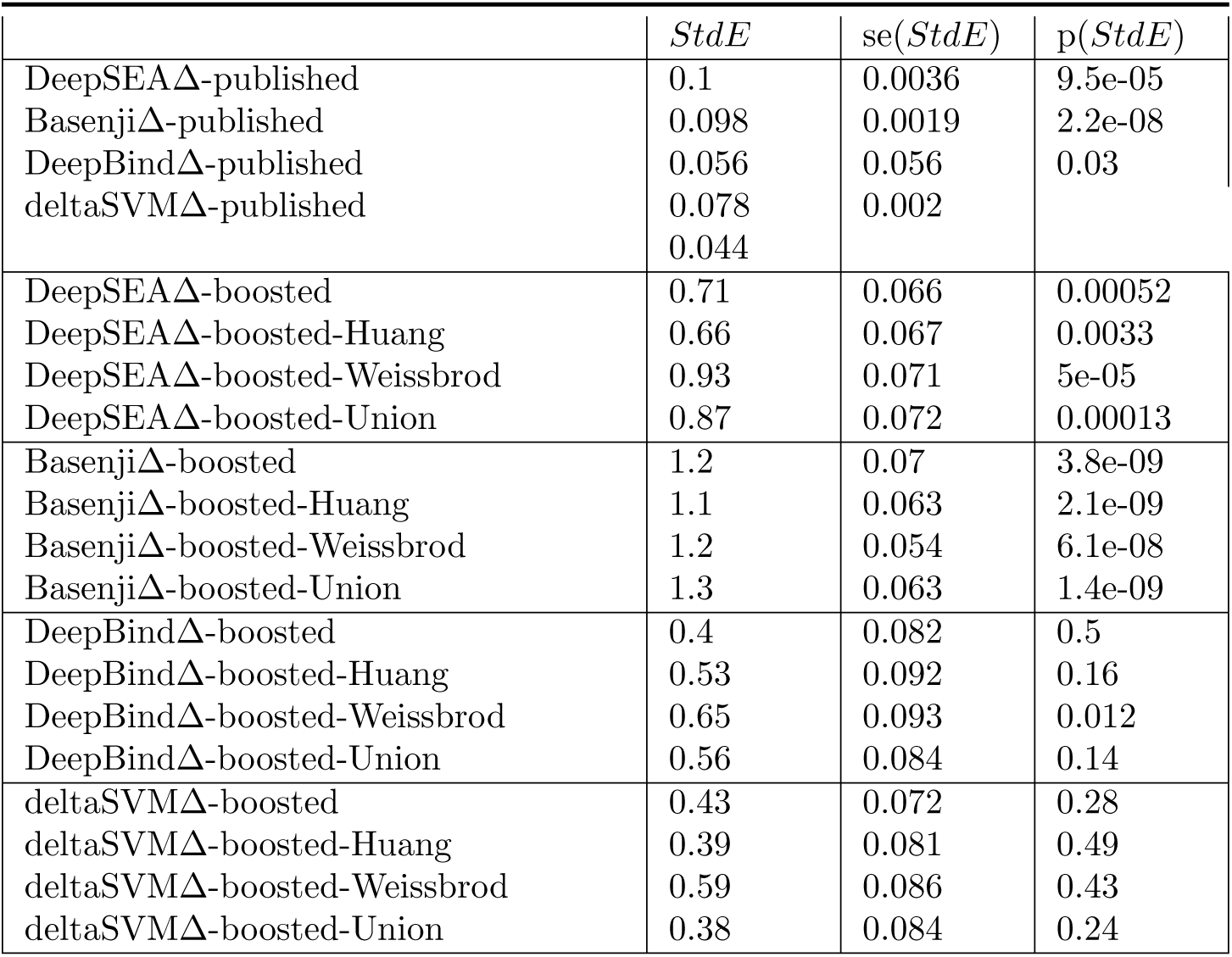
Standardized enrichment of SNP annotations for published and boosted allelic-effect annotations: Standardized enrichment of 4 published allelic-effect annotations for 4 sequence-based models, DeepSEA, Basenji, DeepBind and deltaSVM (DeepSEAΔ-published, BasenjiΔ-published, DeepBindΔ-published, deltaSVMΔ-published) and 16 boosted allelic-effect deep-learning annotations for the same 4 deep learning models and 4 sets of finemapped SNPs for blood-related traits - 8,741 fine-mapped autoimmune disease SNPs^21^ (DeepSEAΔ-boosted, BasenjiΔ-boosted, DeepBindΔ-boosted, deltaSVMΔ-boosted), 4,312 fine-mapped inflammatory bowel disease SNPs^22^ (DeepSEAΔ-boosted-Huang, BasenjiΔ-boosted-Huang, DeepBindΔ- boosted-Huang, deltaSVMΔ-boosted-Huang), 1,429 functionally fine-mapped SNPs for 14 blood-related UK Biobank traits^23^ (DeepSEAΔ-boosted-Weissbrod, BasenjiΔ- boosted-Weissbrod, DeepBindΔ-boosted-Weissbrod, deltaSVMΔ-boosted-Weissbrod), or union of these fine-mapped SNPs (DeepSEAΔ-boosted-Union, BasenjiΔ-boosted-Union, DeepBindΔ-boosted-Union, deltaSVMΔ-boosted-Union). Results are conditioned on 100 baseline-LD-deep annotations. Reports are meta-analyzed across 11 blood and autoimmune traits.

**Table S10.** Additional annotations of baseline-LD-deep-S2G model: List of 7 annotations corresponding to 7 S2G strategies linked to all genes from ref.^38^ added to the baseline-LD model to create the baseline-LD-deep-S2G model.

| Annotations |
| --- |
| ABC (1.4%) |
| ATAC (1.6%) |
| eQTL (2.4%) |
| Roadmap (3.2%) |
| PC-HiC (27%) |
| 5kb (53%) |
| 100kb (81%) |

**Table S11.** S-LDSC results for published-restricted and boosted-restricted allelic-effect annotations restricted using S2G strategies, conditional on the baseline-LD-deep-S2G model annotations: Standardized Effect sizes (*τ\**) and Enrichment (E) of SNP annotations corresponding to each of DeepSEAΔ-published, BasenjiΔ-published, DeepBindΔ-published, deltaSVMΔ-published, DeepSEAΔ-boosted, BasenjiΔ-boosted, DeepBindΔ-boosted and deltaSVMΔ-boosted annotations restricted using 10 S2G strategies conditional on 107 baseline-LD-deep-S2G annotations (100 baseline-LD-deep and 7 additional annotations from Table S10). Reports are meta-analyzed across 11 blood-related traits.

| DeepSEAΔ-published | | | | | | |
| --- | --- | --- | --- | --- | --- | --- |
| | $\tau^*$ | se($\tau^*$) | p($\tau^*$) | $E$ | se($E$) | p($E$) |
| ABC (0.29%) | 1.1 | 0.42 | 0.0092 | 10 | 1.2 | 3.6e-06 |
| TSS (0.40%) | -0.24 | 0.38 | 0.52 | 11 | 1 | 6.3e-06 |
| Coding (0.27%) | 0.35 | 0.28 | 0.21 | 8.7 | 1 | 7.5e-05 |
| ATAC (0.26%) | 0.75 | 0.22 | 0.00089 | 9.1 | 1.1 | 6e-07 |
| eQTL (0.37%) | 0.22 | 0.27 | 0.43 | 5.2 | 0.72 | 0.00028 |
| Roadmap (0.56%) | 0.34 | 0.37 | 0.36 | 8.3 | 1.2 | 3.4e-07 |
| Promoter (0.79%) | -0.06 | 0.23 | 0.8 | 5.3 | 0.55 | 8.1e-05 |
| PC-HiC (3.94%) | -0.28 | 0.18 | 0.12 | 2.7 | 0.18 | 6.1e-08 |
| 5kb (7.15%) | -0.3 | 0.12 | 0.014 | 1.8 | 0.067 | 1.8e-06 |
| 100kb (11.0%) | -0.27 | 0.1 | 0.0064 | 1.5 | 0.051 | 1.4e-05 |
| BasenjiΔ-published | | | | | | |
| | $\tau^*$ | se($\tau^*$) | p($\tau^*$) | $E$ | se($E$) | p($E$) |
| ABC (0.32%) | 1.0 | 0.34 | 0.002 | 9.2 | 0.94 | 7e-06 |
| TSS (0.44%) | 0.036 | 0.46 | 0.94 | 12 | 1.4 | 1.2e-05 |
| Coding (0.33%) | 0.81 | 0.33 | 0.014 | 9.2 | 1.1 | 8.6e-05 |
| ATAC (0.31%) | 0.58 | 0.29 | 0.045 | 8.4 | 1.2 | 1.5e-06 |
| eQTL (0.44%) | 0.27 | 0.24 | 0.26 | 5.2 | 0.5 | 1.5e-05 |
| Roadmap (0.67%) | 0.85 | 0.56 | 0.13 | 8.4 | 1.1 | 1.6e-08 |
| Promoter (0.92%) | 0.2 | 0.34 | 0.55 | 5.5 | 0.55 | 2.3e-05 |
| PC-HiC (4.80%) | 0.091 | 0.25 | 0.72 | 2.8 | 0.15 | 2.8e-10 |
| 5kb (8.58%) | -0.25 | 0.13 | 0.05 | 1.9 | 0.047 | 2.4e-08 |
| 100kb (13.1%) | -0.091 | 0.1 | 0.37 | 1.6 | 0.032 | 1.6e-08 |
| DeepBindΔ-published | | | | | | |
| | $\tau^*$ | se($\tau^*$) | p($\tau^*$) | $E$ | se($E$) | p($E$) |
| ABC (0.30%) | -0.4 | 0.69 | 0.56 | 6.7 | 0.79 | 0.00012 |
| TSS (0.34%) | -0.41 | 0.19 | 0.034 | 10 | 0.91 | 1.4e-05 |
| Coding (0.34%) | -0.95 | 0.96 | 0.32 | 6.2 | 0.7 | 0.00045 |
| ATAC (0.35%) | 0.5 | 0.63 | 0.42 | 5.4 | 0.6 | 5e-06 |
| eQTL (0.28%) | -0.21 | 0.65 | 0.75 | 3.8 | 0.42 | 0.00073 |
| Roadmap (0.73%) | -1.4 | 0.77 | 0.072 | 6.3 | 0.66 | 2.4e-08 |
| Promoter (0.74%) | -1.1 | 1 | 0.27 | 3.7 | 0.47 | 0.00081 |
| PC-HiC (3.24%) | 0.021 | 0.52 | 0.97 | 2 | 0.066 | 8.4e-08 |
| 5kb (9.66%) | 0.15 | 0.33 | 0.66 | 1.4 | 0.034 | 8.8e-05 |
| 100kb (14.2%) | -0.065 | 0.19 | 0.74 | 1.2 | 0.022 | 0.00097 |
| deltaSVMΔ-published | | | | | | |
| | $\tau^*$ | se($\tau^*$) | p($\tau^*$) | $E$ | se($E$) | p($E$) |
| ABC (0.30%) | -0.62 | 0.44 | 0.16 | 6.3 | 0.82 | 0.00024 |
| TSS (0.30%) | 0.35 | 0.5 | 0.48 | 11 | 0.96 | 1.3e-05 |
| Coding (0.27%) | 0.013 | 0.37 | 0.97 | 6.3 | 0.73 | 0.00042 |
| ATAC (0.30%) | -1.3 | 0.64 | 0.042 | 4.4 | 0.57 | 0.00023 |
| eQTL (0.24%) | 0.54 | 0.42 | 0.2 | 4.6 | 0.44 | 0.00026 |
| Roadmap (0.69%) | 0.16 | 0.42 | 0.69 | 6.8 | 0.8 | 5e-08 |
| Promoter (0.73%) | 0.22 | 0.45 | 0.63 | 4 | 0.6 | 0.00086 |
| PC-HiC (3.06%) | 0.054 | 0.31 | 0.86 | 2 | 0.098 | 3.9e-06 |
| 5kb (8.4%) | 0.16 | 0.22 | 0.46 | 1.4 | 0.065 | 0.00065 |
| 100kb (12.5%) | 0.06 | 0.2 | 0.76 | 1.2 | 0.046 | 0.021 |
| DeepSEAΔ-boosted | | | | | | |
| | $\tau^*$ | se($\tau^*$) | p($\tau^*$) | $E$ | se($E$) | p($E$) |
| ABC (0.50%) | 0.29 | 0.18 | 0.1 | 10 | 1.8 | 0.0007 |
| TSS (0.75%) | 0.45 | 0.22 | 0.037 | 14 | 1.6 | 6.3e-05 |
| Coding (0.47%) | 0.44 | 0.18 | 0.013 | 12 | 1.9 | 0.00032 |
| ATAC (0.46%) | 0.18 | 0.14 | 0.2 | 8.6 | 1.6 | 0.00087 |
| eQTL (0.59%) | 0.59 | 0.14 | 1.5e-05 | 11 | 1.4 | 0.00022 |
| Roadmap (1.0%) | 0.2 | 0.18 | 0.28 | 8.3 | 1.4 | 0.00038 |
| Promoter (1.33%) | 0.29 | 0.2 | 0.16 | 6.3 | 1.1 | 0.001 |
| PC-HiC (5.89%) | -0.1 | 0.13 | 0.42 | 3.1 | 0.49 | 0.00038 |
| 5kb (9.89%) | -0.12 | 0.088 | 0.16 | 2.3 | 0.22 | 0.00052 |
| 100kb (13.7%) | -0.079 | 0.087 | 0.37 | 2.1 | 0.2 | 0.00062 |

BasenjiΔ-boosted
| | $\tau^*$ | se($\tau^*$) | p($\tau^*$) | $E$ | se($E$) | p($E$) |
| --- | --- | --- | --- | --- | --- | --- |
| ABC (0.63%) | 1.1 | 0.28 | 0.00016 | 15 | 2.3 | 8.8e-06 |
| TSS (0.88%) | 1.3 | 0.34 | 0.00011 | 17 | 1.9 | 1.8e-06 |
| Coding (0.59%) | 0.55 | 0.21 | 0.01 | 11 | 1.6 | 0.00015 |
| ATAC (0.60%) | 0.35 | 0.19 | 0.059 | 11 | 2.1 | 2.4e-05 |
| eQTL (0.79%) | 0.21 | 0.1 | 0.036 | 7.3 | 0.86 | 0.00015 |
| Roadmap (1.54%) | 0.76 | 0.26 | 0.0036 | 11 | 1.6 | 1.1e-07 |
| Promoter (1.63%) | 0.19 | 0.14 | 0.18 | 6.6 | 0.86 | 4.4e-05 |
| PC-HiC (7.67%) | 0.22 | 0.12 | 0.082 | 4.4 | 0.42 | 2.4e-09 |
| 5kb (11.2%) | 0.01 | 0.07 | 0.88 | 3.1 | 0.15 | 6.8e-08 |
| 100kb (14.5%) | 0.15 | 0.071 | 0.034 | 3.3 | 0.19 | 1.4e-08 |

DeepBindΔ-boosted
| | $\tau^*$ | se($\tau^*$) | p($\tau^*$) | $E$ | se($E$) | p($E$) |
| --- | --- | --- | --- | --- | --- | --- |
| ABC (0.27%) | 0.1 | 0.21 | 0.63 | 9.1 | 3.5 | 0.088 |
| TSS (0.37%) | -0.071 | 0.17 | 0.68 | 12 | 2.8 | 0.0042 |
| Coding (0.34%) | -0.21 | 0.14 | 0.13 | 4.5 | 1.8 | 0.98 |
| ATAC (0.31%) | 0.42 | 0.14 | 0.0027 | 12 | 2.1 | 8.9e-05 |
| eQTL (0.43%) | -0.11 | 0.16 | 0.49 | 3.3 | 2.1 | 0.74 |
| Roadmap (0.57%) | -0.05 | 0.23 | 0.83 | 6 | 2.3 | 0.053 |
| Promoter (0.84%) | -0.12 | 0.23 | 0.61 | 4 | 2 | 0.37 |
| PC-HiC (4.46%) | 0.024 | 0.15 | 0.87 | 2.5 | 0.6 | 0.029 |
| 5kb (8.24%) | -0.16 | 0.1 | 0.12 | 1.2 | 0.32 | 0.52 |
| 100kb (12.9%) | -0.068 | 0.1 | 0.51 | 1.2 | 0.25 | 0.49 |

deltaSVMΔ-boosted
| | $\tau^*$ | se($\tau^*$) | p($\tau^*$) | $E$ | se($E$) | p($E$) |
| --- | --- | --- | --- | --- | --- | --- |
| ABC (0.25%) | 0.51 | 0.17 | 0.0027 | 15 | 2.1 | 0.0001 |
| TSS (0.33%) | 0.27 | 0.25 | 0.28 | 16 | 3.8 | 0.0002 |
| Coding (0.31%) | 0.34 | 0.23 | 0.15 | 11 | 2.5 | 7.8e-05 |
| ATAC (0.28%) | 0.13 | 0.14 | 0.36 | 6.6 | 2.3 | 0.25 |
| eQTL (0.41%) | 0.22 | 0.17 | 0.21 | 6.5 | 2.3 | 0.082 |
| Roadmap (0.53%) | 0.11 | 0.18 | 0.56 | 6.3 | 2.1 | 0.039 |
| Promoter (0.77%) | -0.014 | 0.16 | 0.93 | 4.5 | 1.5 | 0.06 |
| PC-HiC (4.3%) | 0.097 | 0.15 | 0.52 | 2.7 | 0.59 | 0.024 |
| 5kb (8.2%) | -0.041 | 0.14 | 0.77 | 1.5 | 0.43 | 0.12 |
| 100kb (12.8%) | -0.034 | 0.11 | 0.77 | 1.3 | 0.27 | 0.12 |

**Table S12.** Standardized enrichment of published-restricted and boosted-restricted allelic-effect annotations restricted using S2G strategies, conditional on the baseline-LD-deep-S2G model annotations: Standardized enrichment of restricted SNP annotations corresponding to each of DeepSEAΔ-published, BasenjiΔ-published, DeepBindΔ-published, deltaSVMΔ-published, DeepSEAΔ-boosted, BasenjiΔ-boosted, DeepBindΔ-boosted and deltaSVMΔ-boosted annotations restricted using 10 S2G strategies conditional on 107 baseline-LD-deep-S2G annotations (100 baseline-LD-deep and 7 additional annotations from Table S10). Reports are meta-analyzed across 11 blood-related traits.

| DeepSEAΔ-published | | | |
| --- | --- | --- | --- |
|  | *StdE* | *se(StdE)* | *p(StdE)* |
| ABC (0.29%) | 0.28 | 0.034 | 3.6e-06 |
| TSS (0.40%) | 0.38 | 0.036 | 6.3e-06 |
| Coding (0.27%) | 0.21 | 0.025 | 7.5e-05 |
| ATAC (0.26%) | 0.22 | 0.027 | 6e-07 |
| eQTL (0.37%) | 0.14 | 0.02 | 0.00028 |
| Roadmap (0.56%) | 0.29 | 0.042 | 3.4e-07 |
| Promoter (0.79%) | 0.23 | 0.024 | 8.1e-05 |
| PC-HiC (3.94%) | 0.21 | 0.014 | 6.1e-08 |
| 5kb (7.15%) | 0.16 | 0.0061 | 1.8e-06 |
| 100kb (11.0%) | 0.13 | 0.0044 | 1.4e-05 |
| BasenjiΔ-published | | | |
|  | *StdE* | *se(StdE)* | *p(StdE)* |
| ABC (0.32%) | 0.27 | 0.028 | 7e-06 |
| TSS (0.44%) | 0.44 | 0.051 | 1.2e-05 |
| Coding (0.33%) | 0.26 | 0.03 | 8.6e-05 |
| ATAC (0.31%) | 0.23 | 0.033 | 1.5e-06 |
| eQTL (0.44%) | 0.16 | 0.016 | 1.5e-05 |
| Roadmap (0.67%) | 0.34 | 0.044 | 1.6e-08 |
| Promoter (0.92%) | 0.26 | 0.026 | 2.3e-05 |
| PC-HiC (4.80%) | 0.25 | 0.013 | 2.8e-10 |
| 5kb (8.58%) | 0.18 | 0.0045 | 2.4e-08 |
| 100kb (13.1%) | 0.14 | 0.0027 | 1.6e-08 |
| DeepBindΔ-published | | | |
|  | *StdE* | *se(StdE)* | *p(StdE)* |
| ABC (0.30%) | 0.47 | 0.055 | 0.00012 |
| TSS (0.34%) | 0.78 | 0.069 | 1.4e-05 |
| Coding (0.34%) | 0.47 | 0.053 | 0.00045 |
| ATAC (0.35%) | 0.41 | 0.046 | 5e-06 |
| eQTL (0.28%) | 0.36 | 0.04 | 0.00073 |
| Roadmap (0.73%) | 0.67 | 0.07 | 2.4e-08 |
| Promoter (0.74%) | 0.45 | 0.058 | 0.00081 |
| PC-HiC (3.24%) | 0.56 | 0.018 | 8.4e-08 |
| 5kb (9.66%) | 0.44 | 0.011 | 8.8e-05 |
| 100kb (14.2%) | 0.28 | 0.0053 | 0.00097 |
| deltaSVMΔ-published | | | |
|  | *StdE* | *se(StdE)* | *p(StdE)* |
| ABC (0.30%) | 0.29 | 0.037 | 0.00024 |
| TSS (0.30%) | 0.56 | 0.048 | 1.3e-05 |
| Coding (0.27%) | 0.3 | 0.035 | 0.00042 |
| ATAC (0.30%) | 0.22 | 0.029 | 0.00023 |
| eQTL (0.24%) | 0.29 | 0.028 | 0.00026 |
| Roadmap (0.69%) | 0.48 | 0.056 | 5e-08 |
| Promoter (0.73%) | 0.33 | 0.049 | 0.00086 |
| PC-HiC (3.06%) | 0.37 | 0.018 | 3.9e-06 |
| 5kb (8.4%) | 0.3 | 0.014 | 0.00065 |
| 100kb (12.5%) | 0.22 | 0.0084 | 0.021 |
| DeepSEAΔ-boosted | | | |
|  | *StdE* | *se(StdE)* | *p(StdE)* |
| ABC (0.29%) | 0.74 | 0.13 | 0.0007 |
| TSS (0.40%) | 1.2 | 0.14 | 6.3e-05 |
| Coding (0.27%) | 0.8 | 0.13 | 0.00032 |
| ATAC (0.26%) | 0.58 | 0.11 | 0.00087 |
| eQTL (0.37%) | 0.86 | 0.11 | 0.00022 |
| Roadmap (0.56%) | 0.83 | 0.14 | 0.00038 |
| Promoter (0.79%) | 0.72 | 0.12 | 0.001 |
| PC-HiC (3.94%) | 0.74 | 0.11 | 0.00038 |
| 5kb (7.15%) | 0.68 | 0.066 | 0.00052 |
| 100kb (11.0%) | 0.71 | 0.067 | 0.00062 |

BasenjiΔ-boosted
|  | *StdE* | *se(StdE)* | *p(StdE)* |
| --- | --- | --- | --- |
| ABC (0.63%) | 1.2 | 0.18 | 8.8e-06 |
| TSS (0.88%) | 1.5 | 0.18 | 1.8e-06 |
| Coding (0.59%) | 0.84 | 0.12 | 0.00015 |
| ATAC (0.60%) | 0.86 | 0.16 | 2.4e-05 |
| eQTL (0.79%) | 0.65 | 0.076 | 0.00015 |
| Roadmap (1.54%) | 1.4 | 0.19 | 1.1e-07 |
| Promoter (1.63%) | 0.84 | 0.11 | 4.4e-05 |
| PC-HiC (7.67%) | 1.2 | 0.11 | 2.4e-09 |
| 5kb (11.2%) | 0.99 | 0.048 | 6.8e-088 |
| 100kb (14.5%) | 1.2 | 0.067 | 1.4e-08 |

DeepBindΔ-boosted
|  | *StdE* | *se(StdE)* | *p(StdE)* |
| --- | --- | --- | --- |
| ABC (0.27%) | 0.48 | 0.18 | 0.088 |
| TSS (0.37%) | 0.71 | 0.17 | 0.0042 |
| Coding (0.34%) | 0.26 | 0.11 | 0.98 |
| ATAC (0.31%) | 0.66 | 0.12 | 0.00071 |
| eQTL (0.43%) | 0.22 | 0.14 | 0.74 |
| Roadmap (0.57%) | 0.45 | 0.17 | 0.053 |
| Promoter (0.84%) | 0.37 | 0.18 | 0.37 |
| PC-HiC (4.46%) | 0.52 | 0.12 | 0.029 |
| 5kb (8.24%) | 0.33 | 0.088 | 0.52 |
| 100kb (12.9%) | 0.41 | 0.085 | 0.99 |

deltaSVMΔ-boosted
|  | *StdE* | *se(StdE)* | *p(StdE)* |
| --- | --- | --- | --- |
| ABC (0.25%) | 0.75 | 0.14 | 0.0011 |
| TSS (0.33%) | 0.92 | 0.24 | 0.0019 |
| Coding (0.31%) | 0.63 | 0.18 | 0.0052 |
| ATAC (0.28%) | 0.4 | 0.12 | 0.25 |
| eQTL (0.41%) | 0.48 | 0.15 | 0.082 |
| Roadmap (0.53%) | 0.53 | 0.15 | 0.039 |
| Promoter (0.77%) | 0.4 | 0.13 | 0.06 |
| PC-HiC (4.3%) | 0.56 | 0.12 | 0.024 |
| 5kb (8.2%) | 0.42 | 0.12 | 0.12 |
| 100kb (12.8%) | 0.42 | 0.092 | 0.12 |

**Table S13.** S-LDSC results for published-restricted and boosted-restricted allelic-effect annotations restricted using S2G strategies, conditional on the baseline-LD-deep model annotations : Standardized Effect sizes (*τ\**) and Enrichment (E) of 80 restricted SNP annotations corresponding to DeepSEAΔ-published, BasenjiΔ-published, DeepBindΔ-published, deltaSVMΔ-published, DeepSEAΔ-boosted, BasenjiΔ-boosted, DeepBindΔ-boosted and deltaSVMΔ-boosted annotations restricted using 10 S2G strategies. Results are conditional on 100 baseline-LD-deep annotations. Reports are meta-analyzed across 11 blood-related traits.

| AllSNPs |  |  |  |  |  |  |
| --- | --- | --- | --- | --- | --- | --- |
| | $\tau^*$ | se($\tau^*$) | p($\tau^*$) | $E$ | se($E$) | p($E$) |
| ABC (1.3%) | 0.2 | 0.096 | 0.034 | 7.4 | 0.83 | 5.5e-06 |
| TSS (1.6%) | 0.49 | 0.19 | 0.012 | 12 | 0.81 | 4.3e-07 |
| Coding (1.6%) | 0.39 | 0.13 | 0.0026 | 6.9 | 0.63 | 1.3e-05 |
| ATAC (1.6%) | 0.4 | 0.11 | 0.00018 | 6.6 | 0.79 | 5.3e-08 |
| eQTL (2.4%) | 0.15 | 0.06 | 0.012 | 4.1 | 0.35 | 4.5e-06 |
| Roadmap (3.1%) | 0.42 | 0.12 | 0.00038 | 7 | 0.67 | 6e-10 |
| Promoter (4.2%) | -0.093 | 0.097 | 0.33 | 4.3 | 0.34 | 3.1e-05 |
| PC-HiC (27.3%) | 0.02 | 0.035 | 0.57 | 2.1 | 0.062 | 1.2e-10 |
| 5kb (52%) | 0.007 | 0.02 | 0.73 | 1.4 | 0.028 | 5.5e-08 |
| 100kb (81%) | 0.0091 | 0.0091 | 0.32 | 1.2 | 0.0067 | 7.4e-10 |
| DeepSEAΔ-published | | | | | | |
| | $\tau^*$ | se($\tau^*$) | p($\tau^*$) | $E$ | se($E$) | p($E$) |
| ABC (0.29%) | 0.32 | 0.1 | 0.0013 | 9.3 | 1.1 | 3.6e-06 |
| TSS (0.40%) | 0.35 | 0.2 | 0.086 | 13 | 0.95 | 4.5e-07 |
| Coding (0.27%) | 0.63 | 0.15 | 3.9e-05 | 9.8 | 0.99 | 2.3e-05 |
| ATAC (0.26%) | 0.58 | 0.14 | 3.7e-05 | 9.5 | 1.2 | 2.3e-08 |
| eQTL (0.37%) | 0.17 | 0.074 | 0.02 | 5.6 | 0.46 | 2.8e-06 |
| Roadmap (0.56%) | 0.55 | 0.15 | 0.00021 | 8.7 | 0.96 | 1.3e-09 |
| Promoter (0.79%) | -0.019 | 0.11 | 0.86 | 6.5 | 0.49 | 2e-06 |
| PC-HiC (3.94%) | 0.02 | 0.049 | 0.68 | 2.8 | 0.1 | 1.7e-10 |
| 5kb (7.15%) | -0.016 | 0.033 | 0.63 | 2 | 0.038 | 1.1e-08 |
| 100kb (11.0%) | -0.016 | 0.023 | 0.48 | 1.6 | 0.024 | 1.9e-09 |
| BasenjiΔ-published | | | | | | |
| | $\tau^*$ | se($\tau^*$) | p($\tau^*$) | $E$ | se($E$) | p($E$) |
| ABC (0.32%) | 0.31 | 0.1 | 0.0023 | 9.1 | 0.96 | 3.3e-06 |
| TSS (0.44%) | 0.5 | 0.22 | 0.022 | 13 | 1 | 6.6e-07 |
| Coding (0.33%) | 0.78 | 0.17 | 3.5e-06 | 10 | 1 | 1.8e-05 |
| ATAC (0.31%) | 0.52 | 0.13 | 0.0001 | 8.8 | 1.1 | 2.5e-08 |
| eQTL (0.44%) | 0.17 | 0.071 | 0.017 | 5.4 | 0.41 | 1.1e-06 |
| Roadmap (0.67%) | 0.53 | 0.14 | 0.00026 | 8.3 | 0.86 | 3.7e-10 |
| Promoter (0.92%) | 0.019 | 0.11 | 0.86 | 6.4 | 0.47 | 1.4e-06 |
| PC-HiC (4.80%) | 0.034 | 0.052 | 0.51 | 2.8 | 0.093 | 6.5e-11 |
| 5kb (8.58%) | -0.007 | 0.031 | 0.82 | 2 | 0.033 | 5.2e-09 |
| 100kb (13.1%) | -0.001 | 0.021 | 0.96 | 1.6 | 0.022 | 8.1e-10 |
| DeepBindΔ-published | | | | | | |
| | $\tau^*$ | se($\tau^*$) | p($\tau^*$) | $E$ | se($E$) | p($E$) |
| ABC (0.30%) | 0.21 | 0.097 | 0.027 | 7.4 | 0.82 | 1.2e-05 |
| TSS (0.34%) | 0.49 | 0.23 | 0.031 | 12 | 0.82 | 5.9e-07 |
| Coding (0.34%) | 0.26 | 0.46 | 0.57 | 7 | 0.66 | 4e-05 |
| ATAC (0.35%) | 0.4 | 0.11 | 0.00019 | 6.7 | 0.81 | 7.4e-08 |
| eQTL (0.28%) | 0.15 | 0.061 | 0.013 | 4.2 | 0.36 | 6.7e-06 |
| Roadmap (0.73%) | 0.42 | 0.12 | 0.00045 | 7.1 | 0.69 | 9.1e-10 |
| Promoter (0.74%) | 0.53 | 0.15 | 0.00034 | 4.4 | 0.34 | 2.8e-05 |
| PC-HiC (3.24%) | 0.019 | 0.036 | 0.61 | 2.1 | 0.065 | 1.5e-10 |
| 5kb (9.66%) | 0.0044 | 0.021 | 0.83 | 1.4 | 0.029 | 1.4e-07 |
| 100kb (14.2%) | 0.0027 | 0.011 | 0.81 | 1.2 | 0.0069 | 7e-09 |
| deltaSVMΔ-published | | | | | | |
| | $\tau^*$ | se($\tau^*$) | p($\tau^*$) | $E$ | se($E$) | p($E$) |
| ABC (0.30%) | 0.21 | 0.092 | 0.023 | 7.5 | 0.85 | 1.7e-05 |
| TSS (0.30%) | 0.58 | 0.19 | 0.0022 | 12 | 0.87 | 5.4e-07 |
| Coding (0.27%) | 0.31 | 0.26 | 0.25 | 7.4 | 0.7 | 4.9e-05 |
| ATAC (0.30%) | 0.4 | 0.11 | 0.00022 | 6.8 | 0.84 | 1.3e-07 |
| eQTL (0.24%) | 0.17 | 0.064 | 0.0089 | 4.3 | 0.4 | 1.3e-05 |
| Roadmap (0.69%) | 0.46 | 0.13 | 0.0003 | 7.4 | 0.75 | 1.4e-09 |
| Promoter (0.73%) | 0.59 | 0.16 | 0.00017 | 4.7 | 0.36 | 1.6e-05 |
| PC-HiC (3.06%) | 0.021 | 0.039 | 0.6 | 2.1 | 0.072 | 2.1e-10 |
| 5kb (8.4%) | 0.0077 | 0.024 | 0.74 | 1.4 | 0.036 | 4.3e-07 |
| 100kb (12.5%) | 0.0058 | 0.014 | 0.68 | 1.2 | 0.012 | 1.3e-07 |
| DeepSEAΔ-boosted | | | | | | |
| | $\tau^*$ | se($\tau^*$) | p($\tau^*$) | $E$ | se($E$) | p($E$) |
| ABC (0.50%) | 0.36 | 0.12 | 0.003 | 11 | 1.6 | 0.00016 |
| TSS (0.75%) | 0.58 | 0.18 | 0.001 | 16 | 1.5 | 6e-06 |
| Coding (0.47%) | 0.69 | 0.17 | 3.1e-05 | 13 | 1.9 | 0.00013 |
| ATAC (0.46%) | 0.65 | 0.18 | 0.00025 | 14 | 2.5 | 3.1e-06 |
| eQTL (0.59%) | 0.39 | 0.1 | 0.00014 | 9.7 | 1.2 | 7e-05 |
| Roadmap (1.0%) | 0.66 | 0.18 | 0.00019 | 12 | 1.8 | 1e-06 |
| Promoter (1.33%) | 0.13 | 0.12 | 0.27 | 8.5 | 1.1 | 1.4e-05 |
| PC-HiC (5.89%) | 0.053 | 0.11 | 0.62 | 3.6 | 0.41 | 1.2e-06 |
| 5kb (9.89%) | -0.056 | 0.077 | 0.46 | 2.5 | 0.2 | 2.9e-05 |
| 100kb (13.7%) | -0.045 | 0.072 | 0.53 | 2.2 | 0.16 | 0.00011 |
| BasenjiΔ-boosted | | | | | | |
| | $\tau^*$ | se($\tau^*$) | p($\tau^*$) | $E$ | se($E$) | p($E$) |
| ABC (0.63%) | 0.5 | 0.12 | 5.5e-05 | 13 | 1.8 | 6.8e-06 |
| TSS (0.88%) | 1.2 | 0.27 | 9.1e-06 | 18 | 1.8 | 2.8e-07 |
| Coding (0.59%) | 0.82 | 0.21 | 5.9e-05 | 12 | 1.6 | 2.9e-05 |
| ATAC (0.60%) | 0.66 | 0.18 | 0.00017 | 13 | 2.2 | 2.7e-07 |
| eQTL (0.79%) | 0.26 | 0.087 | 0.0027 | 7.9 | 0.87 | 2.3e-05 |
| Roadmap (1.54%) | 0.71 | 0.18 | 4.6e-05 | 11 | 1.4 | 3.5e-09 |
| Promoter (1.63%) | 0.23 | 0.12 | 0.047 | 8.1 | 0.83 | 1.7e-06 |
| PC-HiC (7.67%) | 0.2 | 0.11 | 0.07 | 4.4 | 0.39 | 5.4e-10 |
| 5kb (11.2%) | 0.07 | 0.068 | 0.3 | 3.3 | 0.15 | 2.1e-08 |
| 100kb (14.5%) | 0.17 | 0.069 | 0.013 | 3.4 | 0.2 | 6.4e-09 |
| DeepBindΔ-boosted | | | | | | |
| | $\tau^*$ | se($\tau^*$) | p($\tau^*$) | $E$ | se($E$) | p($E$) |
| ABC (0.27%) | 0.28 | 0.16 | 0.072 | 11 | 3 | 0.0057 |
| TSS (0.37%) | 0.17 | 0.19 | 0.36 | 15 | 2.7 | 0.00041 |
| Coding (0.34%) | -0.18 | 0.13 | 0.18 | 5.7 | 1.8 | 0.088 |
| ATAC (0.31%) | 0.77 | 0.18 | 2.5e-05 | 17 | 3.1 | 5e-06 |
| eQTL (0.43%) | 0.27 | 0.12 | 0.024 | 7.9 | 1.8 | 0.0043 |
| Roadmap (0.57%) | 0.48 | 0.14 | 0.00078 | 12 | 2.1 | 6.1e-05 |
| Promoter (0.84%) | 0.27 | 0.21 | 0.18 | 7.5 | 1.9 | 0.0062 |
| PC-HiC (4.46%) | 0.043 | 0.081 | 0.59 | 2.7 | 0.38 | 0.00065 |
| 5kb (8.24%) | -0.017 | 0.07 | 0.81 | 1.7 | 0.22 | 0.019 |
| 100kb (12.9%) | -0.0068 | 0.056 | 0.9 | 1.4 | 0.14 | 0.049 |
| deltaSVMΔ-boosted | | | | | | |
| | $\tau^*$ | se($\tau^*$) | p($\tau^*$) | $E$ | se($E$) | p($E$) |
| ABC (0.25%) | 0.46 | 0.13 | 0.00033 | 15 | 2.5 | 0.00057 |
| TSS (0.33%) | 0.41 | 0.24 | 0.083 | 19 | 4 | 0.00018 |
| Coding (0.31%) | 0.39 | 0.24 | 0.099 | 13 | 3.5 | 0.0015 |
| ATAC (0.28%) | 0.62 | 0.16 | 0.00017 | 15 | 3 | 6.3e-05 |
| eQTL (0.41%) | 0.34 | 0.14 | 0.013 | 8.9 | 2.1 | 0.0033 |
| Roadmap (0.53%) | 0.58 | 0.18 | 0.0013 | 13 | 2.4 | 0.00012 |
| Promoter (0.77%) | 0.33 | 0.16 | 0.04 | 8 | 1.6 | 0.00063 |
| PC-HiC (4.3%) | 0.034 | 0.071 | 0.63 | 2.5 | 0.32 | 0.0037 |
| 5kb (8.2%) | 0.008 | 0.071 | 0.91 | 1.7 | 0.23 | 0.013 |
| 100kb (12.8%) | 0.0077 | 0.064 | 0.9 | 1.4 | 0.16 | 0.025 |

**Table S14.** S-LDSC results for joint model of published-restricted and boosted-restricted deep learning allelic-effect annotations restricted using S2G strategies, conditional on the baseline-LD-deep-S2G model annotations. Standardized Effect sizes (*τ\**) and Enrichment (E) of the significant SNP annotations in a joint model comprising of the marginally significant published-restricted and boosted-restricted SNP annotations corresponding to published and boosted deep learning allelic-effect annotations combined with S2G strategies. Results are conditional on 107 baseline-LD-deep-S2G model annotations (100 baseline-LD-deep and 7 additional annotations from Table S10). Results are meta-analyzed across 11 blood-related traits.

| Annotation | $\tau^*$ | $se(\tau^*)$ | $p(\tau^*)$ | $E$ | $se(E)$ | $p(E)$ |
| --- | --- | --- | --- | --- | --- | --- |
| DeepSEAΔ-boosted × eQTL<br>(0.6%) | 0.54 | 0.13 | 3.3e-05 | 11 | 1.4 | 0.00044 |
| BasenjiΔ-boosted × ABC<br>(0.6%) | 0.83 | 0.23 | 2e-04 | 14 | 2.1 | 1.4e-05 |
| BasenjiΔ-boosted × TSS<br>(0.9%) | 1.1 | 0.29 | 1e-04 | 16 | 1.8 | 2.7e-06 |

**Table S15.** Additional annotations of baseline-LD-deep-S2G-geneset model: List of 8 jointly significant gene set-specific S2G annotations from ref.^38^ added to the baseline-LD-deep-S2G model to create the baseline-LD-deep-S2G-geneset model. They include 7 annotations from the Enhancer-driven+PPI-enhancer joint model in ref^38^ and 1 jointly significant pLI S2G annotation.

| Annotations |
| --- |
| ATAC-distal × Promoter (1.8%) |
| EDS-binary × 100kb (14.6%) |
| SEG-GTEx × Coding (0.17%) |
| PPI-enhancer × ABC (0.58%) |
| PPI-enhancer × TSS (0.33%) |
| PPI-enhancer × Coding (0.24%) |
| PPI-enhancer × ATAC (0.41%) |
| pLI × Roadmap (0.56%) |

**Table S16.** S-LDSC results for gene set-specific boosted-restricted annotations, conditional on baseline-LD-deep-S2G-geneset model annotations: Standardized Effect sizes (*τ\**) and Enrichment (E) of 80 restricted SNP annotations corresponding to 4 allelic-effect annotations (DeepSEAΔ-boosted, BasenjiΔ-boosted, DeepBindΔ-boosted and deltaSVMΔ-boosted), 2 gene scores (PPI-enhancer and pLI) and 10 S2G strategies, conditional on 115 baseline-LD-deep-S2G-geneset annotations. Reports are meta-analyzed across 11 blood-related traits.

| DeepSEAΔ-boosted (PPI-enhancer) | | | | | | |
| --- | --- | --- | --- | --- | --- | --- |
| | $\tau^*$ | se($\tau^*$) | p($\tau^*$) | $E$ | se($E$) | p($E$) |
| ABC (0.23%) | 0.8 | 0.18 | 6.1e-06 | 21 | 2.6 | 1.2e-07 |
| TSS (0.16%) | 0.59 | 0.25 | 0.016 | 28 | 5.1 | 9.5e-05 |
| Coding (0.08%) | 0.66 | 0.19 | 0.00061 | 36 | 5.7 | 7.9e-05 |
| ATAC (0.12%) | 0.38 | 0.15 | 0.013 | 28 | 6.1 | 0.00019 |
| eQTL (0.09%) | 0.32 | 0.13 | 0.014 | 17 | 4.1 | 0.001 |
| Roadmap (0.31%) | 0.75 | 0.20 | 0.00026 | 23 | 3.9 | 2.1e-06 |
| Promoter (0.22%) | 0.61 | 0.17 | 0.00032 | 23 | 3.4 | 1.3e-05 |
| PC-HiC (1.98%) | 0.17 | 0.076 | 0.03 | 5.4 | 0.63 | 6e-07 |
| 5kb (1.47%) | 0.053 | 0.057 | 0.35 | 5.7 | 0.41 | 3.6e-07 |
| 100kb (3.54%) | 0.25 | 0.056 | 1.1e-05 | 5.2 | 0.26 | 3.2e-08 |
| BasenjiΔ-boosted (PPI-enhancer) | | | | | | |
| | $\tau^*$ | se($\tau^*$) | p($\tau^*$) | $E$ | se($E$) | p($E$) |
| ABC (0.32%) | 0.85 | 0.18 | 1.6e-06 | 19 | 2.2 | 2.5e-07 |
| TSS (0.21%) | 0.73 | 0.34 | 0.034 | 25 | 4.3 | 9.6e-06 |
| Coding (0.11%) | 0.75 | 0.25 | 0.0025 | 36 | 6.1 | 8.9e-06 |
| ATAC (0.18%) | 0.19 | 0.16 | 0.24 | 21 | 4.8 | 0.00017 |
| eQTL (0.13%) | 0.17 | 0.1 | 0.093 | 12 | 2.8 | 0.00019 |
| Roadmap (0.56%) | 0.74 | 0.19 | 0.00011 | 18 | 2.6 | 1.3e-08 |
| Promoter (0.31%) | 0.5 | 0.19 | 0.0072 | 19 | 3.1 | 3.4e-06 |
| PC-HiC (2.79%) | 0.2 | 0.077 | 0.0099 | 5.6 | 0.55 | 3.9e-09 |
| 5kb (2.04%) | 0.06 | 0.055 | 0.27 | 6.1 | 0.35 | 1.4e-08 |
| 100kb (4.70%) | 0.28 | 0.054 | 1.5e-07 | 5.6 | 0.3 | 8.1e-10 |
| DeepBindΔ-boosted (PPI-enhancer) | | | | | | |
| | $\tau^*$ | se($\tau^*$) | p($\tau^*$) | $E$ | se($E$) | p($E$) |
| ABC (0.12%) | 0.42 | 0.17 | 0.015 | 23 | 4.3 | 0.00036 |
| TSS (0.08%) | 0.26 | 0.26 | 0.31 | 28 | 8.2 | 0.005 |
| Coding (0.05%) | 0.013 | 0.15 | 0.93 | 18 | 5.3 | 0.024 |
| ATAC (0.08%) | 0.39 | 0.18 | 0.029 | 22 | 5.6 | 0.007 |
| eQTL (0.07%) | 0.26 | 0.14 | 0.064 | 15 | 5.2 | 0.023 |
| Roadmap (0.18%) | 0.70 | 0.18 | 0.0001 | 24 | 4.4 | 5.5e-05 |
| Promoter (0.13%) | 0.10 | 0.15 | 0.5 | 15 | 3.9 | 0.01 |
| PC-HiC (1.43%) | 0.24 | 0.094 | 0.0095 | 5.3 | 0.82 | 3.3e-05 |
| 5kb (1.05%) | 0.021 | 0.058 | 0.71 | 4.3 | 0.52 | 0.0002 |
| 100kb (2.86%) | 0.28 | 0.062 | 4.9e-06 | 4.5 | 0.33 | 2.6e-06 |
| deltaSVMΔ-boosted (PPI-enhancer) | | | | | | |
| | $\tau^*$ | se($\tau^*$) | p($\tau^*$) | $E$ | se($E$) | p($E$) |
| ABC (0.11%) | 0.85 | 0.17 | 2e-07 | 34 | 4.3 | 5.1e-05 |
| TSS (0.07%) | 0.88 | 0.32 | 0.0059 | 36 | 7.2 | 0.00037 |
| Coding (0.05%) | 0.29 | 0.18 | 0.11 | 29 | 6.7 | 0.0075 |
| ATAC (0.08%) | 0.53 | 0.18 | 0.0042 | 27 | 6.2 | 0.0056 |
| eQTL (0.08%) | 0.3 | 0.16 | 0.067 | 17 | 6.1 | 0.042 |
| Roadmap (0.17%) | 0.82 | 0.25 | 0.0003 | 29 | 5.5 | 5.1e-05 |
| Promoter (0.12%) | 0.19 | 0.17 | 0.26 | 16 | 4.4 | 0.012 |
| PC-HiC (1.37%) | 0.27 | 0.076 | 0.00048 | 5.2 | 0.63 | 5.4e-05 |
| 5kb (1.03%) | -0.0014 | 0.067 | 0.98 | 4 | 0.55 | 0.0012 |
| 100kb (2.88%) | 0.28 | 0.064 | 1.5e-05 | 4.4 | 0.35 | 1.6e-05 |
| DeepSEAΔ-boosted (pLI) | | | | | | |
| | $\tau^*$ | se($\tau^*$) | p($\tau^*$) | $E$ | se($E$) | p($E$) |
| ABC (0.32%) | 0.14 | 0.15 | 0.35 | 11 | 2 | 0.00077 |
| TSS (0.41%) | 0.088 | 0.17 | 0.6 | 13 | 1.7 | 0.00016 |
| Coding (0.24%) | 0.26 | 0.15 | 0.074 | 14 | 2.4 | 0.00032 |
| ATAC (0.16%) | 0.041 | 0.12 | 0.73 | 9.2 | 2.6 | 0.022 |
| eQTL (0.18%) | 0.14 | 0.098 | 0.14 | 8.4 | 2.1 | 0.005 |
| Roadmap (0.39%) | -0.19 | 0.21 | 0.37 | 7.4 | 2.2 | 0.066 |
| Promoter (0.68%) | 0.091 | 0.12 | 0.46 | 7.2 | 1.2 | 0.00055 |
| PC-HiC (3.55%) | 0.0045 | 0.085 | 0.96 | 3.6 | 0.42 | 1.3e-05 |
| 5kb (4.91%) | -0.015 | 0.049 | 0.77 | 3 | 0.18 | 9.3e-06 |
| 100kb (11.1%) | -0.079 | 0.063 | 0.2 | 2.3 | 0.16 | 2.6e-05 |
| BasenjiΔ-boosted (pLI) | | | | | | |
| | $\tau^*$ | se($\tau^*$) | p($\tau^*$) | $E$ | se($E$) | p($E$) |
| ABC (0.42%) | 0.26 | 0.15 | 0.075 | 12 | 1.7 | 3.6e-05 |
| TSS (0.49%) | 0.19 | 0.15 | 0.21 | 14 | 1.6 | 1.8e-05 |
| Coding (0.31%) | 0.28 | 0.14 | 0.042 | 14 | 2 | 3.7e-05 |
| ATAC (0.20%) | 0.11 | 0.13 | 0.4 | 12 | 3 | 0.0015 |
| eQTL (0.23%) | 0.053 | 0.099 | 0.59 | 7 | 1.8 | 0.0031 |
| Roadmap (0.59%) | 0.96 | 0.31 | 0.0017 | 16 | 2.4 | 5.2e-07 |
| Promoter (0.83%) | 0.15 | 0.14 | 0.27 | 8.1 | 1.2 | 3.1e-05 |
| PC-HiC (4.78%) | 0.061 | 0.068 | 0.37 | 4.1 | 0.32 | 1.6e-08 |
| 5kb (5.70%) | -0.019 | 0.049 | 0.69 | 3.5 | 0.16 | 5.6e-08 |
| 100kb (12.5%) | 0.015 | 0.064 | 0.81 | 3.1 | 0.17 | 1.1e-08 |
| DeepBindΔ-boosted (pLI) | | | | | | |
| | $\tau^*$ | se($\tau^*$) | p($\tau^*$) | $E$ | se($E$) | p($E$) |
| ABC (0.16%) | 0.14 | 0.16 | 0.39 | 13 | 3.7 | 0.011 |
| TSS (0.20%) | -0.061 | 0.16 | 0.7 | 13 | 2.9 | 0.0066 |
| Coding (0.17%) | -0.16 | 0.14 | 0.26 | 7.1 | 2.8 | 0.57 |
| ATAC (0.10%) | 0.35 | 0.12 | 0.0028 | 16 | 3.4 | 0.0056 |
| eQTL (0.12%) | 0.25 | 0.12 | 0.033 | 10 | 3.1 | 0.015 |
| Roadmap (0.21%) | 0.94 | 0.19 | 7.5e-07 | 23 | 3.1 | 2.5e-05 |
| Promoter (0.40%) | 0.12 | 0.16 | 0.43 | 8.4 | 2.2 | 0.013 |
| PC-HiC (2.54%) | 0.038 | 0.094 | 0.69 | 2.9 | 0.51 | 0.0042 |
| 5kb (3.68%) | 0.029 | 0.047 | 0.54 | 2.3 | 0.21 | 0.00097 |
| 100kb (9.58%) | -0.09 | 0.058 | 0.12 | 1.4 | 0.17 | 0.068 |
| deltaSVMΔ-boosted (pLI) | | | | | | |
| | $\tau^*$ | se($\tau^*$) | p($\tau^*$) | $E$ | se($E$) | p($E$) |
| ABC (0.15%) | 0.44 | 0.17 | 0.0086 | 18 | 3.6 | 0.0015 |
| TSS (0.18%) | 0.13 | 0.16 | 0.41 | 16 | 3.2 | 0.0013 |
| Coding (0.15%) | 0.29 | 0.13 | 0.029 | 16 | 2.8 | 0.001 |
| ATAC (0.10%) | 0.28 | 0.12 | 0.021 | 14 | 3.6 | 0.014 |
| eQTL (0.12%) | 0.24 | 0.13 | 0.067 | 10 | 3.4 | 0.026 |
| Roadmap (0.20%) | 0.92 | 0.18 | 3.2e-07 | 23 | 3.1 | 4.2e-05 |
| Promoter (0.37%) | -0.091 | 0.12 | 0.43 | 4.6 | 1.7 | 0.17 |
| PC-HiC (2.48%) | -0.0024 | 0.093 | 0.98 | 2.6 | 0.52 | 0.03 |
| 5kb (3.63%) | 0.00069 | 0.046 | 0.99 | 2.1 | 0.21 | 0.008 |
| 100kb (9.4%) | -0.056 | 0.056 | 0.31 | 1.5 | 0.16 | 0.055 |

**Table S17.** Standardized enrichment of gene set-specific boosted-restricted annotations, conditional on baseline-LD-deep-S2G-geneset model annotations: Standardized enrichment of 80 restricted SNP annotations corresponding to 4 allelic-effect annotations (DeepSEAΔ-boosted, BasenjiΔ-boosted, DeepBindΔ-boosted and deltaSVMΔ-boosted), 2 gene scores (PPI-enhancer and pLI) and 10 S2G strategies, conditional on 115 baseline-LD-deep-S2G-geneset annotations. Reports are meta-analyzed across 11 blood-related traits.

| DeepSEAD-boosted (PPI-enhancer) |  |  |  |
| --- | --- | --- | --- |
|  | *StdE* | *se(StdE)* | *p(StdE)* |
| ABC (0.23%) | 1.2 | 0.15 | 1.6e-05 |
| TSS (0.16%) | 1.1 | 0.2 | 9.5e-05 |
| Coding (0.08%) | 1 | 0.16 | 7.9e-05 |
| ATAC (0.12%) | 0.97 | 0.21 | 0.00019 |
| eQTL (0.09%) | 0.52 | 0.12 | 0.001 |
| Roadmap (0.31%) | 1.3 | 0.22 | 2.1e-06 |
| Promoter (0.22%) | 1.1 | 0.16 | 1.3e-05 |
| PC-HiC (1.98%) | 0.75 | 0.088 | 6e-07 |
| 5kb (1.47%) | 0.69 | 0.05 | 3.6e-07 |
| 100kb (3.54%) | 0.96 | 0.049 | 3.2e-08 |
| Basenji-boosted (PPI-enhancer) |  |  |  |
|  | *StdE* | *se(StdE)* | *p(StdE)* |
| ABC (0.32%) | 1.3 | 0.2 | 1.6e-06 |
| TSS (0.21%) | 1.1 | 0.2 | 9.6e-06 |
| Coding (0.11%) | 1.2 | 0.2 | 8.9e-06 |
| ATAC (0.18%) | 0.91 | 0.21 | 0.00017 |
| eQTL (0.13%) | 0.46 | 0.1 | 0.00019 |
| Roadmap (0.56%) | 1.3 | 0.2 | 1.3e-08 |
| Promoter (0.31%) | 1.1 | 0.17 | 3.4e-06 |
| PC-HiC (2.79%) | 0.93 | 0.09 | 3.9e-09 |
| 5kb (2.04%) | 0.86 | 0.05 | 1.4e-08 |
| 100kb (4.70%) | 1.2 | 0.063 | 8.1e-10 |
| DeepBind-boosted (PPI-enhancer) |  |  |  |
|  | *StdE* | *se(StdE)* | *p(StdE)* |
| ABC (0.12%) | 0.78 | 0.15 | 0.00036 |
| TSS (0.08%) | 0.76 | 0.23 | 0.005 |
| Coding (0.05%) | 0.43 | 0.12 | 0.024 |
| ATAC (0.08%) | 0.63 | 0.16 | 0.007 |
| eQTL (0.07%) | 0.4 | 0.13 | 0.023 |
| Roadmap (0.18%) | 1 | 0.19 | 5.5e-05 |
| Promoter (0.13%) | 0.53 | 0.14 | 0.01 |
| PC-HiC (1.43%) | 0.63 | 0.097 | 3.3e-05 |
| 5kb (1.05%) | 0.44 | 0.053 | 0.0002 |
| 100kb (2.86%) | 0.75 | 0.055 | 2.6e-06 |
| deltaSVM-boosted (PPI-enhancer) |  |  |  |
|  | *StdE* | *se(StdE)* | *p(StdE)* |
| ABC (0.11%) | 1.1 | 0.14 | 5.1e-05 |
| TSS (0.07%) | 1.2 | 0.27 | 0.00037 |
| Coding (0.05%) | 0.64 | 0.15 | 0.0075 |
| ATAC (0.08%) | 0.71 | 0.17 | 0.0056 |
| eQTL (0.08%) | 0.42 | 0.16 | 0.042 |
| Roadmap (0.17%) | 1.2 | 0.23 | 5.1e-05 |
| Promoter (0.12%) | 0.54 | 0.15 | 0.012 |
| PC-HiC (1.37%) | 0.6 | 0.073 | 5.4e-05 |
| 5kb (1.03%) | 0.4 | 0.055 | 0.0012 |
| 100kb (2.88%) | 0.72 | 0.058 | 1.6e-05 |
| DeepSEAD-boosted (pLI) |  |  |  |
|  | *StdE* | *se(StdE)* | *p(StdE)* |
| ABC (0.32%) | 0.61 | 0.11 | 0.00077 |
| TSS (0.41%) | 0.8 | 0.1 | 0.00016 |
| Coding (0.24%) | 0.66 | 0.11 | 0.00032 |
| ATAC (0.16%) | 0.33 | 0.092 | 0.022 |
| eQTL (0.18%) | 0.33 | 0.082 | 0.005 |
| Roadmap (0.39%) | 0.42 | 0.13 | 0.066 |
| Promoter (0.68%) | 0.56 | 0.089 | 0.00055 |
| PC-HiC (3.55%) | 0.63 | 0.073 | 1.3e-05 |
| 5kb (4.91%) | 0.6 | 0.037 | 9.3e-06 |
| 100kb (11.1%) | 0.7 | 0.049 | 2.6e-05 |

BasenjiΔ-boosted (pLI)
|  | *StdE* | *se(StdE)* | *p(StdE)* |
| --- | --- | --- | --- |
| ABC (0.42%) | 0.76 | 0.1 | 3.6e-05 |
| TSS (0.49%) | 0.95 | 0.1 | 1.8e-05 |
| Coding (0.31%) | 0.73 | 0.11 | 3.7e-05 |
| ATAC (0.20%) | 0.49 | 0.12 | 0.0015 |
| eQTL (0.23%) | 0.3 | 0.08 | 0.0031 |
| Roadmap (0.59%) | 1.1 | 0.17 | 5.2e-07 |
| Promoter (0.83%) | 0.68 | 0.1 | 3.1e-05 |
| PC-HiC (4.78%) | 0.83 | 0.066 | 1.6e-08 |
| 5kb (5.70%) | 0.76 | 0.035 | 5.6e-08 |
| 100kb (12.5%) | 1 | 0.055 | 1.1e-08 |

DeepBindΔ-boosted (pLI)
|  | *StdE* | *se(StdE)* | *p(StdE)* |
| --- | --- | --- | --- |
| ABC (0.16%) | 0.49 | 0.14 | 0.011 |
| TSS (0.20%) | 0.54 | 0.12 | 0.0066 |
| Coding (0.17%) | 0.27 | 0.11 | 0.57 |
| ATAC (0.10%) | 0.45 | 0.096 | 0.0056 |
| eQTL (0.12%) | 0.33 | 0.098 | 0.015 |
| Roadmap (0.21%) | 0.95 | 0.13 | 2.5e-05 |
| Promoter (0.40%) | 0.5 | 0.13 | 0.013 |
| PC-HiC (2.54%) | 0.44 | 0.076 | 0.0042 |
| 5kb (3.68%) | 0.41 | 0.038 | 0.00097 |
| 100kb (9.58%) | 0.42 | 0.048 | 0.068 |

deltaSVMΔ-boosted (pLI)
|  | *StdE* | *se(StdE)* | *p(StdE)* |
| --- | --- | --- | --- |
| ABC (0.15%) | 0.66 | 0.14 | 0.0015 |
| TSS (0.18%) | 0.64 | 0.13 | 0.0013 |
| Coding (0.15%) | 0.58 | 0.1 | 0.001 |
| ATAC (0.10%) | 0.39 | 0.097 | 0.014 |
| v eQTL (0.12%) | 0.32 | 0.11 | 0.026 |
| Roadmap (0.20%) | 0.93 | 0.13 | 4.2e-05 |
| Promoter (0.37%) | 0.38 | 0.098 | 0.023 |
| PC-HiC (2.48%) | 0.46 | 0.076 | 0.0043 |
| 5kb (3.63%) | 0.4 | 0.037 | 0.0021 |
| 100kb (9.4%) | 0.4 | 0.047 | 0.081 |

**Table S18.** S-LDSC results for joint model of gene set-specific boosted-restricted annotations, conditional on baseline-LD-deep-S2G-geneset model annotations: Standardized Effect sizes (*τ\**) and Enrichment (E) of the Bonferroni significant boosted-restricted S2G annotations linked to PPI-enhancer genes that were marginally significant in Figure 2. The results were conditioned either on 115 baseline- LD-deep-S2G-geneset annotations, or baseline-LD-deep-S2G-geneset plus 3 annotations from Figure 1, or baseline-LD-deep-S2G-geneset plus 3 annotations from Figure 1 plus BasenjiΔ-boosted*×*Roadmap. Reports are meta-analyzed across 11 blood-related traits.

| | $\tau^*$ | se($\tau^*$) | p($\tau^*$) | $E$ | se($E$) | p($E$) |
| --- | --- | --- | --- | --- | --- | --- |
| baseline-LD-deep-S2G-geneset model |  |  |  |  |  |  |
| BasenjiΔ-boosted<br>(PPI-enhancer) × ABC (0.32%) | 0.72 | 0.17 | 3.2e-05 | 17 | 1.8 | 9.5e-07 |
| BasenjiΔ-boosted<br>(PPI-enhancer) × Roadmap (0.56%) | 0.68 | 0.18 | 9.7e-05 | 17 | 2.6 | 1.9e-08 |
| baseline-LD-deep-S2G-geneset + 3 annotations from Figure 1 |  |  |  |  |  |  |
| BasenjiΔ-boosted<br>(PPI-enhancer) × ABC (0.32%) | 0.69 | 0.18 | 0.0001 | 17 | 1.8 | 9.7e-07 |
| BasenjiΔ-boosted<br>(PPI-enhancer) × Roadmap (0.56%) | 0.66 | 0.18 | 0.0002 | 18 | 2.7 | 1.4e-08 |
| baseline-LD-deep-S2G-geneset + 3 annotations from Figure 1 + BasenjiΔ-boosted×Roadmap | | | | | | |
| BasenjiΔ-boosted<br>(PPI-enhancer) × ABC (0.32%) | 0.63 | 0.18 | 0.0003 | 16 | 1.5 | 8.2e-06 |
| BasenjiΔ-boosted (PPI-enhancer)<br>× Roadmap (0.56%) | 0.64 | 0.18 | 0.0003 | 18 | 2.8 | 3.3e-08 |

**Table S19.** S-LDSC results for boosted-restricted deep learning allelic-effect annotations restricted using S2G strategies, conditional on the baseline-LD- deep-S2G model annotations plus the local GC-content annotation and annotations restricted using the local GC-content annotation: Standardized Effect sizes (*τ\**) and Enrichment (E) of restricted SNP annotations corresponding to each of DeepSEAΔ-boosted, BasenjiΔ-boosted, DeepbindΔ-boosted and deltaSVMΔ-boosted annotations restricted using the local GC-content and 10 S2G strategies conditional on 100 baseline-LD-deep annotations and unrestricted S2G annotations and S2G annotations restricted using local GC-content annotation. Reports are meta-analyzed across 11 blood-related traits.

| DeepSEAΔ-boosted | | | | | | |
| --- | --- | --- | --- | --- | --- | --- |
| | $\tau^*$ | se($\tau^*$) | p($\tau^*$) | $E$ | se($E$) | p($E$) |
| ABC (0.50%) | 0.22 | 0.18 | 0.21 | 10 | 1.8 | 0.00082 |
| TSS (0.75%) | 0.5 | 0.22 | 0.023 | 15 | 1.7 | 2.4e-05 |
| Coding (0.47%) | 0.6 | 0.2 | 0.0023 | 13 | 1.9 | 9.2e-05 |
| ATAC (0.46%) | 0.11 | 0.14 | 0.42 | 9.4 | 1.9 | 0.00078 |
| eQTL (0.59%) | 0.57 | 0.14 | 8.3e-05 | 11 | 1.4 | 0.00025 |
| Roadmap (1.0%) | 0.21 | 0.18 | 0.25 | 9.3 | 1.4 | 0.0001 |
| Promoter (1.33%) | 0.35 | 0.21 | 0.092 | 7.6 | 1.2 | 0.00019 |
| PC-HiC (5.89%) | -0.25 | 0.13 | 0.047 | 2.9 | 0.5 | 0.0021 |
| 5kb (9.89%) | -0.11 | 0.09 | 0.2 | 2.4 | 0.23 | 0.00028 |
| 100kb (13.7%) | -0.1 | 0.089 | 0.25 | 2.1 | 0.2 | 0.00047 |
| BasenjiΔ-boosted | | | | | | |
| | $\tau^*$ | se($\tau^*$) | p($\tau^*$) | $E$ | se($E$) | p($E$) |
| ABC (0.63%) | 1.1 | 0.29 | 0.00018 | 15 | 2.3 | 6.1e-06 |
| TSS (0.88%) | 1.4 | 0.36 | 6.5e-05 | 18 | 1.9 | 3.7e-07 |
| Coding (0.59%) | 0.76 | 0.23 | 0.0011 | 12 | 1.7 | 3.7e-05 |
| ATAC (0.60%) | 0.38 | 0.22 | 0.08 | 12 | 2.2 | 1.1e-05 |
| eQTL (0.79%) | 0.17 | 0.1 | 0.11 | 7.4 | 0.87 | 0.00011 |
| Roadmap (1.54%) | 0.76 | 0.27 | 0.0051 | 12 | 1.7 | 3e-08 |
| Promoter (1.63%) | 0.31 | 0.14 | 0.025 | 7.6 | 0.83 | 4.6e-06 |
| PC-HiC (7.67%) | 0.18 | 0.14 | 0.19 | 4.4 | 0.45 | 2.4e-09 |
| 5kb (11.2%) | 0.031 | 0.068 | 0.64 | 3.2 | 0.15 | 2.9e-08 |
| 100kb (14.5%) | 0.13 | 0.07 | 0.064 | 3.3 | 0.2 | 6.6e-09 |
| DeepBindΔ-boosted | | | | | | |
| | $\tau^*$ | se($\tau^*$) | p($\tau^*$) | $E$ | se($E$) | p($E$) |
| ABC (0.27%) | 0.025 | 0.21 | 0.91 | 8.7 | 3.6 | 0.13 |
| TSS (0.37%) | -0.11 | 0.16 | 0.49 | 12 | 2.5 | 0.002 |
| Coding (0.34%) | -0.22 | 0.14 | 0.11 | 5.2 | 1.8 | 0.22 |
| ATAC (0.31%) | 0.27 | 0.14 | 0.059 | 11 | 2.1 | 0.0016 |
| eQTL (0.43%) | -0.37 | 0.17 | 0.029 | 0.73 | 2.1 | 0.78 |
| Roadmap (0.57%) | -0.028 | 0.2 | 0.89 | 6.4 | 2.1 | 0.032 |
| Promoter (0.84%) | -0.19 | 0.21 | 0.38 | 4.1 | 1.9 | 0.28 |
| PC-HiC (4.46%) | -0.06 | 0.14 | 0.66 | 2.2 | 0.55 | 0.11 |
| 5kb (8.24%) | -0.14 | 0.1 | 0.17 | 1.3 | 0.32 | 0.63 |
| 100kb (12.9%) | -0.06 | 0.1 | 0.56 | 1.3 | 0.26 | 0.56 |
| deltaSVMΔ-boosted | | | | | | |
| | $\tau^*$ | se($\tau^*$) | p($\tau^*$) | $E$ | se($E$) | p($E$) |
| ABC (0.25%) | 0.44 | 0.17 | 0.009 | 14 | 2.8 | 0.0014 |
| TSS (0.33%) | 0.29 | 0.26 | 0.26 | 18 | 4.3 | 0.0008 |
| Coding (0.31%) | 0.37 | 0.24 | 0.12 | 13 | 3.5 | 0.002 |
| ATAC (0.28%) | 0.039 | 0.13 | 0.77 | 7.2 | 2.2 | 0.6 |
| eQTL (0.41%) | 0.11 | 0.18 | 0.53 | 6.5 | 2.4 | 0.43 |
| Roadmap (0.53%) | 0.11 | 0.18 | 0.56 | 7.7 | 2.1 | 0.03 |
| Promoter (0.77%) | -0.0084 | 0.16 | 0.96 | 5.2 | 1.5 | 0.027 |
| PC-HiC (4.3%) | 0.013 | 0.17 | 0.94 | 2.5 | 0.64 | 0.041 |
| 5kb (8.2%) | 0.014 | 0.14 | 0.92 | 1.7 | 0.43 | 0.072 |
| 100kb (12.8%) | 0.034 | 0.1 | 0.74 | 1.5 | 0.24 | 0.073 |

**Table S20.**
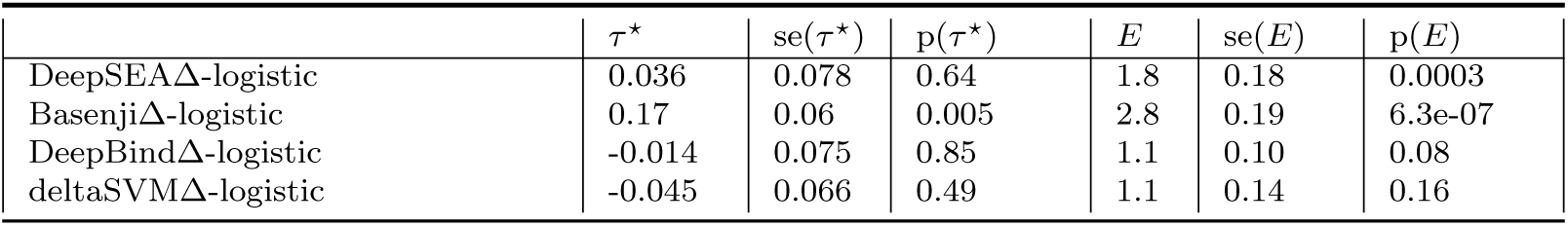
S-LDSC results for annotations generated using the logistic regression model: Standardized Effect sizes (*τ\**) and Enrichment (E) of SNP annotations generated by training fine-mapped SNPs on features from DeepSEA, Basenji, DeepBind and deltaSVM approaches using the logistic regression model (instead of the gradient boosting model). Results are conditional on 100 baseline-LD-deep annotations. Reports are meta-analyzed across 11 blood-related traits.

| | $\tau^*$ | $se(\tau^*)$ | $p(\tau^*)$ | $E$ | $se(E)$ | $p(E)$ |
| --- | --- | --- | --- | --- | --- | --- |
| DeepSEAΔ-logistic | 0.036 | 0.078 | 0.64 | 1.8 | 0.18 | 0.0003 |
| BasenjiΔ-logistic | 0.17 | 0.06 | 0.005 | 2.8 | 0.19 | 6.3e-07 |
| DeepBindΔ-logistic | -0.014 | 0.075 | 0.85 | 1.1 | 0.10 | 0.08 |
| deltaSVMΔ-logistic | -0.045 | 0.066 | 0.49 | 1.1 | 0.14 | 0.16 |

**Table S21.** S-LDSC results for boosted-restricted deep learning allelic-effect annotations generated using the logistic regression model: Standardized Effect sizes (*τ\**) and Enrichment (E) of restricted SNP annotations (across 10 S2G strategies) corresponding to annotations generated by training fine-mapped SNPs on features from DeepSEA, Basenji, DeepBind and deltaSVM approaches using the logistic regression model (instead of the gradient boosting model). Results are conditional on 107 baseline- LD-deep-S2G annotations (100 baseline-LD-deep and 7 additional annotations from Table S10). Results are meta-analyzed across 11 blood-related traits.

| DeepSEAΔ-boosted | | | | | | |
| --- | --- | --- | --- | --- | --- | --- |
| | $\tau^*$ | se($\tau^*$) | p($\tau^*$) | $E$ | se($E$) | p($E$) |
| ABC (0.44%) | 0.20 | 0.14 | 0.15 | 8 | 2.3 | 0.004 |
| TSS (0.68%) | 0.28 | 0.19 | 0.14 | 11 | 2.1 | 0.0001 |
| Coding (0.39%) | 0.30 | 0.17 | 0.077 | 11 | 1.9 | 0.0005 |
| ATAC (0.50%) | 0.08 | 0.05 | 0.11 | 8.2 | 1.4 | 0.0001 |
| eQTL (0.47%) | 0.49 | 0.13 | 0.0001 | 11 | 1.4 | 0.0001 |
| Roadmap (0.84%) | 0.44 | 0.28 | 0.12 | 8.6 | 1.2 | 9.8e-05 |
| Promoter (1.04%) | 0.13 | 0.24 | 0.58 | 5.4 | 1.0 | 0.076 |
| PC-HiC (5.06%) | -0.005 | 0.10 | 0.88 | 2.8 | 0.44 | 0.0003 |
| 5kb (8.18%) | 0.05 | 0.078 | 0.52 | 1.9 | 0.20 | 0.002 |
| 100kb (11.4%) | 0.03 | 0.044 | 0.47 | 1.4 | 0.2 | 0.004 |
| BasenjiΔ-boosted | | | | | | |
| | $\tau^*$ | se($\tau^*$) | p($\tau^*$) | $E$ | se($E$) | p($E$) |
| ABC (0.48%) | 0.95 | 0.27 | 0.0004 | 14 | 2.4 | 2.2e-05 |
| TSS (0.76%) | 1.2 | 0.35 | 0.0006 | 16 | 1.7 | 8.9e-06 |
| Coding (0.27%) | 0.29 | 0.16 | 0.078 | 11 | 1.7 | 2.8e-05 |
| ATAC (0.20%) | -0.013 | 0.11 | 0.91 | 10 | 1.8 | 2.8e-05 |
| eQTL (0.38%) | -0.13 | 0.095 | 0.15 | 7.0 | 0.87 | 0.0008 |
| Roadmap (0.48%) | 0.24 | 0.12 | 0.054 | 10 | 1.8 | 5.2e-06 |
| Promoter (0.66%) | -0.061 | 0.12 | 0.61 | 5.9 | 0.78 | 2.8e-05 |
| PC-HiC (2.58%) | -0.025 | 0.087 | 0.77 | 3.1 | 0.56 | 1.8e-06 |
| 5kb (6.48%) | 0.0031 | 0.061 | 0.96 | 2.1 | 0.17 | 3.1e-07 |
| 100kb (9.73%) | -0.01 | 0.054 | 0.85 | 2.2 | 0.19 | 2.8e-05 |
| DeepBindΔ-boosted | | | | | | |
| | $\tau^*$ | se($\tau^*$) | p($\tau^*$) | $E$ | se($E$) | p($E$) |
| ABC (0.19%) | -0.16 | 0.15 | 0.28 | 3.3 | 3.1 | 0.34 |
| TSS (0.22%) | -0.13 | 0.18 | 0.47 | 8.2 | 3.5 | 0.56 |
| Coding (0.21%) | 0.37 | 0.13 | 0.0043 | 13 | 2.5 | 0.003 |
| ATAC (0.23%) | 0.07 | 0.14 | 0.63 | 6.4 | 2.4 | 0.087 |
| eQTL (0.35%) | 0.28 | 0.19 | 0.14 | 7.8 | 2.7 | 0.10 |
| Roadmap (0.44%) | 0.061 | 0.21 | 0.77 | 6.9 | 2.2 | 0.052 |
| Promoter (0.61%) | 0.049 | 0.13 | 0.7 | 4.3 | 1.4 | 0.13 |
| PC-HiC (4.01%) | 0.26 | 0.12 | 0.027 | 3.1 | 0.51 | 0.004 |
| 5kb (7.90%) | 0.13 | 0.095 | 0.17 | 1.8 | 0.3 | 0.069 |
| 100kb (12.5%) | 0.24 | 0.087 | 0.0063 | 1.7 | 0.22 | 0.027 |
| deltaSVMΔ-boosted | | | | | | |
| | $\tau^*$ | se($\tau^*$) | p($\tau^*$) | $E$ | se($E$) | p($E$) |
| ABC (0.20%) | -0.25 | 0.15 | 0.1 | 2.5 | 2.8 | 0.82 |
| TSS (0.25%) | 0.18 | 0.17 | 0.29 | 15 | 3 | 0.0066 |
| Coding (0.22%) | 0.029 | 0.19 | 0.88 | 7.6 | 3.8 | 0.15 |
| ATAC (0.25%) | 0.017 | 0.14 | 0.9 | 5.7 | 2.4 | 0.25 |
| eQTL (0.37%) | 0.19 | 0.24 | 0.42 | 5.5 | 2.8 | 0.25 |
| Roadmap (0.48%) | 0.16 | 0.18 | 0.35 | 7.8 | 2.2 | 0.042 |
| Promoter (0.67%) | -0.1 | 0.14 | 0.47 | 3.3 | 1.5 | 0.041 |
| PC-HiC (4.29%) | -0.18 | 0.14 | 0.18 | 1.2 | 0.58 | 0.36 |
| 5kb (8.10%) | 0.084 | 0.13 | 0.51 | 1.7 | 0.4 | 0.77 |
| 100kb (12.8%) | 0.08 | 0.11 | 0.49 | 1.4 | 0.29 | 0.99 |

**Table S22.** S-LDSC results for gene set-specific published-restricted annotations, conditional on baseline-LD-deep-S2G-geneset model annotations: Standardized Effect sizes (*τ\**) and Enrichment (E) of 80 published-restricted SNP annotations corresponding to the 4 models (DeepSEAΔ-published, BasenjiΔ-published, DeepBindΔ- published, deltaSVMΔ-published) for which we observed significant enrichment signal for the published allelic-effect annotations, 2 gene scores (PPI-enhancer and pLI) and 10 S2G strategies, conditional on 115 baseline-LD-deep-S2G-geneset annotations. Reports are meta-analyzed across 11 blood-related traits.

| DeepSEAΔ-published (PPI-enhancer) | | | | | | |
| --- | --- | --- | --- | --- | --- | --- |
| | $\tau^*$ | se($\tau^*$) | p($\tau^*$) | $E$ | se($E$) | p($E$) |
| ABC (0.13%) | 0.69 | 0.18 | 0.00013 | 24 | 3.1 | 1.6e-05 |
| TSS (0.08%) | 0.54 | 0.22 | 0.015 | 22 | 2.6 | 2.5e-06 |
| Coding (0.04%) | 0.74 | 0.21 | 0.00055 | 27 | 3.1 | 4.1e-06 |
| ATAC (0.07%) | 0.66 | 0.22 | 0.002 | 23 | 3.9 | 2.4e-07 |
| eQTL (0.06%) | 0.16 | 0.082 | 0.049 | 9.6 | 1.5 | 3e-06 |
| Roadmap (0.18%) | 0.61 | 0.16 | 0.00015 | 15 | 1.9 | 1.2e-09 |
| Promoter (0.12%) | 0.34 | 0.13 | 0.006 | 15 | 1.5 | 1.9e-06 |
| PC-HiC (1.28%) | 0.13 | 0.044 | 0.0041 | 4 | 0.25 | 1.6e-10 |
| 5kb (0.93%) | 0.018 | 0.03 | 0.56 | 4.5 | 0.17 | 5.4e-10 |
| 100kb (2.43%) | 0.14 | 0.03 | 3.4e-06 | 3.6 | 0.13 | 5.3e-11 |
| BasenjiΔ-published (PPI-enhancer) | | | | | | |
| | $\tau^*$ | se($\tau^*$) | p($\tau^*$) | $E$ | se($E$) | p($E$) |
| ABC (0.14%) | 0.78 | 0.21 | 0.0001 | 23 | 3.5 | 1.6e-06 |
| TSS (0.09%) | 0.65 | 0.26 | 0.014 | 23 | 2.6 | 2.6e-06 |
| Coding (0.05%) | 0.73 | 0.22 | 0.001 | 26 | 3 | 2.6e-06 |
| ATAC (0.08%) | 0.51 | 0.21 | 0.017 | 19 | 3.3 | 5e-07 |
| eQTL (0.07%) | 0.17 | 0.08 | 0.038 | 9.3 | 1.4 | 8.7e-07 |
| Roadmap (0.23%) | 0.55 | 0.16 | 0.00061 | 13 | 1.7 | 1.1e-09 |
| Promoter (0.15%) | 0.37 | 0.14 | 0.0062 | 15 | 1.4 | 6.1e-07 |
| PC-HiC (1.59%) | 0.12 | 0.046 | 0.0083 | 3.9 | 0.23 | 5.7e-11 |
| 5kb (1.16%) | 0.028 | 0.03 | 0.35 | 4.5 | 0.18 | 2.9e-10 |
| 100kb (2.99%) | 0.14 | 0.029 | 8.8e-07 | 3.5 | 0.13 | 3.2e-11 |
| DeepBindΔ-published (PPI-enhancer) | | | | | | |
| | $\tau^*$ | se($\tau^*$) | p($\tau^*$) | $E$ | se($E$) | p($E$) |
| ABC (0.34%) | 0.88 | 0.18 | 1.4e-06 | 17 | 2 | 2.7e-07 |
| TSS (0.19%) | 0.53 | 0.23 | 0.021 | 21 | 2 | 9.3e-07 |
| Coding (0.14%) | 0.71 | 0.19 | 0.00019 | 20 | 1.9 | 2.6e-06 |
| ATAC (0.24%) | 0.66 | 0.17 | 7.9e-05 | 16 | 2.7 | 2.2e-07 |
| eQTL (0.22%) | 0.15 | 0.069 | 0.029 | 7.6 | 1.1 | 2.1e-06 |
| Roadmap (0.62%) | 0.56 | 0.14 | 5.1e-05 | 12 | 1.4 | 1e-09 |
| Promoter (0.38%) | 0.15 | 0.096 | 0.11 | 11 | 0.98 | 1.8e-06 |
| PC-HiC (5.2%) | 0.11 | 0.034 | 0.00067 | 3.1 | 0.16 | 2.3e-10 |
| 5kb (3.8%) | 0.019 | 0.023 | 0.4 | 3.4 | 0.12 | 3.4e-10 |
| 100kb (10.2%) | 0.11 | 0.024 | 2.9e-06 | 2.7 | 0.089 | 8.8e-11 |
| deltaSVMΔ-published (PPI-enhancer) | | | | | | |
| | $\tau^*$ | se($\tau^*$) | p($\tau^*$) | $E$ | se($E$) | p($E$) |
| ABC (0.21%) | 0.69 | 0.18 | 0.00011 | 18 | 2.2 | 2.7e-07 |
| TSS (0.12%) | 0.42 | 0.21 | 0.043 | 21 | 2.2 | 1.6e-06 |
| Coding (0.09%) | 0.6 | 0.19 | 0.0017 | 21 | 2.1 | 5.6e-06 |
| ATAC (0.15%) | 0.52 | 0.17 | 0.0021 | 16 | 2.7 | 5.2e-07 |
| eQTL (0.14%) | 0.15 | 0.069 | 0.027 | 7.7 | 1.2 | 5.2e-06 |
| Roadmap (0.39%) | 0.59 | 0.15 | 5.6e-05 | 12 | 1.5 | 1.9e-09 |
| Promoter (0.24%) | 0.15 | 0.096 | 0.12 | 11 | 1 | 2.1e-06 |
| PC-HiC (3.27%) | 0.12 | 0.033 | 0.00038 | 3.1 | 0.16 | 5.1e-10 |
| 5kb (2.41%) | 0.017 | 0.025 | 0.5 | 3.4 | 0.12 | 5.4e-10 |
| 100kb (6.56%) | 0.12 | 0.025 | 2.5e-06 | 2.7 | 0.091 | 1.4e-10 |
| DeepSEAΔ-published (pLI) | | | | | | |
| | $\tau^*$ | se($\tau^*$) | p($\tau^*$) | $E$ | se($E$) | p($E$) |
| ABC (0.17%) | 0.064 | 0.2 | 0.74 | 9.6 | 1.3 | 8.8e-06 |
| TSS (0.22%) | -0.11 | 0.15 | 0.48 | 12 | 1.1 | 9e-06 |
| Coding (0.13%) | 0.11 | 0.13 | 0.4 | 11 | 1.2 | 1.1e-05 |
| ATAC (0.09%) | 0.17 | 0.094 | 0.07 | 10 | 1.8 | 4.8e-05 |
| eQTL (0.11%) | 0.063 | 0.075 | 0.4 | 5.8 | 0.93 | 0.00013 |
| Roadmap (0.21%) | -0.24 | 0.34 | 0.47 | 8 | 1 | 2.1e-05 |
| Promoter (0.38%) | -0.0007 | 0.11 | 0.99 | 6.5 | 0.63 | 1.3e-05 |
| PC-HiC (2.29%) | -0.0093 | 0.052 | 0.86 | 3 | 0.12 | 7.1e-10 |
| 5kb (3.25%) | -0.012 | 0.022 | 0.59 | 2.4 | 0.052 | 5.7e-09 |
| 100kb (8.12%) | -0.048 | 0.025 | 0.054 | 1.9 | 0.038 | 3.3e-10 |

BasenjiΔ-published (pLI)
| | $\tau^*$ | se($\tau^*$) | p($\tau^*$) | $E$ | se($E$) | p($E$) |
| --- | --- | --- | --- | --- | --- | --- |
| ABC (0.19%) | -0.058 | 0.16 | 0.72 | 9.6 | 1.1 | 9.6e-06 |
| TSS (0.24%) | -0.13 | 0.14 | 0.35 | 12 | 1.1 | 1.4e-05 |
| Coding (0.16%) | 0.1 | 0.14 | 0.44 | 11 | 1.2 | 7.8e-06 |
| ATAC (0.10%) | 0.14 | 0.091 | 0.13 | 9.6 | 1.6 | 2.7e-05 |
| eQTL (0.13%) | 0.054 | 0.067 | 0.41 | 5.7 | 0.79 | 5.1e-05 |
| Roadmap (0.25%) | 0.16 | 0.5 | 0.75 | 10 | 1.4 | 3e-07 |
| Promoter (0.44%) | -0.0033 | 0.11 | 0.98 | 6.5 | 0.59 | 7e-06 |
| PC-HiC (2.79%) | -0.02 | 0.049 | 0.68 | 3 | 0.11 | 1.7e-10 |
| 5kb (3.89%) | -0.016 | 0.021 | 0.46 | 2.4 | 0.047 | 1.9e-09 |
| 100kb (9.76%) | -0.045 | 0.023 | 0.053 | 1.9 | 0.035 | 8.2e-11 |

DeepBindΔ-published (pLI)
| | $\tau^*$ | se($\tau^*$) | p($\tau^*$) | $E$ | se($E$) | p($E$) |
| --- | --- | --- | --- | --- | --- | --- |
| ABC (0.48%) | 0.44 | 0.17 | 0.009 | 9.1 | 0.93 | 3.2e-06 |
| TSS (0.51%) | 0.29 | 0.26 | 0.26 | 13 | 0.97 | 3.6e-06 |
| Coding (0.45%) | 0.37 | 0.24 | 0.12 | 9.6 | 0.93 | 3.7e-06 |
| ATAC (0.31%) | 0.039 | 0.13 | 0.77 | 8.4 | 1.4 | 7e-06 |
| eQTL (0.42%) | 0.11 | 0.18 | 0.53 | 4.9 | 0.72 | 8.8e-05 |
| Roadmap (0.70%) | 0.11 | 0.18 | 0.56 | 9 | 0.86 | 5.9e-08 |
| Promoter (1.19%) | -0.0084 | 0.16 | 0.96 | 5.4 | 0.52 | 2.2e-05 |
| PC-HiC (9.2%) | 0.013 | 0.17 | 0.94 | 2.3 | 0.08 | 2.1e-10 |
| 5kb (13.9%) | 0.014 | 0.14 | 0.92 | 1.8 | 0.046 | 1.7e-08 |
| 100kb (28%) | 0.034 | 0.1 | 0.74 | 1.4 | 0.022 | 3.4e-10 |

deltaSVMΔ-published (pLI)
| | $\tau^*$ | se($\tau^*$) | p($\tau^*$) | $E$ | se($E$) | p($E$) |
| --- | --- | --- | --- | --- | --- | --- |
| ABC (0.31%) | 0.18 | 0.16 | 0.25 | 9.2 | 0.99 | 3.3e-06 |
| TSS (0.33%) | 0.11 | 0.13 | 0.38 | 13 | 1 | 3.9e-06 |
| Coding (0.28%) | 0.032 | 0.12 | 0.79 | 9.7 | 0.98 | 5.6e-06 |
| ATAC (0.19%) | 0.2 | 0.089 | 0.023 | 8.4 | 1.4 | 1.8e-05 |
| eQTL (0.27%) | 0.11 | 0.065 | 0.093 | 4.9 | 0.78 | 0.00018 |
| Roadmap (0.45%) | 0.7 | 0.1 | 1.8e-11 | 9.4 | 0.93 | 8.3e-08 |
| Promoter (0.76%) | 0.068 | 0.076 | 0.37 | 5.4 | 0.58 | 3.2e-05 |
| PC-HiC (5.87%) | 0.032 | 0.045 | 0.48 | 2.3 | 0.088 | 5.3e-10 |
| 5kb (8.78%) | 0.025 | 0.017 | 0.13 | 1.8 | 0.052 | 3.4e-08 |
| 100kb (22%) | -0.025 | 0.018 | 0.17 | 1.4 | 0.028 | 5.5e-10 |

**Table S23.** S-LDSC results for gene set-specific published-restricted annotations, conditional on baseline-LD-deep-S2G-geneset model annotations plus the 2 jointly significant gene set-specific boosted-restricted annotations from Figure 2 Panel C: Standardized Effect sizes (*τ\**) and Enrichment (E) of 80 restricted SNP annotations corresponding to 4 published allelic effect annotations (DeepSEAΔ- published, BasenjiΔ-published, DeepBindΔ-published and deltaSVMΔ-published) for which we observed significant enrichment signal for the published allelic-effect annotations, 2 gene scores (PPI-enhancer and pLI) and 10 S2G strategies, conditional on 115 baseline-LD-deep-S2G-geneset annotations and 2 jointly significant annotations from Figure 2 Panel C. Reports are meta-analyzed across 11 blood-related traits.

| DeepSEADΔ-published (PPI-enhancer) | | | | | | |
| --- | --- | --- | --- | --- | --- | --- |
| | $\tau^*$ | se($\tau^*$) | p($\tau^*$) | $E$ | se($E$) | p($E$) |
| ABC (0.13%) | 0.46 | 0.22 | 0.035 | 18 | 2.1 | 3.7e-07 |
| TSS (0.08%) | 0.55 | 0.18 | 0.002 | 19 | 3.3 | 2.6e-06 |
| Coding (0.04%) | 0.5 | 0.21 | 0.016 | 23 | 2.8 | 3.8e-05 |
| ATAC (0.07%) | 0.66 | 0.22 | 0.002 | 23 | 3.9 | 2.4e-07 |
| eQTL (0.06%) | 0.092 | 0.078 | 0.24 | 8.8 | 1.5 | 1.2e-05 |
| Roadmap (0.18%) | -0.086 | 0.21 | 0.69 | 13 | 1.6 | 1.4e-08 |
| Promoter (0.12%) | 0.13 | 0.14 | 0.36 | 14 | 1.4 | 2.5e-05 |
| PC-HiC (1.28%) | 0.082 | 0.038 | 0.029 | 3.9 | 0.23 | 2.4e-10 |
| 5kb (0.93%) | -0.033 | 0.036 | 0.36 | 4.5 | 0.17 | 5.4e-10 |
| 100kb (2.43%) | 0.078 | 0.04 | 0.053 | 3.6 | 0.13 | 5.9e-11 |
| BasenjiΔ-published (PPI-enhancer) | | | | | | |
| | $\tau^*$ | se($\tau^*$) | p($\tau^*$) | $E$ | se($E$) | p($E$) |
| ABC (0.14%) | 0.52 | 0.29 | 0.073 | 17 | 1.8 | 1.4e-06 |
| TSS (0.09%) | 0.099 | 0.21 | 0.64 | 19 | 2 | 7.1e-05 |
| Coding (0.05%) | 0.43 | 0.21 | 0.037 | 22 | 2.6 | 2.7e-05 |
| ATAC (0.08%) | 0.19 | 0.18 | 0.28 | 16 | 2.8 | 5.2e-06 |
| eQTL (0.07%) | 0.095 | 0.075 | 0.21 | 8.6 | 1.3 | 3.3e-06 |
| Roadmap (0.23%) | -0.27 | 0.2 | 0.18 | 12 | 1.6 | 3.5e-09 |
| Promoter (0.15%) | 0.13 | 0.14 | 0.38 | 13 | 1.4 | 8.6e-06 |
| PC-HiC (1.59%) | 0.073 | 0.038 | 0.054 | 3.8 | 0.21 | 7.7e-11 |
| 5kb (1.16%) | -0.028 | 0.037 | 0.45 | 4.5 | 0.19 | 2.4e-10 |
| 100kb (2.99%) | 0.083 | 0.038 | 0.031 | 3.6 | 0.13 | 3e-11 |
| DeepBindΔ-published (PPI-enhancer) | | | | | | |
| | $\tau^*$ | se($\tau^*$) | p($\tau^*$) | $E$ | se($E$) | p($E$) |
| ABC (0.34%) | 0.54 | 0.22 | 0.015 | 15 | 1.7 | 1e-06 |
| TSS (0.19%) | -0.1 | 0.19 | 0.58 | 17 | 1.6 | 3.7e-05 |
| Coding (0.14%) | 0.42 | 0.18 | 0.02 | 19 | 1.9 | 1.4e-05 |
| ATAC (0.24%) | 0.44 | 0.17 | 0.0085 | 14 | 2.4 | 8.6e-07 |
| eQTL (0.22%) | 0.1 | 0.066 | 0.12 | 7.2 | 1.1 | 4.3e-06 |
| Roadmap (0.62%) | -0.11 | 0.14 | 0.43 | 11 | 1.4 | 1.2e-09 |
| Promoter (0.38%) | -0.018 | 0.12 | 0.88 | 9.6 | 0.96 | 2.2e-05 |
| PC-HiC (5.2%) | 0.087 | 0.031 | 0.0049 | 3 | 0.15 | 2.9e-10 |
| 5kb (3.8%) | -0.016 | 0.027 | 0.56 | 3.5 | 0.14 | 2.4e-10 |
| 100kb (10.2%) | 0.071 | 0.031 | 0.021 | 2.7 | 0.091 | 8.2e-11 |
| deltaSVMΔ-published (PPI-enhancer) | | | | | | |
| | $\tau^*$ | se($\tau^*$) | p($\tau^*$) | $E$ | se($E$) | p($E$) |
| ABC (0.21%) | 0.34 | 0.21 | 0.11 | 15 | 1.8 | 1.2e-06 |
| TSS (0.12%) | -0.19 | 0.19 | 0.3 | 17 | 1.7 | 9.1e-05 |
| Coding (0.09%) | 0.3 | 0.18 | 0.094 | 19 | 2 | 3.5e-05 |
| ATAC (0.15%) | 0.27 | 0.17 | 0.1 | 13 | 2.3 | 2.9e-06 |
| eQTL (0.14%) | 0.1 | 0.066 | 0.11 | 7.3 | 1.1 | 1.2e-05 |
| Roadmap (0.39%) | -0.16 | 0.14 | 0.26 | 11 | 1.4 | 5.7e-09 |
| Promoter (0.24%) | -0.022 | 0.12 | 0.85 | 9.4 | 1 | 3e-05 |
| PC-HiC (3.27%) | 0.09 | 0.032 | 0.0055 | 3 | 0.15 | 6.8e-10 |
| 5kb (2.41%) | -0.023 | 0.03 | 0.43 | 3.4 | 0.13 | 4.5e-10 |
| 100kb (6.56%) | 0.078 | 0.033 | 0.017 | 2.7 | 0.09 | 1.4e-10 |
| DeepSEADΔ-published (pLI) | | | | | | |
| | $\tau^*$ | se($\tau^*$) | p($\tau^*$) | $E$ | se($E$) | p($E$) |
| ABC (0.17%) | -0.013 | 0.2 | 0.95 | 9.1 | 1.2 | 1.4e-05 |
| TSS (0.22%) | -0.041 | 0.14 | 0.77 | 12 | 1.1 | 8.7e-06 |
| Coding (0.13%) | 0.12 | 0.13 | 0.36 | 11 | 1.2 | 9.5e-06 |
| ATAC (0.09%) | 0.18 | 0.094 | 0.054 | 10 | 1.8 | 4.6e-05 |
| eQTL (0.11%) | 0.074 | 0.073 | 0.31 | 5.9 | 0.92 | 0.00011 |
| Roadmap (0.21%) | -0.3 | 0.34 | 0.37 | 7.7 | 1 | 3.8e-05 |
| Promoter (0.38%) | 0.017 | 0.099 | 0.86 | 6.5 | 0.63 | 1.2e-05 |
| PC-HiC (2.29%) | -0.01 | 0.052 | 0.85 | 3 | 0.12 | 7.4e-10 |
| 5kb (3.25%) | -0.006 | 0.022 | 0.78 | 2.4 | 0.053 | 5.8e-09 |
| 100kb (8.12%) | -0.047 | 0.025 | 0.058 | 1.9 | 0.037 | 3.7e-10 |
| BasenjiΔ-published (pLI) | | | | | | |
| | $\tau^*$ | se($\tau^*$) | p($\tau^*$) | $E$ | se($E$) | p($E$) |
| ABC (0.19%) | -0.14 | 0.17 | 0.42 | 9.2 | 1.1 | 1.7e-05 |
| TSS (0.24%) | -0.081 | 0.14 | 0.56 | 12 | 1.1 | 1.6e-05 |
| Coding (0.16%) | 0.11 | 0.13 | 0.43 | 11 | 1.2 | 6.9e-06 |
| ATAC (0.10%) | 0.15 | 0.091 | 0.11 | 9.5 | 1.6 | 2.8e-05 |
| eQTL (0.13%) | 0.063 | 0.065 | 0.33 | 5.8 | 0.78 | 4.3e-05 |
| Roadmap (0.25%) | -0.42 | 0.35 | 0.22 | 9 | 1.2 | 8.9e-07 |
| Promoter (0.44%) | 0.013 | 0.11 | 0.9 | 6.5 | 0.59 | 6.9e-06 |
| PC-HiC (2.79%) | -0.023 | 0.049 | 0.63 | 2.9 | 0.11 | 1.7e-10 |
| 5kb (3.89%) | -0.011 | 0.021 | 0.6 | 2.4 | 0.05 | 2e-09 |
| 100kb (9.76%) | -0.045 | 0.023 | 0.049 | 1.9 | 0.034 | 8.6e-11 |
| DeepBindΔ-published (pLI) | | | | | | |
| | $\tau^*$ | se($\tau^*$) | p($\tau^*$) | $E$ | se($E$) | p($E$) |
| ABC (0.48%) | -0.11 | 0.15 | 0.5 | 8.1 | 0.89 | 1.7e-05 |
| TSS (0.51%) | -0.21 | 0.14 | 0.14 | 11 | 0.97 | 2.3e-05 |
| Coding (0.45%) | -0.083 | 0.12 | 0.48 | 8.8 | 0.92 | 8.1e-06 |
| ATAC (0.31%) | 0.088 | 0.072 | 0.22 | 7.3 | 1.3 | 3e-05 |
| eQTL (0.42%) | 0.048 | 0.057 | 0.4 | 4.7 | 0.72 | 0.00014 |
| Roadmap (0.70%) | -2 | 0.8 | 0.014 | 7.7 | 0.78 | 3.4e-07 |
| Promoter (1.19%) | -0.058 | 0.075 | 0.44 | 4.9 | 0.58 | 7.8e-05 |
| PC-HiC (9.2%) | -0.021 | 0.042 | 0.61 | 2.3 | 0.079 | 2.4e-10 |
| 5kb (13.9%) | -0.0035 | 0.016 | 0.82 | 1.8 | 0.045 | 1.3e-08 |
| 100kb (28%) | -0.025 | 0.016 | 0.12 | 1.4 | 0.022 | 3.3e-10 |
| deltaSVMΔ-published (pLI) | | | | | | |
| | $\tau^*$ | se($\tau^*$) | p($\tau^*$) | $E$ | se($E$) | p($E$) |
| ABC (0.31%) | -0.14 | 0.15 | 0.35 | 7.9 | 0.94 | 2.3e-05 |
| TSS (0.33%) | -0.14 | 0.14 | 0.33 | 11 | 1 | 2.9e-05 |
| Coding (0.28%) | -0.095 | 0.12 | 0.42 | 8.8 | 0.96 | 1.3e-05 |
| ATAC (0.19%) | 0.073 | 0.076 | 0.34 | 7.1 | 1.2 | 8.2e-05 |
| eQTL (0.27%) | 0.05 | 0.059 | 0.4 | 4.6 | 0.78 | 0.00031 |
| Roadmap (0.45%) | -0.63 | 0.48 | 0.19 | 7.6 | 0.86 | 5.2e-06 |
| Promoter (0.76%) | -0.047 | 0.077 | 0.54 | 4.9 | 0.63 | 0.00013 |
| PC-HiC (5.87%) | -0.015 | 0.044 | 0.73 | 2.3 | 0.087 | 6.4e-10 |
| 5kb (8.78%) | -0.0012 | 0.017 | 0.94 | 1.8 | 0.051 | 2.6e-08 |
| 100kb (22%) | -0.025 | 0.018 | 0.17 | 1.4 | 0.028 | 5.1e-10 |

**Table S24.** Top significant features of Imperio-DeepSEA and Imperio-Basenji models: We report the top 10 chromatin marks (if significant) based on their magnitude of effect size. We consider 4 different models - the Imperio model fitted and evaluated on all genes for DeepSEA (Imperio-DeepSEA) and Basenji (Imperio-Basenji) and the Imperio model fitted and evaluated on PPI-enhancer genes for DeepSEA (Imperio-DeepSEA (PPI-enhancer)) and Basenji (Imperio-Basenji (PPI-enhancer)).

| Imperio model | Top features |
| --- | --- |
| Imperio-DeepSEA | TF::BRF1::HeLa-S3 (TSS), TF::Znf143::HeLa-S3 (Roadmap), TF::ZNF274::HeLa-S3 (TSS), TF::USF2::HepG2 (Roadmap), TF::USF2::HepG2 (TSS), TF::Znf143::GM12878 (Roadmap), TF::Pol2::GM12878 (Roadmap), TF::BRF1::HeLa-S3 (Roadmap), TF::Mxi1::K562 (Roadmap), TF::BHLHE40::K562 (Roadmap) |
| Imperio-Basenji | CAGE:spleen, adult, pool1 (Roadmap), CAGE:spinal cord, adult, donor10252 (TSS), DNASE:SW480 (TSS), HISTONE:H3K9ac neural progenitor cell derived from H9 (TSS), CAGE:Smooth muscle cells - airway, control, donor1 (Roadmap), CAGE:epithelioid sarcoma cell line:HS-ES-1 (Roadmap), DNASE:LNCaP clone FGC treated with 17B-hydroxy-17-methylestra-4,9,11-trien-3-one (Roadmap), HISTONE:H3K9ac keratinocyte female (TSS), HISTONE:H3K27me3 fibroblast of arm male adult (53 years) (TSS), HISTONE:H3K27me3 PC9 (TSS) |
| Imperio-DeepSEA (PPI-enhancer) | TF::Znf143::GM12878 (TSS), TF::ZNF274::K562 (TSS), TF::BRF1::HeLa-S3 (TSS), TF::Rad21::A549 (TSS), TF::USF2::HepG2 (TSS), TF::Znf143::HeLa-S3 (TSS), TF::BHLHE40::K562 (TSS), TF::YY1::NT2-D1 (TSS), forskolin::ERRA::HepG2 (Roadmap), TF::BRCA1::HeLa-S3 (TSS) |
| Imperio-Basenji (PPI-enhancer) | HISTONE:H3K4me3 kidney epithelial cell (TSS), CAGE:spleen, adult, pool1 (Roadmap), CAGE:Macrophage monocyte derived, donor3 (Roadmap), HISTONE:H3K4me2 OCI-LY3 (TSS), CAGE:CD14+ Monocytes, donor2 (Roadmap), CAGE:peripheral neuroectodermal tumor cell line:KU-SN (Roadmap), CAGE:caudate nucleus, adult, donor10252 (Roadmap), CAGE:CD34 cells differentiated to erythrocyte lineage (Roadmap), CAGE:carcinoid cell line:NCI-H1770 (Roadmap), CAGE:immature langerhans cells, donor1 (TSS) |

**Table S25.** Comparison of 5 Imperio models utilizing a single S2G strategy with respect to using all 5 S2G strategies: We perform the Imperio prediction model for DeepSEA and Basenji features corresponding to only one of the 5 S2G strategies, and compare the resulting model fit with that of the full Imperio model corresponding to all 5 S2G strategies. We use two measures of model fit - the *r*^2^ metric and the correlation of predicted expression (corr.pred) with original expression on the genes of a holdout chromosome (chr8).

| S2G | Imperio-DeepSEA |  | Imperio-Basenji |  |
| --- | --- | --- | --- | --- |
| | $r^2$ | corr.pred | $r^2$ | corr.pred |
| ABC | 0.36 | 0.60 | 0.39 | 0.66 |
| Roadmap | 0.33 | 0.63 | 0.37 | 0.68 |
| TSS | 0.59 | 0.69 | 0.61 | 0.71 |
| 5kb | 0.40 | 0.66 | 0.52 | 0.73 |
| 100kb | 0.29 | 0.52 | 0.35 | 0.61 |
| All | 0.66 | 0.72 | 0.69 | 0.76 |

**Table S26.** Proportion of cis-heritability captured by Imperio and ExPecto predictions of gene expression across individuals: Results are averaged across all 22,020 genes for the 2 Imperio models (Imperio-DeepSEA and Imperio-Basenji) and the ExPecto-DeepSEA model.

| Model | Correlation |
| --- | --- |
| Imperio-DeepSEA | 0.82 |
| Imperio-Basenji | 0.79 |
| ExPecto-DeepSEA | 0.75 |

**Table S27.** Correlation of boosted-restricted deep learning allelic-effect annotations restricted using S2G strategies with Imperio annotations: We report the correlation of boosted-restricted deep learning allelic-effect annotations restricted using S2G strategies with Imperio annotations for DeepSEA and Basenji deep learning models.

| restricted S2G | Imperio-DeepSEA | Imperio-Basenji |
| --- | --- | --- |
| BasenjiΔ-boosted × ABC | 0.12 | 0.10 |
| BasenjiΔ-boosted × TSS | 0.15 | 0.16 |
| BasenjiΔ-boosted × Coding | 0.03 | 0.04 |
| BasenjiΔ-boosted × ATAC | 0.05 | 0.07 |
| BasenjiΔ-boosted × eQTL | 0.07 | 0.08 |
| BasenjiΔ-boosted × Roadmap | 0.11 | 0.13 |
| BasenjiΔ-boosted × Promoter | 0.12 | 0.15 |
| BasenjiΔ-boosted × PC-HiC | 0.11 | 0.12 |
| BasenjiΔ-boosted × 5kb | 0.12 | 0.14 |
| BasenjiΔ-boosted × 100kb | 0.11 | 0.13 |
| DeepSEAΔ-boosted × ABC | 0.13 | 0.10 |
| DeepSEAΔ-boosted × TSS | 0.17 | 0.17 |
| DeepSEAΔ-boosted × Coding | 0.03 | 0.05 |
| DeepSEAΔ-boosted × ATAC | 0.05 | 0.06 |
| DeepSEAΔ-boosted × eQTL | 0.07 | 0.07 |
| DeepSEAΔ-boosted × Roadmap | 0.13 | 0.13 |
| DeepSEAΔ-boosted × Promoter | 0.12 | 0.15 |
| DeepSEAΔ-boosted × PC-HiC | 0.12 | 0.11 |
| DeepSEAΔ-boosted × 5kb | 0.12 | 0.12 |
| DeepSEAΔ-boosted × 100kb | 0.11 | 0.10 |

**Table S28.** S-LDSC results for Imperio and ExPecto annotations, conditional on the baseline-LD-deep-S2G-geneset model annotations: Standardized Effect sizes (*τ\**) and Enrichment (E) of Imperio annotations for DeepSEA (Imperio-DeepSEA) and Basenji (Imperio-Basenji) along with a similarly defined ExPecto (ExPecto-DeepSEA) annotation. Results were conditional on 115 baseline-LD-deep-S2G-geneset annotations. Reports are meta-analyzed across 11 blood-related traits.

| | $\tau^*$ | $se(\tau^*)$ | $p(\tau^*)$ | $E$ | $se(E)$ | $p(E)$ |
| --- | --- | --- | --- | --- | --- | --- |
| Imperio-DeepSEA (0.11%) | 0.29 | 0.086 | 0.00075 | 6.5 | 0.47 | 4.9e-06 |
| Imperio-Basenji (0.10%) | 0.48 | 0.096 | 4.9e-07 | 8.2 | 0.61 | 6.1e-07 |
| ExPecto-DeepSEA (0.07%) | 0.13 | 0.063 | 0.045 | 5.5 | 0.53 | 2.1e-05 |

**Table S29.** Standardized enrichment results for Imperio and ExPecto annotations conditional on the baseline-LD-deep model annotations: Standardized Enrichment of Imperio annotations for DeepSEA (Imperio-DeepSEA) and Basenji (Imperio-Basenji) along with a similarly defined ExPecto (ExPecto-DeepSEA) annotations. Results were conditional on 115 baseline-LD-deep-S2G-geneset annotations. Reports are meta-analyzed across 11 blood-related traits.

|  | StdE | se(StdE) | p(StdE) |
| --- | --- | --- | --- |
| Imperio-DeepSEA (0.11%) | 0.045 | 0.0033 | 6.7e-06 |
| Imperio-Basenji (0.10%) | 0.057 | 0.0043 | 7.3e-07 |
| ExPecto-DeepSEA (0.07%) | 0.03 | 0.003 | 5.5e-05 |

**Table S30.** S-LDSC results for gene-set specific Imperio annotations, conditional on the baseline-LD-deep-S2G-geneset model annotations: Standardized Effect sizes (*τ\**) and Enrichment (E) of gene set-specific Imperio annotations corresponding to 2 deep learning models (DeepSEA and Basenji) and 2 gene sets (pLI and PP-enhancer). Results were conditional on 115 baseline-LD-deep-S2G-geneset annotations. Reports are meta-analyzed across 11 blood-related traits.

| | $\tau^*$ | $se(\tau^*)$ | $p(\tau^*)$ | $E$ | $se(E)$ | $p(E)$ |
| --- | --- | --- | --- | --- | --- | --- |
| Imperio-DeepSEA<br>(PPI-enhancer) (0.04%) | 0.78 | 0.16 | 1.5e-06 | 20 | 2.3 | 3.5e-08 |
| Imperio-DeepSEA<br>(pLI) (0.04%) | 0.21 | 0.098 | 0.035 | 8.7 | 1.1 | 1.5e-05 |
| Imperio-Basenji<br>(PPI-enhancer) (0.04%) | 0.53 | 0.14 | 0.0003 | 18 | 2.4 | 4.7e-07 |
| Imperio-Basenji<br>(pLI) (0.05%) | 0.24 | 0.093 | 0.011 | 9 | 1.1 | 6.9e-07 |

**Table S31.** Standardized enrichment results for gene-set specific Imperio annotations conditional on the baseline-LD-deep-S2G-geneset model: Standardized Enrichment of gene set-specific Imperio annotations corresponding to 2 deep learning models (DeepSEA and Basenji) and 2 gene sets (pLI and PP-enhancer) and. Results were conditional on 115 baseline-LD-deep-S2G-geneset annotations. Reports are meta-analyzed across 11 blood-related traits.

|  | StdE | se(StdE) | p(StdE) |
| --- | --- | --- | --- |
| Imperio-DeepSEA<br>(PPI-enhancer) (0.04%) | 0.11 | 0.013 | 3.5e-08 |
| Imperio-DeepSEA<br>(pLI) (0.04%) | 0.044 | 0.0055 | 1.5e-05 |
| Imperio-Basenji<br>(PPI-enhancer) (0.04%) | 0.081 | 0.011 | 4.7e-07 |
| Imperio-Basenji<br>(pLI) (0.05%) | 0.051 | 0.0061 | 6.9e-07 |

**Table S32.** S-LDSC results for joint model of gene set-specific Imperio annotations conditional on the baseline-LD-deep-S2G-geneset model annotations: Joint Standardized Effect sizes (*τ\**) and Enrichment (E) of the 2 marginally significant gene-set specific Imperio annotations, Imperio-DeepSEA (PPI-enhancer) and Imperio-Basenji (PPI-enhancer). Results were conditional on 115 baseline-LD-deep-S2G-geneset annotations. Reports are meta-analyzed across 11 blood-related traits.

| | $\tau^*$ | $se(\tau^*)$ | $p(\tau^*)$ | $E$ | $se(E)$ | $p(E)$ |
| --- | --- | --- | --- | --- | --- | --- |
| Imperio-DeepSEA<br>(PPI-enhancer) (0.04%) | 0.57 | 0.14 | 3.9e-05 | 19 | 2.1 | 5.1e-08 |
| Imperio-Basengi<br>(PPI-enhancer) (0.04%) | 0.16 | 0.14 | 0.26 | 16 | 2.2 | 3.5e-06 |

**Table S33.** S-LDSC results for gene-set specific Imperio annotations, conditional on baseline-LD-deep-S2G-geneset model annotations plus 1 Imperio-Basenji annotation from Figure 3B: Standardized Effect sizes (*τ\**) and Enrichment (E) of gene-set specific Imperio annotations corresponding to 2 deep learning models (DeepSEA and Basenji) and 2 gene-sets (pLI and PP-enhancer). Results were conditional on 115 baseline-LD-deep-S2G-geneset model annotations and 1 Imperio-Basenji annotation from Figure 3B. Reports are meta-analyzed across 11 blood-related traits.

| | $\tau^*$ | $se(\tau^*)$ | $p(\tau^*)$ | $E$ | $se(E)$ | $p(E)$ |
| --- | --- | --- | --- | --- | --- | --- |
| Imperio-DeepSEA<br>(PPI-enhancer) (0.04%) | 0.65 | 0.14 | 7.9e-06 | 19 | 2.3 | 4.2e-08 |
| Imperio-DeepSEA<br>(pLI) (0.04%) | 0.02 | 0.11 | 0.89 | 7.8 | 1.2 | 1e-04 |
| Imperio-Basenji<br>(PPI-enhancer) (0.04%) | 0.42 | 0.14 | 0.002 | 18 | 2 | 5.5e-07 |
| Imperio-Basenji<br>(pLI) (0.05%) | 0.04 | 0.10 | 0.66 | 8.9 | 1.16 | 2.4e-06 |

**Table S34.** S-LDSC results for Imperio+ExPecto annotations, conditional on the baseline-LD-deep-S2G-geneset model annotations: Standardized Effect sizes (*τ\**) and Enrichment (E) of the Δ*_s_* SNP level annotations computed using a combination both ExPecto^4^ and Imperio features for DeepSEA and Basenji models. Results were conditional either on 115 baseline-LD-deep-S2G-geneset annotations or baseline-LD-deep-S2G-geneset plus 1 Imperio-Basenji annotation from Figure 3B. Reports are meta-analyzed across 11 blood-related traits.

| conditional on the baseline-LD-deep-S2G-geneset model |  |  |  |  |  |  |
| --- | --- | --- | --- | --- | --- | --- |
| | $\tau^*$ | $se(\tau^*)$ | $p(\tau^*)$ | $E$ | $se(E)$ | $p(E)$ |
| ExPecto-Imperio-DeepSEA<br>(0.08%) | 0.26 | 0.087 | 0.0023 | 6.1 | 0.42 | 2.5e-06 |
| ExPecto-Imperio-Basenji<br>(0.08%) | 0.41 | 0.093 | 9.7e-06 | 7.1 | 0.5 | 3.6e-07 |
| conditional on the baseline-LD+geneset+Imperio-Basenji-BLD model |  |  |  |  |  |  |
| | $\tau^*$ | $se(\tau^*)$ | $p(\tau^*)$ | $E$ | $se(E)$ | $p(E)$ |
| ExPecto-Imperio-DeepSEA<br>(0.08%) | -0.042 | 0.12 | 0.73 | 5.9 | 0.42 | 3.6e-06 |
| ExPecto-Imperio-Basenji<br>(0.08%) | -0.17 | 0.13 | 0.2 | 7.1 | 0.49 | 3.6e-07 |

**Table S35.** S-LDSC results for partially restricted gene set-specific Imperio annotations defined by restricting only the fitting of feature weights, conditional on the baseline-LD-deep-S2G-geneset model annotations: Standardized Effect sizes (*τ\**) and Enrichment (E) of the intermediate Imperio (Imperio-int1) annotations computed by using all genes for fitting the model and gene sets for computing the expression allelic effects for 2 deep learning models (DeepSEA and Basenji) and 2 gene sets (pLI and PPI-enhancer). Results were conditional either on 115 baseline-LD-deep- S2G-geneset annotations or baseline-LD-deep-S2G-geneset plus 2 significant Imperio annotations from Figure 3. Reports are meta-analyzed across 11 blood-related traits.

| conditional on the baseline-LD-deep-S2G-geneset model |  |  |  |  |  |  |
| --- | --- | --- | --- | --- | --- | --- |
| | $\tau^*$ | $se(\tau^*)$ | $p(\tau^*)$ | $E$ | $se(E)$ | $p(E)$ |
| Imperio-int1-DeepSEA (PPI-enhancer) (0.04%) | 0.62 | 0.13 | 3.9e-06 | 16 | 1.7 | 2.5e-08 |
| Imperio-int1-Basenji (PPI-enhancer) (0.04%) | 0.53 | 0.18 | 0.0025 | 19 | 2.8 | 1.4e-06 |
| Imperio-int1-DeepSEA (pLI) (0.04%) | 0.091 | 0.092 | 0.32 | 6.6 | 0.74 | 6.7e-06 |
| Imperio-int1-Basenji (pLI) (0.04%) | 0.065 | 0.1 | 0.53 | 7.3 | 1.1 | 3.6e-05 |
| conditional on baseline-LD-deep-S2G-geneset plus 2 significant Imperio annotations from Figure 3 |  |  |  |  |  |  |
| | $\tau^*$ | $se(\tau^*)$ | $p(\tau^*)$ | $E$ | $se(E)$ | $p(E)$ |
| Imperio-int1-DeepSEA (PPI-enhancer) (0.04%) | 0.21 | 0.23 | 0.36 | 16 | 1.7 | 3.3e-08 |
| Imperio-int1-Basenji (PPI-enhancer) (0.04%) | 0.077 | 0.14 | 0.63 | 17 | 2.6 | 1.2e-05 |
| Imperio-int1-DeepSEA (pLI) (0.04%) | 0.028 | 0.088 | 0.75 | 7.9 | 0.79 | 8.7e-07 |
| Imperio-int1-Basenji (pLI) (0.04%) | -0.034 | 0.51 | 0.91 | 8.5 | 1.2 | 6.3e-06 |

**Table S36.** S-LDSC results for partially restricted gene set-specific Imperio annotations defined by restricting only the gene expression predictions, conditional on the baseline-LD-deep-S2G-geneset model annotations: Standardized Effect sizes (*τ\**) and Enrichment (E) of the intermediate Imperio (Imperio-int-2) annotations computed by using genes in a geneset for fitting the model and all genes for computing the expression allelic effects for 2 deep learning models (DeepSEA and Basenji) and 2 gene sets (pLI and PPI-enhancer). Results were conditional either on 115 baseline-LD-deep-S2G-geneset annotations or baseline-LD-deep-S2G-geneset plus 2 significant Imperio annotations from Figure 3. Reports are meta-analyzed across 11 blood-related traits.

| conditional on the baseline-LD-deep-S2G-geneset model |  |  |  |  |  |  |
| --- | --- | --- | --- | --- | --- | --- |
| | $\tau^*$ | $se(\tau^*)$ | $p(\tau^*)$ | $E$ | $se(E)$ | $p(E)$ |
| Imperio-int2-DeepSEA (PPI-enhancer) (0.08%) | 0.46 | 0.14 | 0.00099 | 8.6 | 0.85 | 3.7e-06 |
| Imperio-int2-Basenji (PPI-enhancer) (0.10%) | 0.59 | 0.1 | 3.3e-09 | 9.4 | 0.66 | 2.2e-07 |
| Imperio-int2-DeepSEA (pLI) (0.09%) | 0.51 | 0.11 | 2.9e-06 | 8.8 | 0.75 | 5.7e-07 |
| Imperio-int2-Basenji (pLI) (0.10%) | 0.48 | 0.11 | 4.3e-06 | 8.6 | 0.61 | 4.3e-07 |
| conditional on baseline-LD-deep-S2G-geneset plus 2 significant Imperio annotations from Figure 3 |  |  |  |  |  |  |
| | $\tau^*$ | $se(\tau^*)$ | $p(\tau^*)$ | $E$ | $se(E)$ | $p(E)$ |
| Imperio-int2-DeepSEA (PPI-enhancer) (0.08%) | -0.12 | 0.20 | 0.54 | 8.2 | 0.62 | 8.5e-07 |
| Imperio-int2-Basenji (PPI-enhancer) (0.10%) | 0.44 | 0.26 | 0.09 | 9.6 | 0.67 | 1.5e-07 |
| Imperio-int2-DeepSEA (pLI) (0.09%) | 0.023 | 0.24 | 0.89 | 8.5 | 0.62 | 5.7e-07 |
| Imperio-int2-Basenji (pLI) (0.10%) | 0.021 | 0.18 | 0.91 | 8.4 | 0.78 | 9.9e-07 |

**Table S37.**
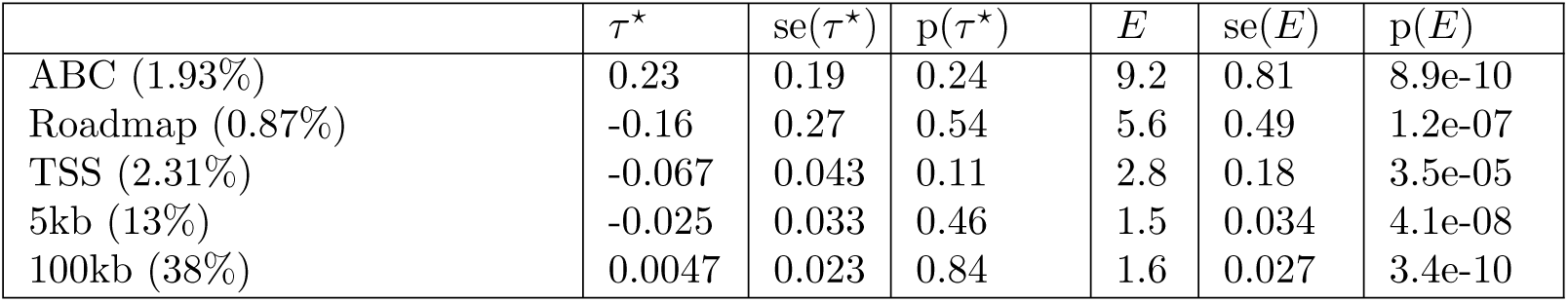
S-LDSC results for annotations defined by the total number of genes linked to each SNP by each S2G strategy, conditional on the baseline- LD-deep-S2G-geneset model annotations: Standardized Effect sizes (*τ\**) and Enrichment (E) of the number of genes (*N_sd_*) linked to each SNP *s* by the S2G strategy *d*. The number of genes was thresholded at 5 and annotations were standardized to probabilistic scale. Results were conditional either on 115 baseline-LD-deep-S2G-geneset annotations. Reports are meta-analyzed across 11 blood-related traits.

| | $\tau^*$ | $se(\tau^*)$ | $p(\tau^*)$ | $E$ | $se(E)$ | $p(E)$ |
| --- | --- | --- | --- | --- | --- | --- |
| ABC (1.93%) | 0.23 | 0.19 | 0.24 | 9.2 | 0.81 | 8.9e-10 |
| Roadmap (0.87%) | -0.16 | 0.27 | 0.54 | 5.6 | 0.49 | 1.2e-07 |
| TSS (2.31%) | -0.067 | 0.043 | 0.11 | 2.8 | 0.18 | 3.5e-05 |
| 5kb (13%) | -0.025 | 0.033 | 0.46 | 1.5 | 0.034 | 4.1e-08 |
| 100kb (38%) | 0.0047 | 0.023 | 0.84 | 1.6 | 0.027 | 3.4e-10 |

**Table S38.**
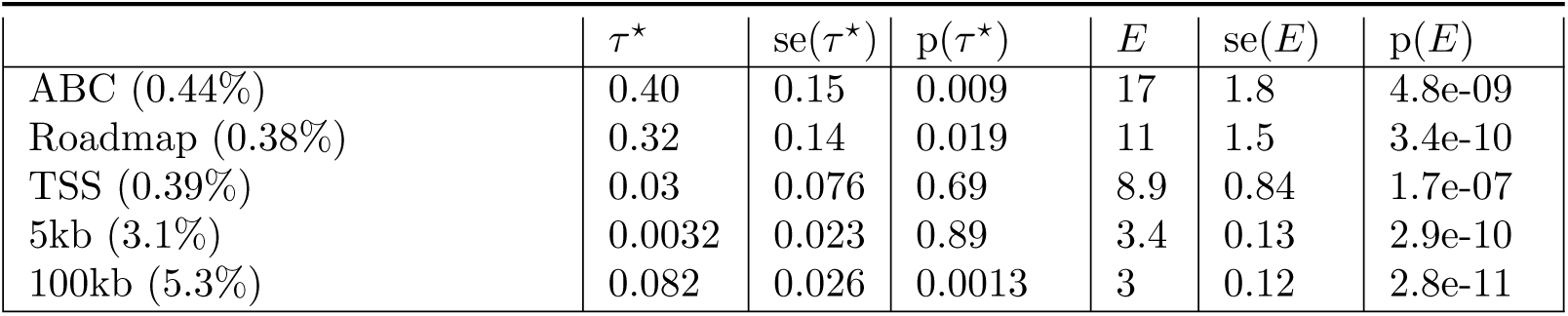
S-LDSC results for annotations defined by the number of PPI- enhancer genes linked to each SNP by each S2G strategy, conditional on the baseline-LD-deep-S2G-geneset model annotations: Standardized Effect sizes (*τ\**) and Enrichment (E) of the number of PPI-enhancer genes linked to each SNP *s* by the S2G strategy *d*. The number of genes was thresholded at 5 and annotations were standardized to probabilistic scale. Results were conditional either on 115 baseline- LD-deep-S2G-geneset annotations. Reports are meta-analyzed across 11 blood-related traits.

**Table S39.** S-LDSC results for Imperio annotations defined using the maximum across genes proximal to the annotated SNPs (instead of the sum), conditional on the baseline-LD-deep-S2G-geneset model annotations plus the two significant annotations from Figure 3B,D: Standardized Effect sizes (*τ\**) and Enrichment (E) of the SNP level annotations computed using the maximum across genes proximal to the annotated SNPs (instead of the sum), conditional on the baseline- LD-deep-S2G-geneset model annotations plus the two significant annotations from Figure 3B,D. Reports are meta-analyzed across 11 blood-related traits.

| | $\tau^*$ | $se(\tau^*)$ | $p(\tau^*)$ | $E$ | $se(E)$ | $p(E)$ |
| --- | --- | --- | --- | --- | --- | --- |
| Imperio-DeepSEA (0.58%) | -0.21 | 0.07 | 0.0035 | 1.4 | 0.15 | 0.058 |
| Imperio-Basengi (0.44%) | 0.13 | 0.056 | 0.023 | 2.6 | 0.14 | 2.3e-06 |
| Imperio-DeepSEA<br>(PPI-enhancer) (0.34%) | -0.11 | 0.055 | 0.045 | 1.6 | 0.17 | 0.0095 |
| Imperio-Basengi<br>(PPI-enhancer) (0.34%) | 0.17 | 0.082 | 0.036 | 2.9 | 0.16 | 5.5e-07 |

**Table S40.**
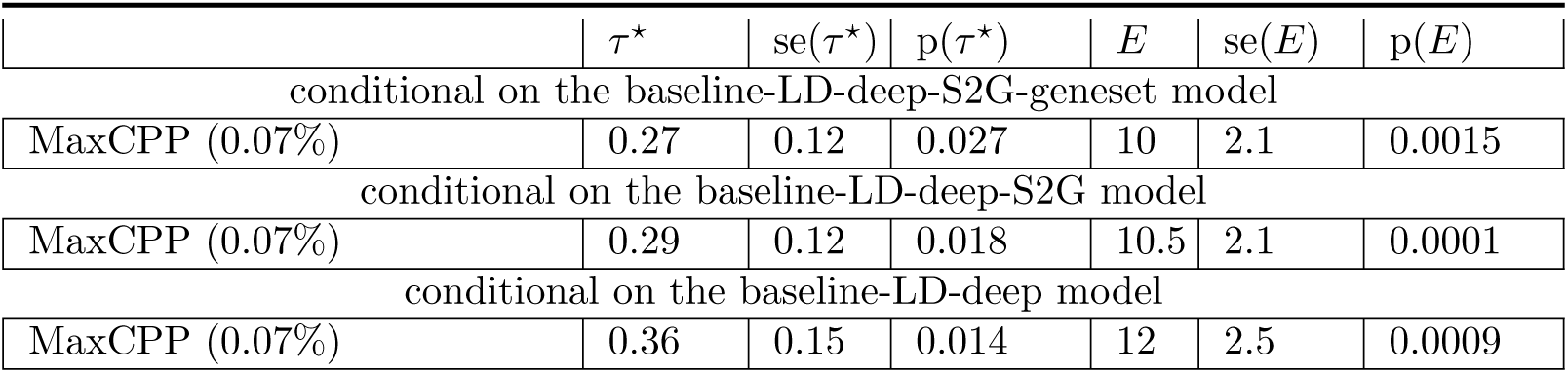
S-LDSC results for Whole blood MaxCPP annotations conditional on different baseline models: Standardized Effect sizes (*τ\**) and Enrichment (E) of Whole blood MaxCPP (MaxCPP) annotations. Results were conditional on either 115 baseline-LD-deep-geneset, or 107 baseline-LD-deep-S2G, or 100 baseline-LD-deep annotations. Reports are meta-analyzed across 11 blood-related traits.

**Table S41.** S-LDSC results for combined joint model: Standardized Effect sizes (*τ\**) and Enrichment (E) in a joint model comprising of significant SNP annotations from Figure 1, Figure 2 and Figure 3. Only results for the 3 jointly Bonferroni significant annotations are reported. The results were conditioned on 115 baseline-LD-deep-S2G- geneset annotations. Reports are meta-analyzed across 11 blood-related traits.

| | $\tau^*$ | $se(\tau^*)$ | $p(\tau^*)$ | $E$ | $se(E)$ | $p(E)$ |
| --- | --- | --- | --- | --- | --- | --- |
| BasenjiΔ-boosted × TSS<br>(0.9%) | 1.1 | 0.29 | 0.0001 | 16 | 1.8 | 8.3e-07 |
| Imperio-Basenji<br>(0.10%) | 0.33 | 0.09 | 0.0001 | 8.2 | 0.61 | 6.4e-07 |
| Imperio-DeepSEA<br>(PPI-enhancer) (0.04%) | 0.67 | 0.15 | 5e-06 | 20 | 2.3 | 3.9e-08 |

**Table S42.** Δlogl_SS_ results for the the combined joint model and other heritability models. We report Δlog*l_SS_* derived from the log*l*_SS_ metric, proposed in ref.^39^, for the different heritability models studied in this paper: baseline-LD, baseline-LD-deep, baseline-LD-deep-S2G, baseline-LD-S2G-geneset and combined joint model in Figure 4 (Table S2). We compute Δlog*l_SS_* as the difference in log*l*_SS_ for each model with respect to s baselineLD-no-funct model with 17 annotations that include no functional annotations^37, 39^. We also report the percentage increase in Δlog*l_SS_* for each model over the baseline-LD model. We do not report AIC as the number of annotations are not too different to alter conclusions based on just the log*l*_SS_. We report three summary Δlog*l*_SS_ results - one averaged across 30 UK Biobank traits^37^ (All), one averaged across 6 blood-related traits from UK Biobank (Blood) and one averaged across the other 24 non blood related traits from UK Biobank (Non-blood) (Table S43).

| Model | $\Delta\log l_{SS}$<br>(All) | % incr.<br>over<br>baseline-LD<br>(All) | pval<br>(All) | $\Delta\log l_{SS}$<br>(Blood) | % incr.<br>over<br>baseline-LD<br>(Blood) | pval<br>(Blood) | $\Delta\log l_{SS}$<br>(Non-blood) | % incr.<br>over<br>baseline-LD<br>(Non-blood) | pval<br>(Non-blood) |
| --- | --- | --- | --- | --- | --- | --- | --- | --- | --- |
| baseline-LD<br>(n=86) | 1379 | 0 | - | 2668 | 0 | - | 1121 | 0 | - |
| baseline-LD-deep<br>(n=100) | 1501 | 8.8% | 5e-44 | 2997 | 12% | 2e-131 | 1179 | 5.1% | 4e-18 |
| baseline-LD-deep-S2G<br>(n=107) | 1542 | 12% | 1e-56 | 3102 | 16% | 3e-170 | 1218 | 8.6% | 5e-30 |
| baseline-LD-deep-S2G<br>-geneset (n=115) | 1582 | 14.7% | 2e-66 | 3142 | 17.8% | 8e-181 | 1259 | 12% | 1e-40 |
| Combined joint model<br>(Figure 4) n=118) | 1603 | 16.2% | 7e-75 | 3209 | 20.3% | 8e-207 | 1280 | 14.2% | 8e-49 |

**Table S43.** List of UKBiobank traits used for log*l*_SS_ calculations. The list consists of 6 blood-related traits and 24 non blood-related traits.

| Trait | Category | N |
| --- | --- | --- |
| disease AID Sure | Blood | 459324 |
| blood Eosinophil count | Blood | 439938 |
| Blood Platelet count | Blood | 444382 |
| Blood Red count | Blood | 445174 |
| Blood RBC distr. width | Blood | 442700 |
| blood White count | Blood | 444502 |
| reproduction Menarche Age | Non-blood | 242278 |
| reproduction Menopause Age | Non-blood | 143025 |
| Body balding | Non-blood | 208336 |
| Body BMIz | Non-blood | 457824 |
| cov EDU Years | Non-blood | 454813 |
| disease Dermatology | Non-blood | 459324 |
| disease Allergy Eczema | Non-blood | 458699 |
| lung FVCzSmoke | Non-blood | 371949 |
| lung FEV1FVCzSmoke | Non-blood | 371949 |
| pigment Hair | Non-blood | 452720 |
| bmd Heel Tscorez | Non-blood | 445921 |
| body Heightz | Non-blood | 458303 |
| disease Hi-chol-self-rep | Non-blood | 459324 |
| disease Hypothyroidism self rep | Non-blood | 459324 |
| Other Morning-person | Non-blood | 41050 |
| Mental Neuroticism | Non-blood | 372066 |
| disease Respiratory ENT | Non-blood | 459324 |
| pigment Skin | Non-blood | 453609 |
| cov Smoking Status | Non-blood | 457683 |
| pigment Sunburn | Non-blood | 344229 |
| bp SystolicadjMedz | Non-blood | 422771 |
| pigment Tanning | Non-blood | 449984 |
| disease T2D | Non-blood | 459324 |
| body WHRadjBMIz | Non-blood | 458417 |

## Supplementary Figures

**Figure S1.**
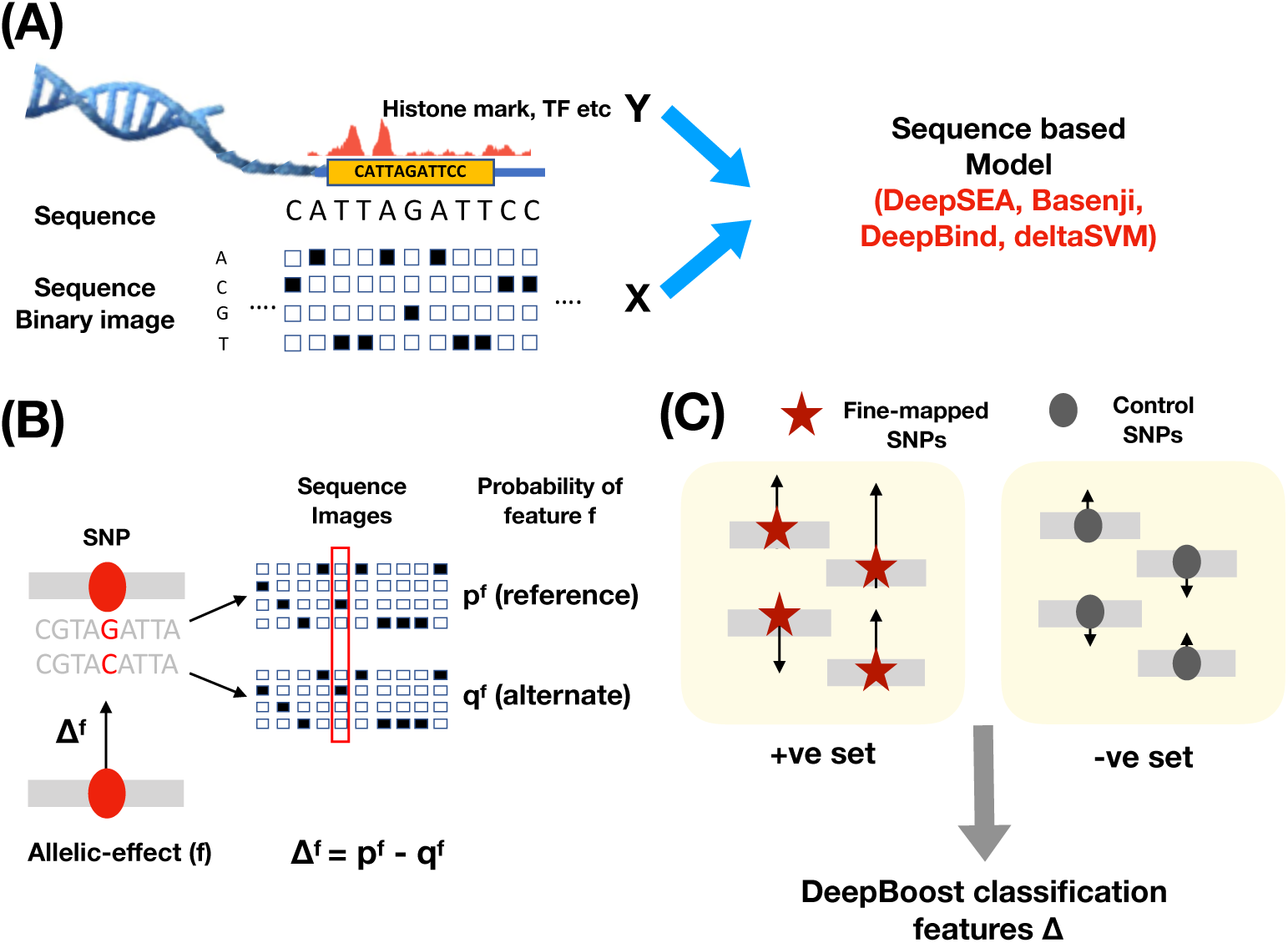
Illustration of the DeepBoost model: (A) An overview of a sequence based genomic deep learning model like DeepSEA and Basenji, that trains on sequence images for a region and the chromatin features around that region using a deep Convolutional Neural Net (CNN) model. (B) Illustration of how the alelic effect annotation for a particular feature *f* is computed at a SNP *s*. The number of features *f* is 2,002 for the DeepSEA model, 4,229 for the Basenji model, 927 for Deepbind and 1329 for the deltaSVM model used. The length of the vertical arrow at the SNP site denotes the magnitude of the allelic effect and its direction represents the sign (up and down for positive and negative allelic effect respectively). (C) Illustration of the DeepBoost classification model where we classify positive set of fine-mapped SNPs from the negative set of matched controls using the allelic effect features.

**Figure S2.**
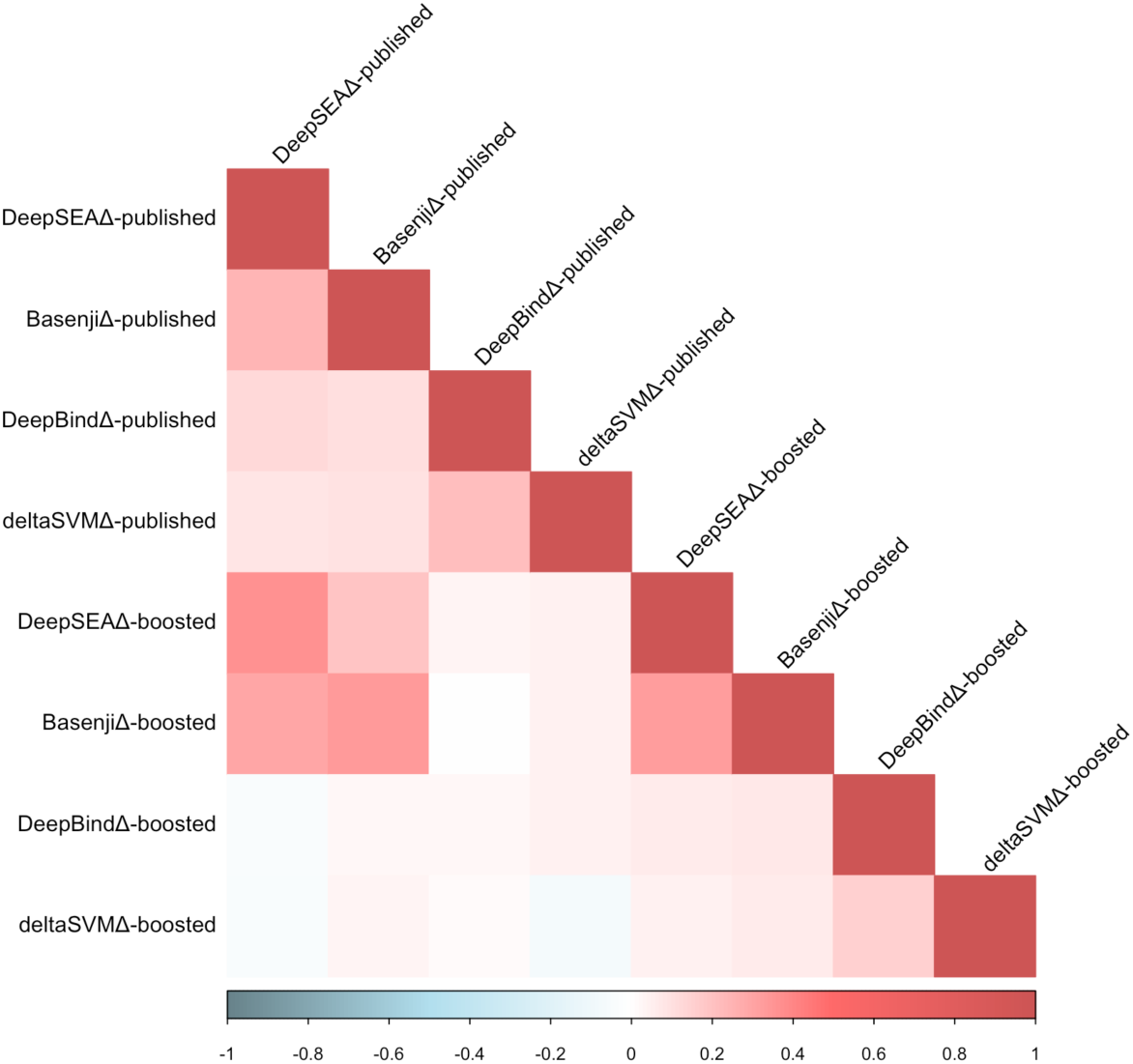
Correlations between published and boosted allelic-effect annotations. Correlation matrix of boosted and published allelic-effect annotations for DeepSEA, Basenji, DeepBind and deltaSVM models. We observed mildly positive correlations between published and boosted annotations for the same model (average *r*=0.16), and we observed weakly positive correlations across all annotations (*r*=0.10).

**Figure S3.**
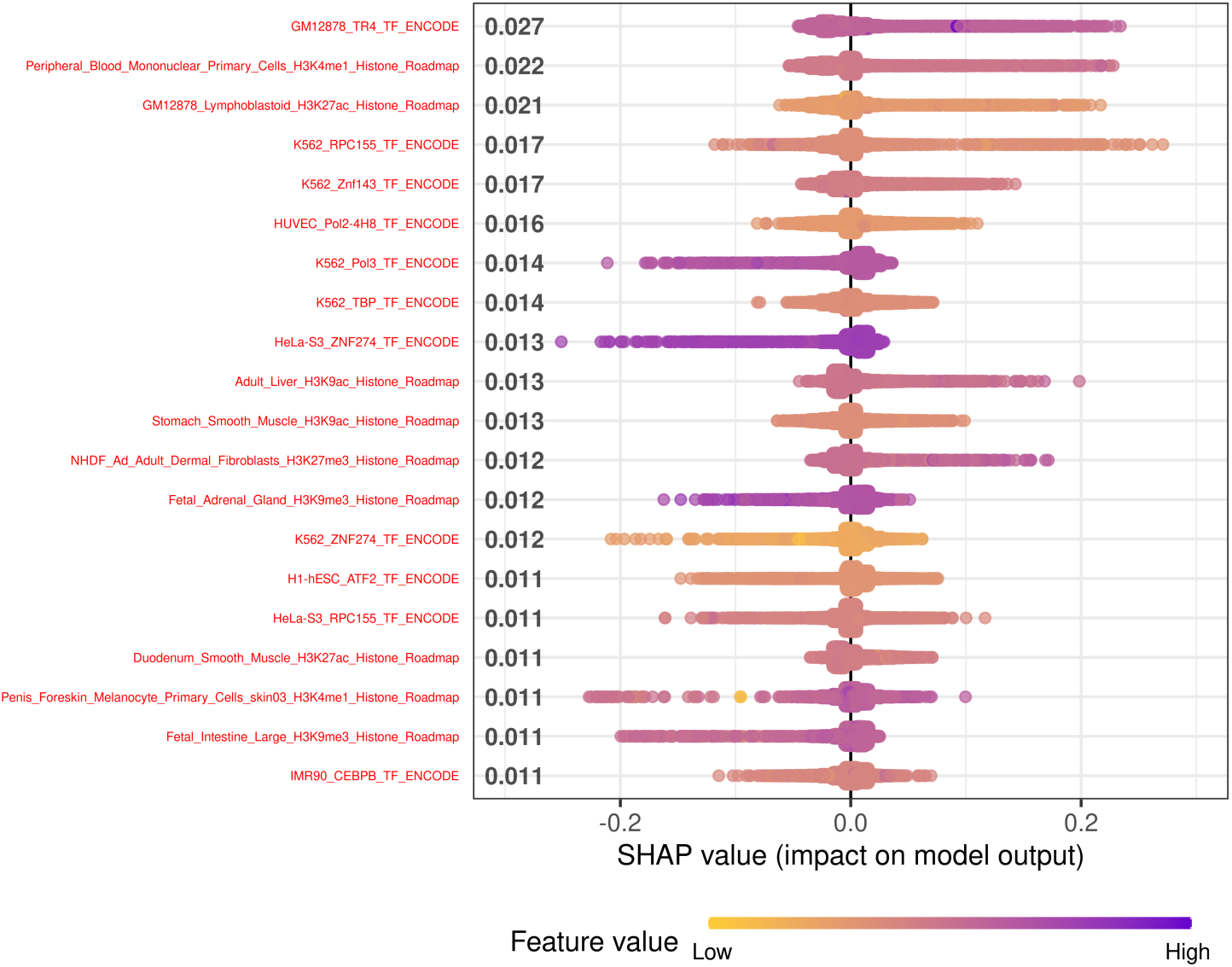
Feature importance of boosted annotations for DeepBoost using the DeepSEA model. We applied SHAP^42^ to assess which deep learning features were most important for the prediction of boosted annotations using the DeepSEA (Methods). We report the top 20 features with signed SHAP scored ordered from top to bottom based on importance as in ref^37^.

**Figure S4.**
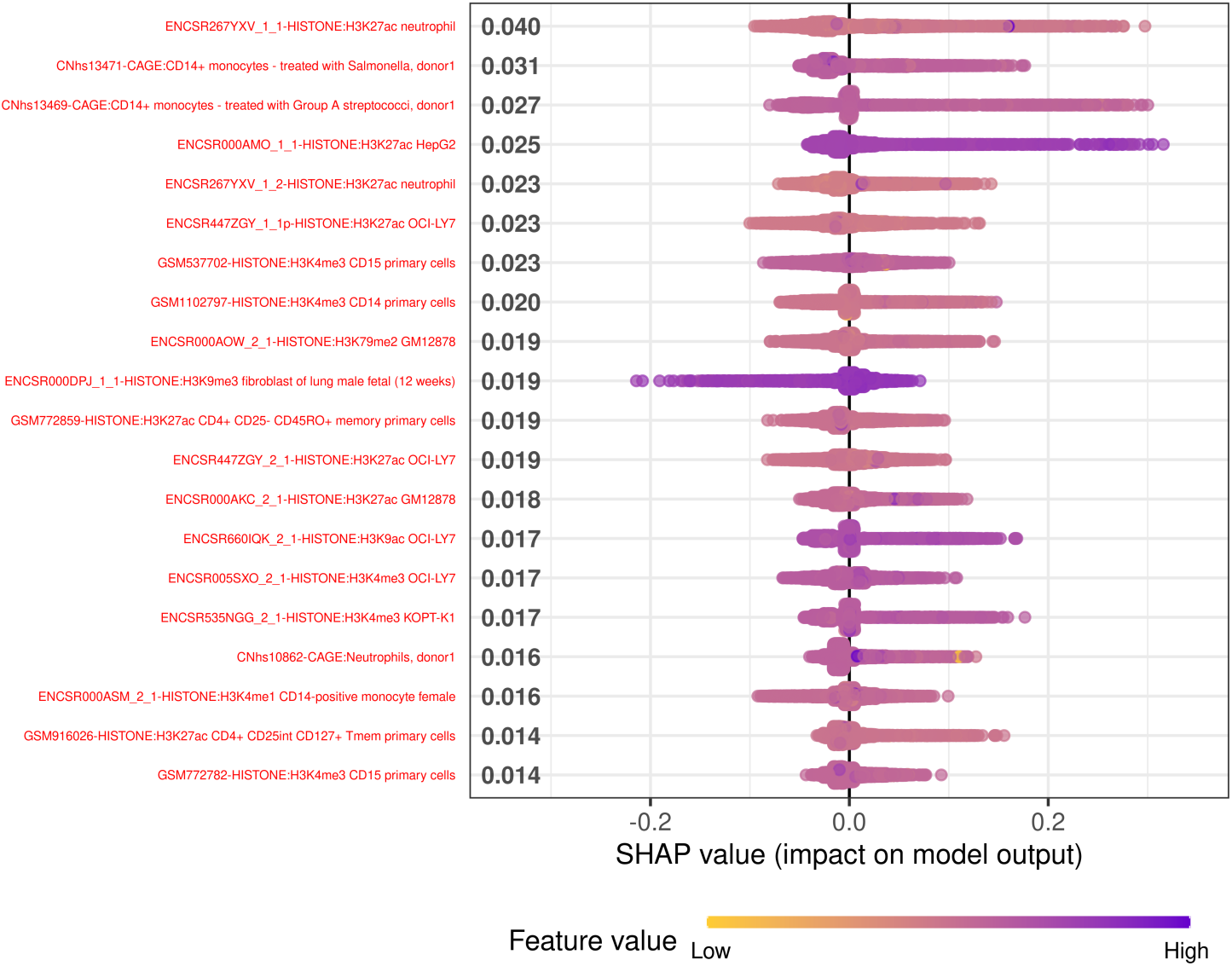
Feature importance of boosted annotations for DeepBoost using the Basenji model. We applied SHAP^42^ to assess which deep learning features were most important for the prediction of boosted annotations using the Basenji (Methods). We report the top 20 features with signed SHAP scored ordered from top to bottom based on importance as in ref^37^.

**Figure S5.**
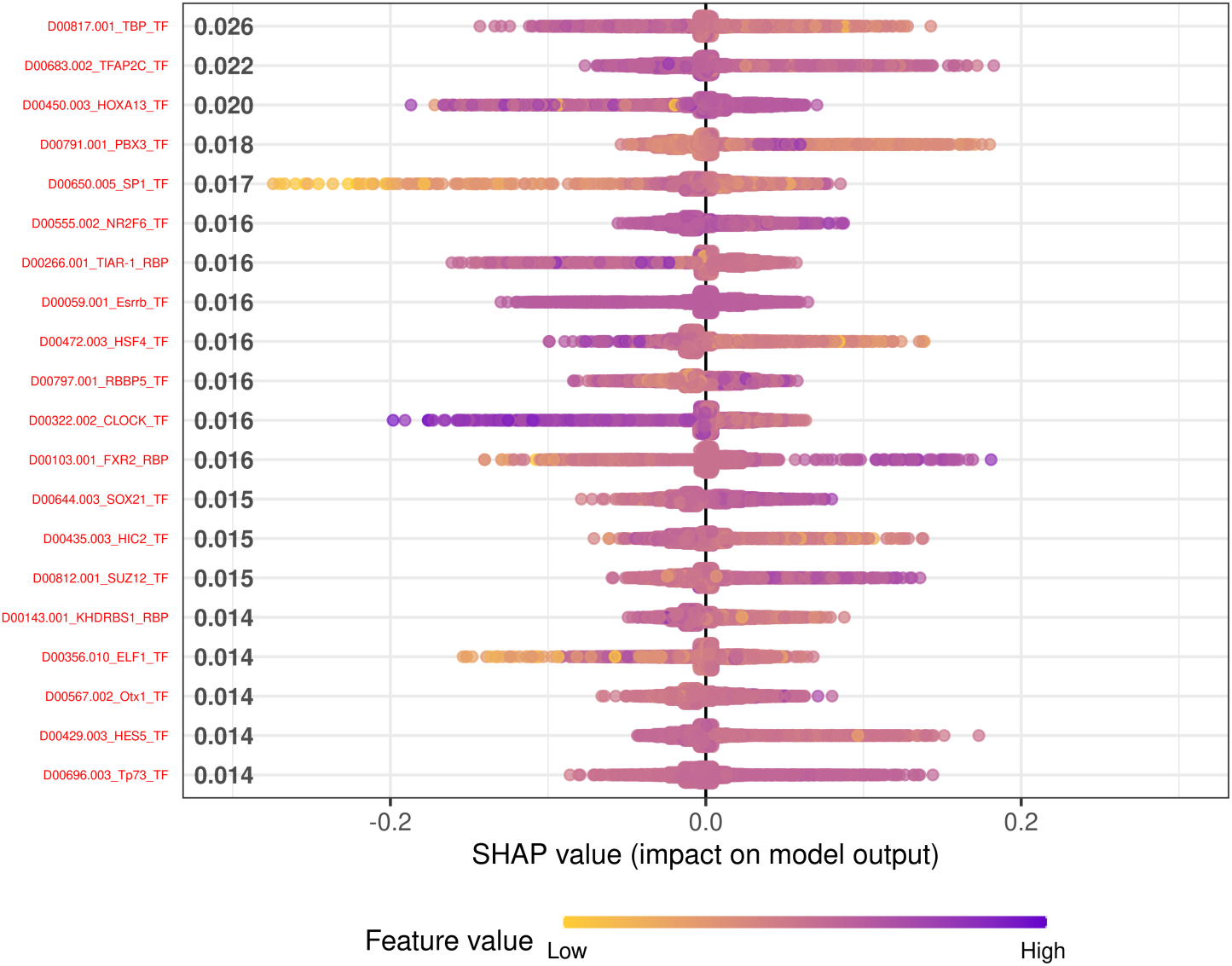
Feature importance of boosted annotations for DeepBoost using the DeepBind model. We applied SHAP^42^ to assess which deep learning features were most important for the prediction of boosted annotations using the DeepBind method (Methods). We report the top 20 features with signed SHAP scored ordered from top to bottom based on importance as in ref^37^.

**Figure S6.**
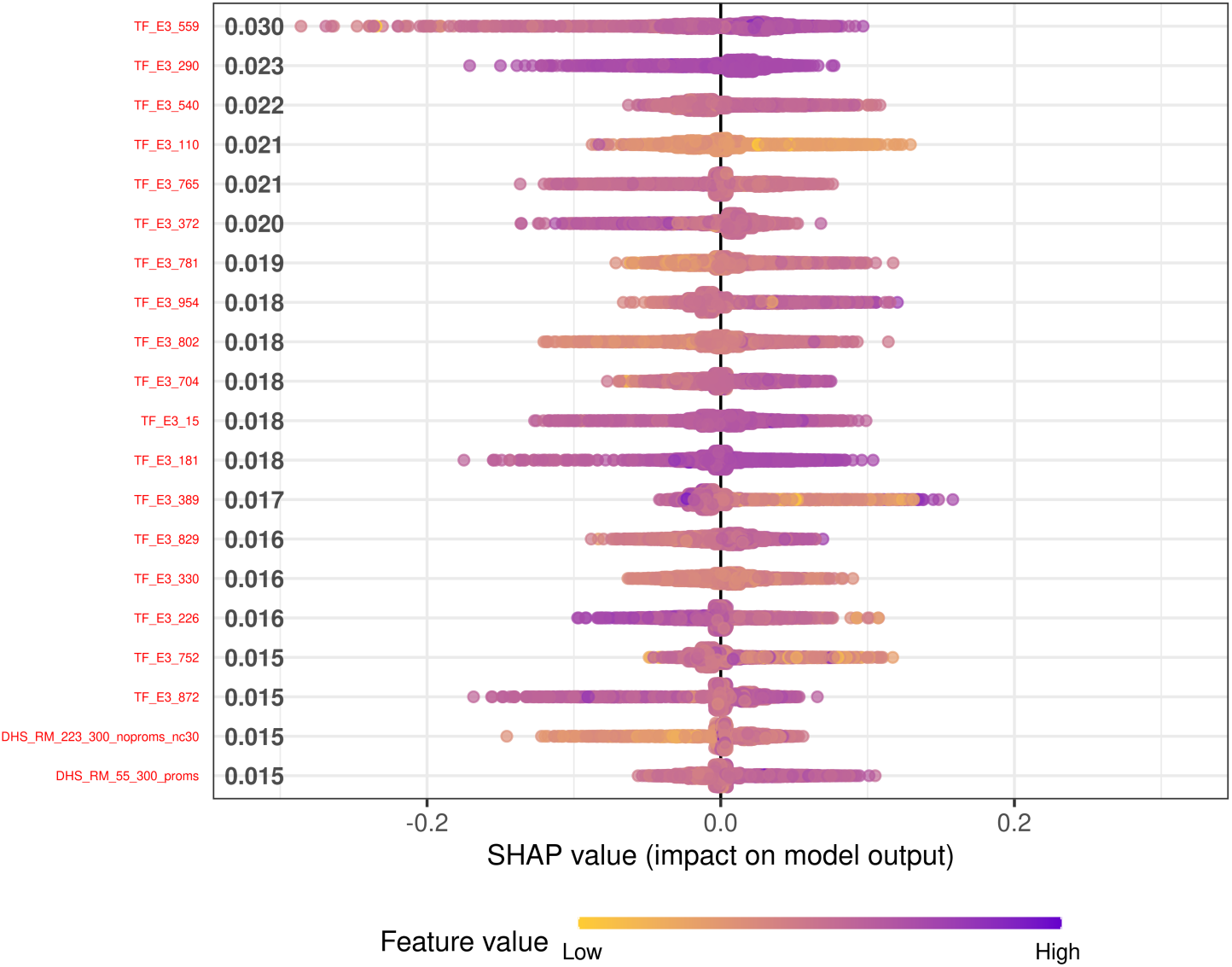
Feature importance of boosted annotations for DeepBoost using the deltaSVM model. We applied SHAP^42^ to assess which deep learning features were most important for the prediction of boosted annotations using the deltaSVM method (Methods). We report the top 20 features with signed SHAP scored ordered from top to bottom based on importance as in ref^37^. **TF E3 559:** eGFP-ZNF507 ChIP-seq on human K562 genetically modified using stable transfection; **TF E3 290:** lung embryo (67 days), **TF E3 540:** ARID1B ChIP-seq on human K562, **TF E3 110:** NT2/D1, **TF E3 765**: EGR1 ChIP-seq on human K562, **TF E3 372**: HAIB ChIP TAF1 in MCF-7, **TF E3 781**: LEF1 ChIP-seq on human K562, **TF E3 954**: MNT ChIP-seq on human MCF-7, **TF E3 802**: CBFA2T3 ChIP-seq on human K562, **TF E3 704**: L3MBTL2 ChIP-seq on human K562, **TF E3 15**: NFE2L2 ChIP-seq on human A549, **TF E3 181**: eGFP-ZNF394 ChIP-seq on human HEK293 genetically modified using site-specific recombination originated from HEK293eGFP-ZNF394 ChIP-seq on human HEK293 genetically modified using site-specific recombination originated from HEK293, **TF E3 389**: HNRNPLL ChIP-seq on human HepG2, **TF E3 829**: eGFP-HINFP ChIP-seq on human K562 genetically modified using stable transfection, **TF E3 330**: CEBPB ChIP-seq protocol v042211.1 on human K562, **TF E3 226**: eGFP-ZSCAN4 ChIP-seq on human HEK293 genetically modified using site-specific recombination originated from HEK293, **TF E3 752**: HNRNPL ChIP-seq on human K562, **TF E3 872**: CUX1 ChIP-seq on human MCF-7.

**Figure S7.**
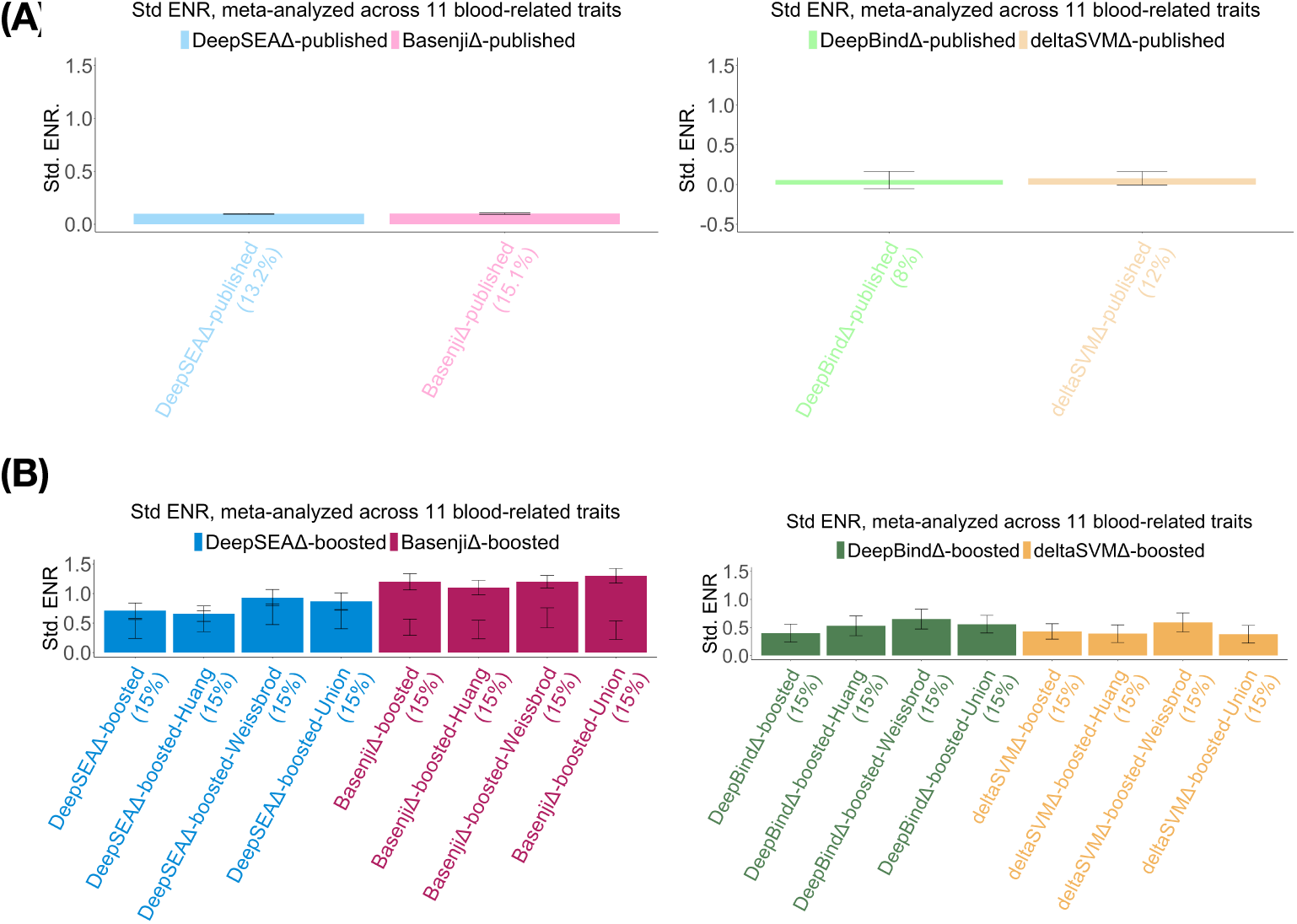
Standardized enrichment of SNP annotations for published and boosted deep learning allelic-effect annotations. Barplot representing standardized enrichment metric, as proposed in ref.^86^, for (A) 4 published DeepSEA, Basenji, DeepBind and deltaSVM allelic-effect annotations and (B) 16 boosted annotations for DeepSEA, Basenji, DeepBind and deltaSVM models, using 3 sets of fine-mapped SNPs and their union. All results are conditional on the baseline-LD-deep model annotations.

**Figure S8.**
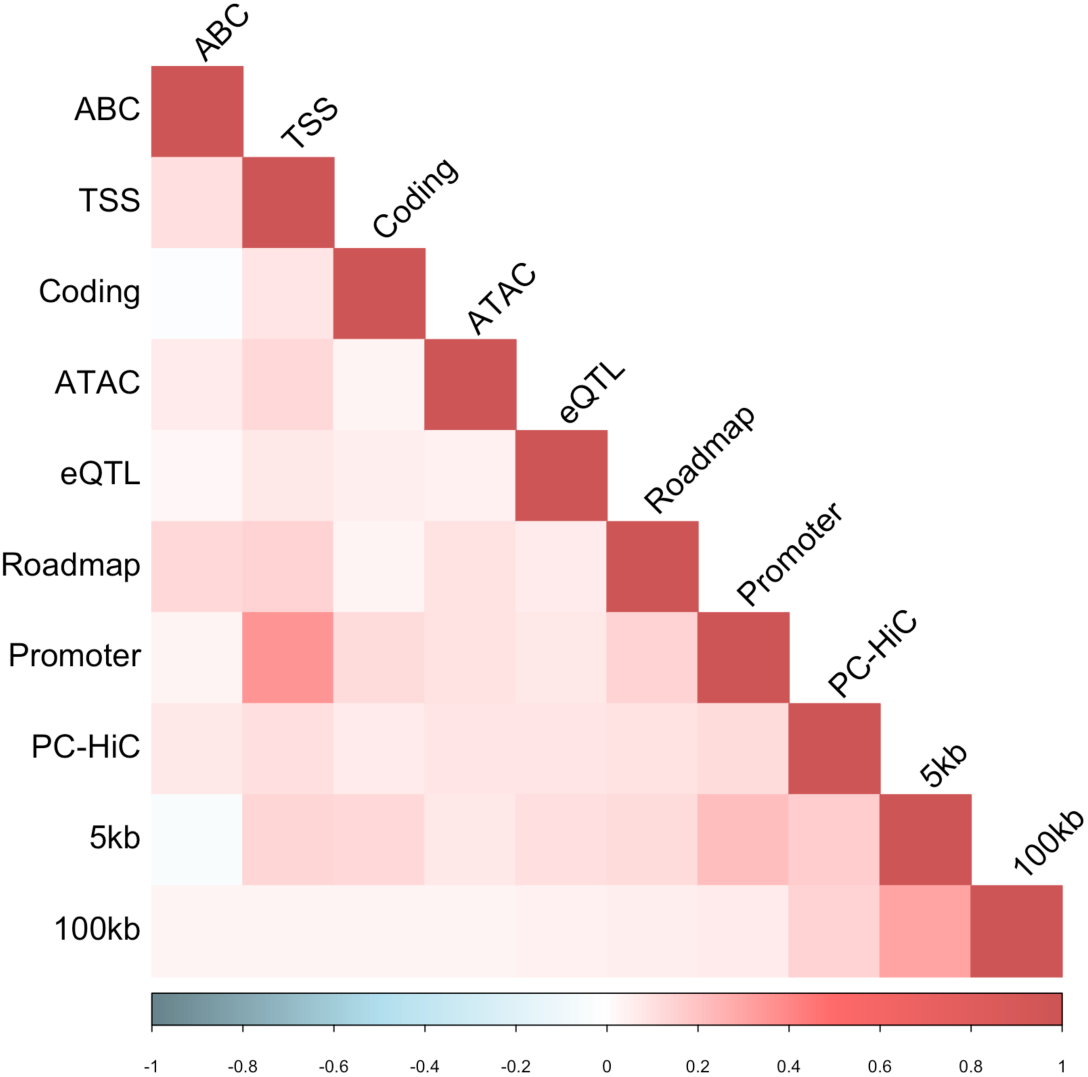
Correlation between S2G annotations. Correlation matrix of S2G annotations derived from all 10 SNP-to-gene (S2G) linking strategies (Table 1), as defined by the sets of SNPs linked to all genes.

**Figure S9.**
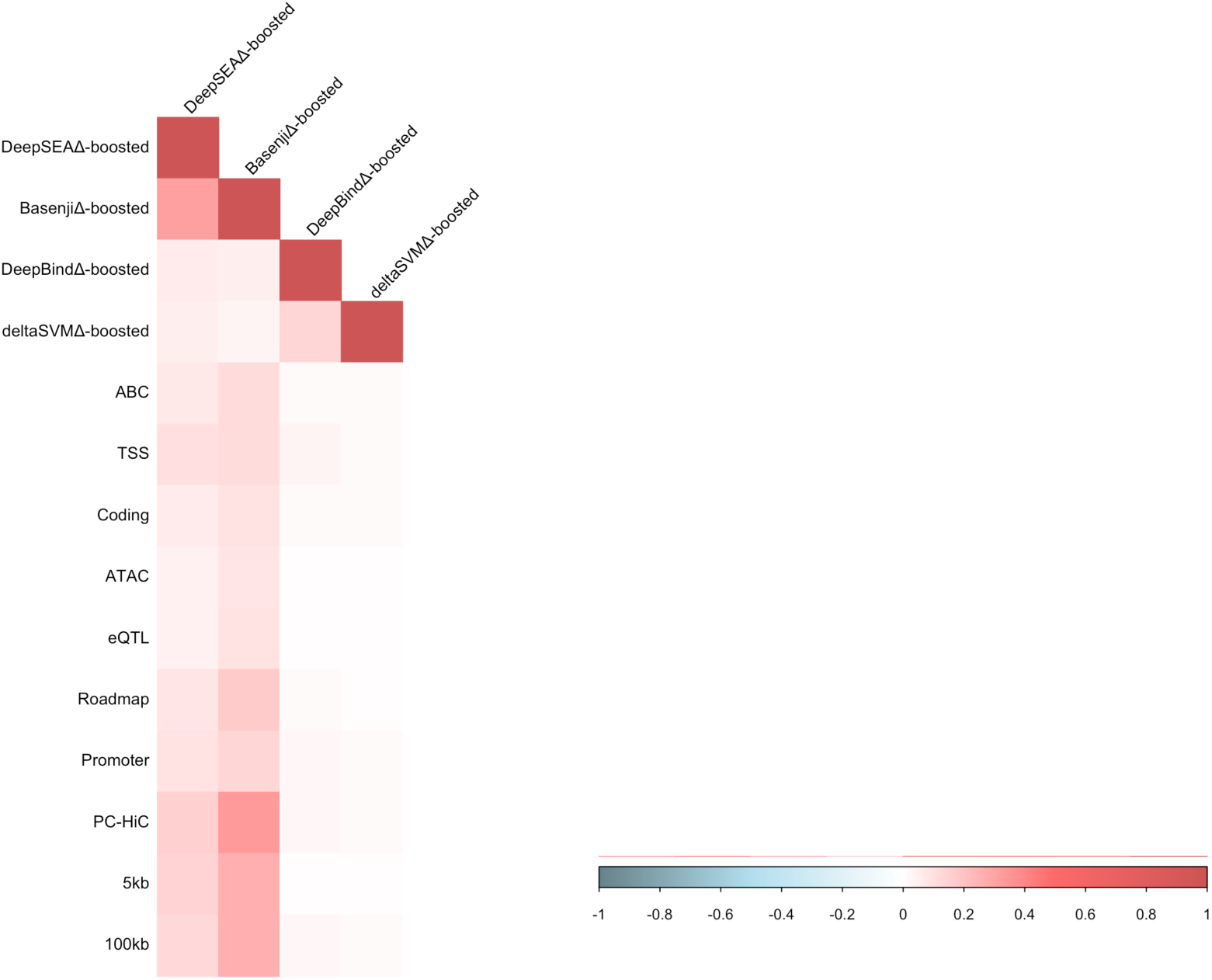
Correlation between boosted allelic-effect annotations and S2G annotations. Correlation matrix of 4 boosted allelic effect annotations, DeepSEAΔ-boosted, BasenjiΔ-boosted, DeepBindΔ-boosted and deltaSVMΔ-boosted, and 10 S2G annotations. The correlations range from weakly positive to moderately positive.

**Figure S10.**
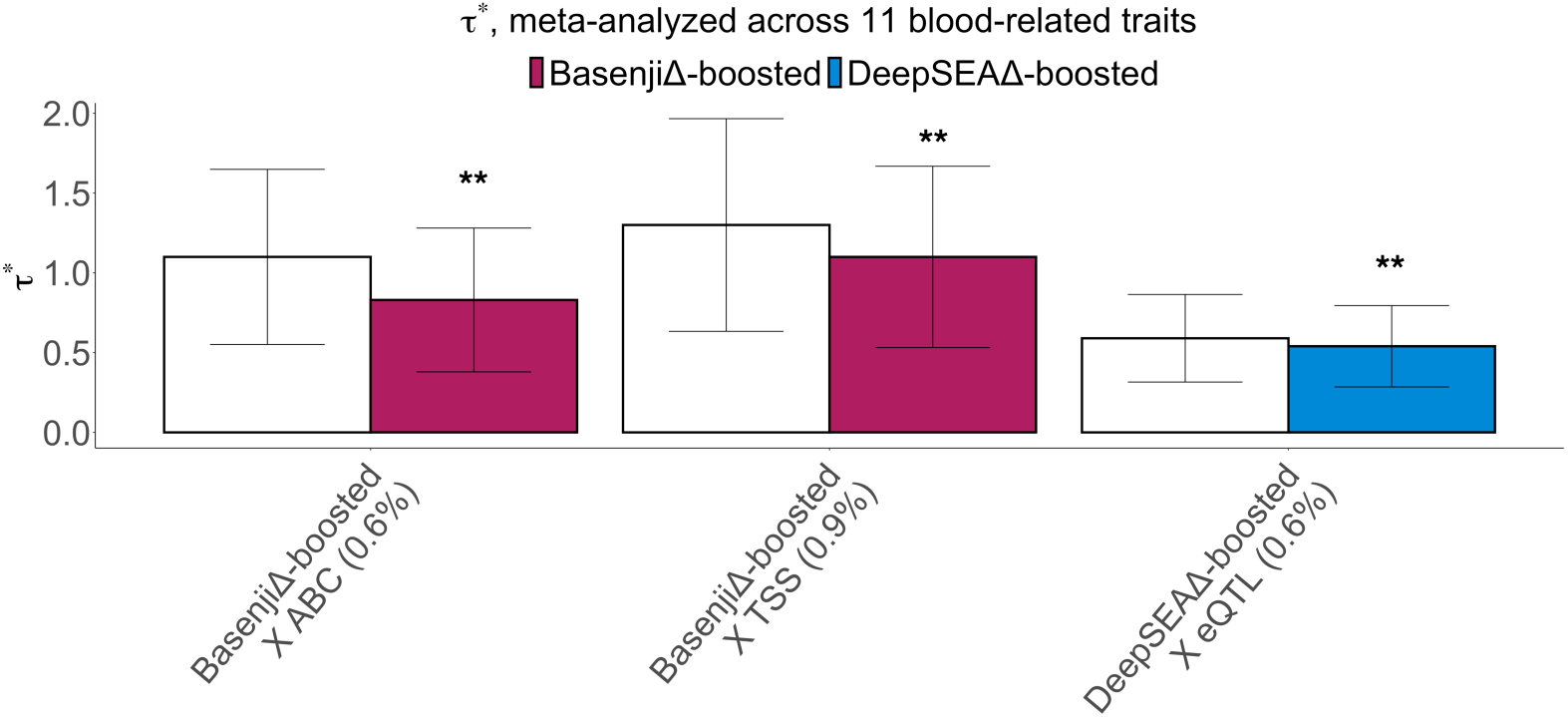
S-LDSC results for joint model of published-restricted and boosted-restricted deep learning allelic-effect annotations restricted using S2G strategies, conditional on the baseline-LD-deep-S2G model annotations. Standardized effect size (*τ\**) conditional on baseline-LD-deep-S2G and other significant restricted S2G annotations (right column, shading) compared to the effect size from Figure 1 Panel B right panel (left column, white). Results are meta-analyzed across 11 blood-related traits. ** denotes *P <* 0.05*/*174. Error bars denote 95% confidence intervals. Numerical results are reported in Table S14.

**Figure S11.**
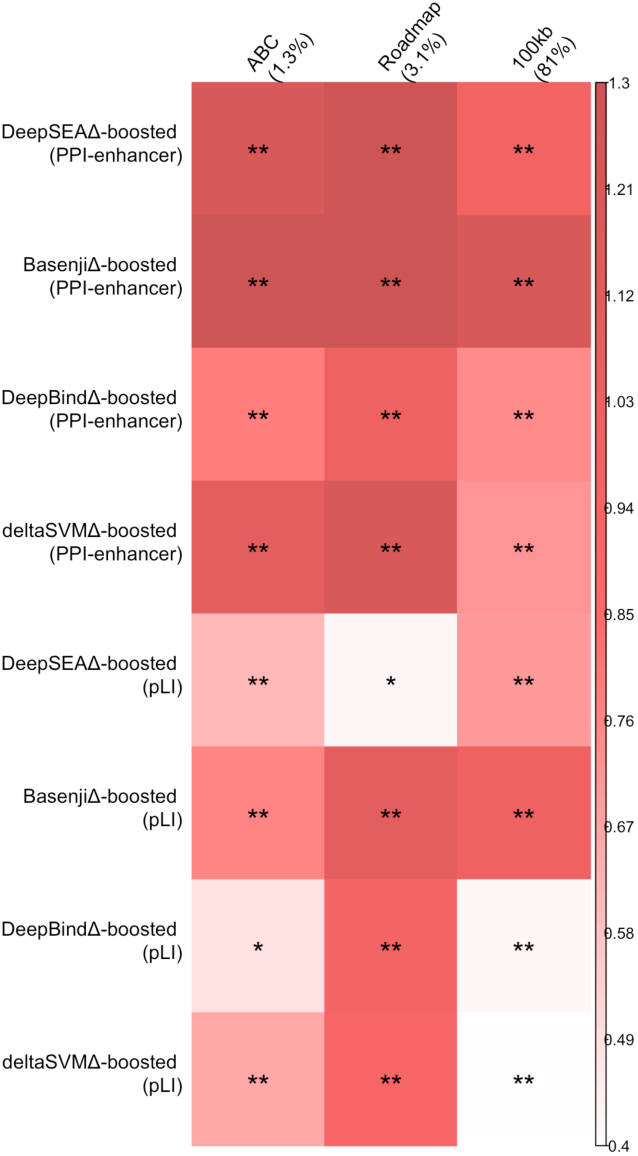
Standardized enrichment of gene set-specific boosted-restricted annotations, conditional on baseline-LD-deep-S2G-geneset model annotations. Standardized enrichment metric, as proposed in ref.^86^, for 80 SNP annotations corresponding to 2 gene scores (PPI-enhancer^38^, pLI^33^) with 10 S2G annotations prioritized by 4 boosted allelic-effect annotations (DeepSEAΔ-boosted, BasenjiΔ-boosted, DeepBindΔ-boosted and deltaSVMΔ- boosted). Results only shown for those allelic-effect models and S2G strategies that show Bonferroni significance. ** denotes *P <* 0.05*/*174. Error bars denote 95% confidence intervals. Numerical results are reported in Table S17.

**Figure S12.**
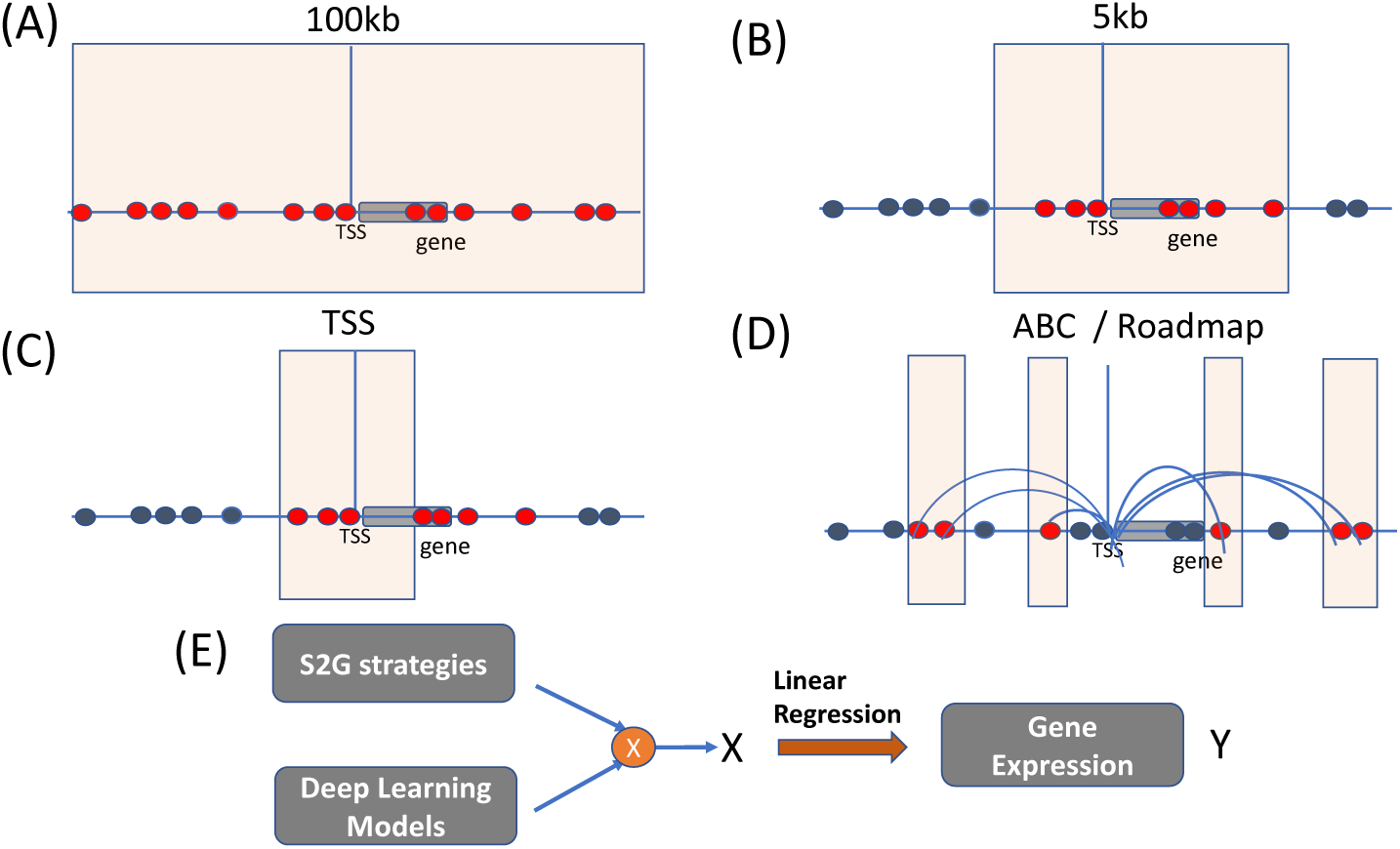
Illustration of the Imperio model: A schematic representation of the different S2G straategie used in the Imperio model : (A) 100kb, (B) 5kb, (C) TSS and (D) ABC or Roadmap. (E) Illustration of how the deep learning variant level or allelic effect annotations are combined with these S2G strategies to generate the featues which are used as predictors in a regression model with GTEx Whole blood expression (log CPM) used as response.

**Figure S13.**
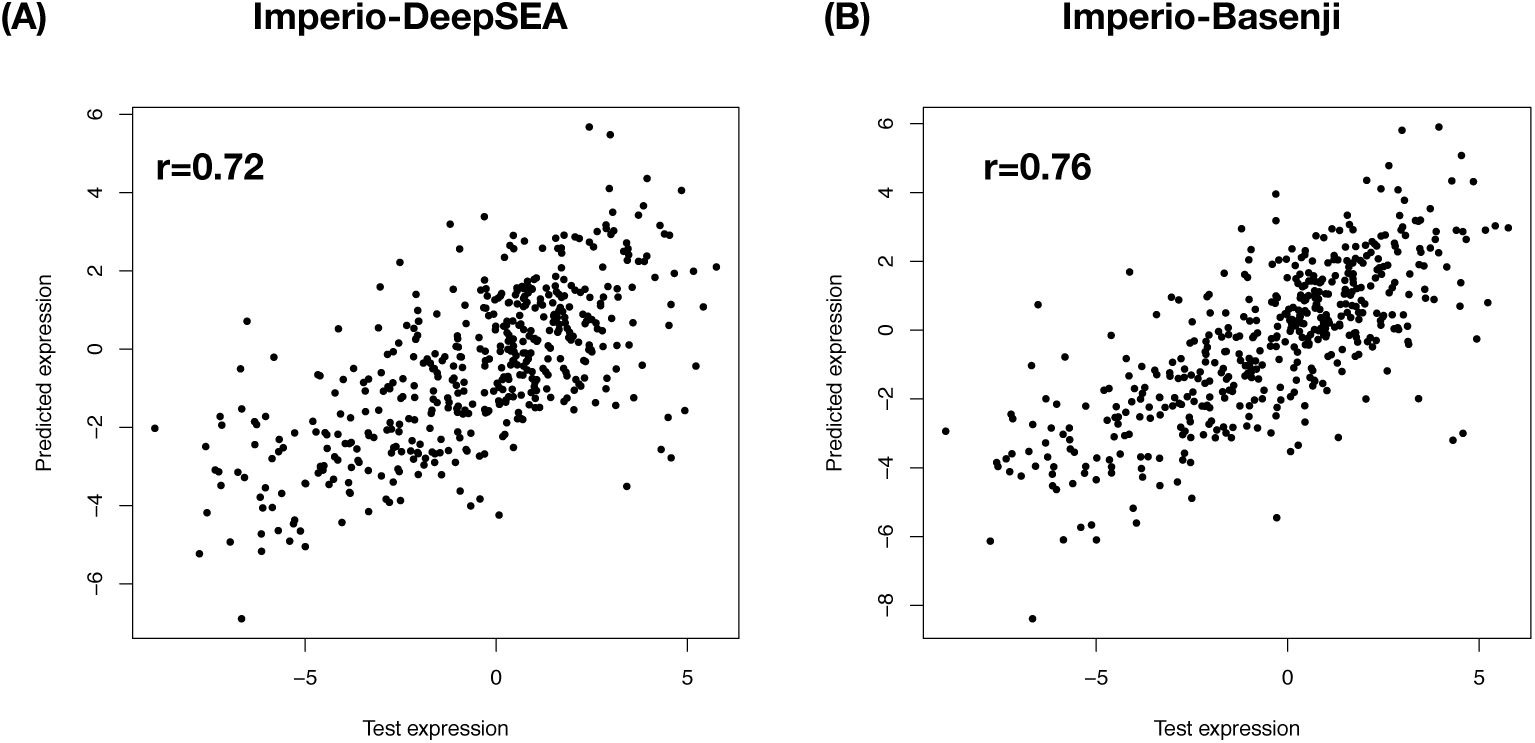
Accuracy of Imperio in predicting gene expression across genes on chromosome 8. For both Imperio-DeepSEA and Imperio-Basenji, we plot predicted expression vs. observed log RPKM expression, for 990 genes on chromosome 8.

**Figure S14.**
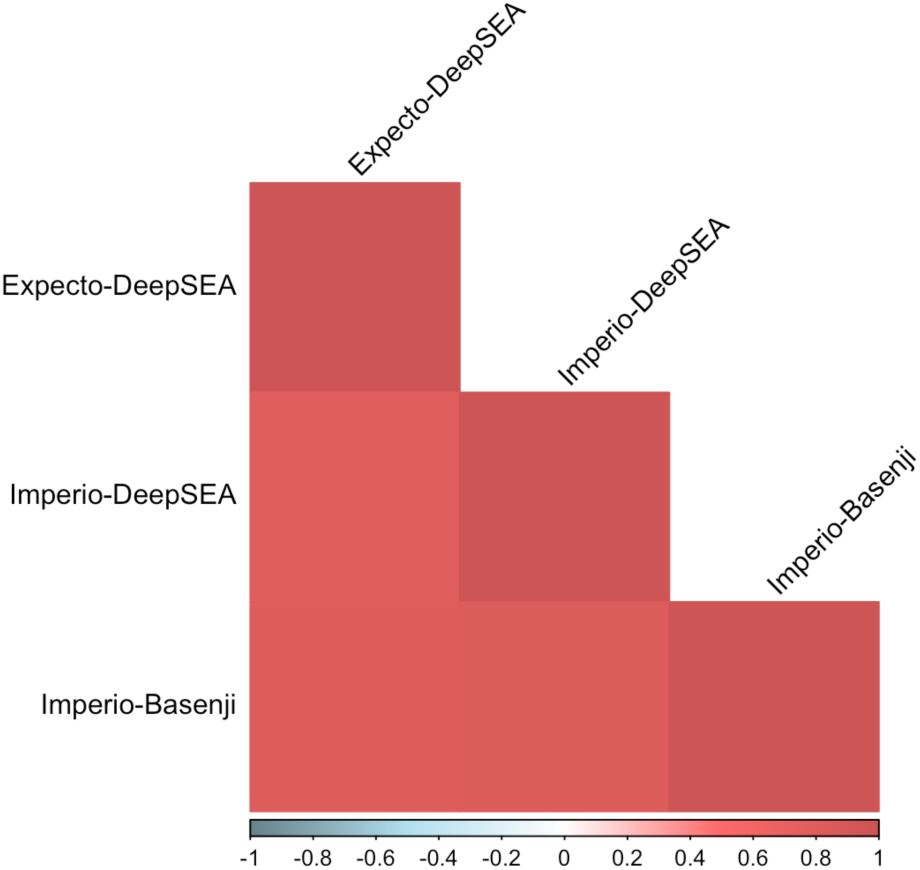
Correlation in predicted expression between Imperio and ExPecto models. Correlation in predicted expression for 990 chr8 genes used as held-out test set for the ExPecto method^4^ and the two Imperio models corresponding to DeepSEA and Basenji deep learning models.

**Figure S15.**
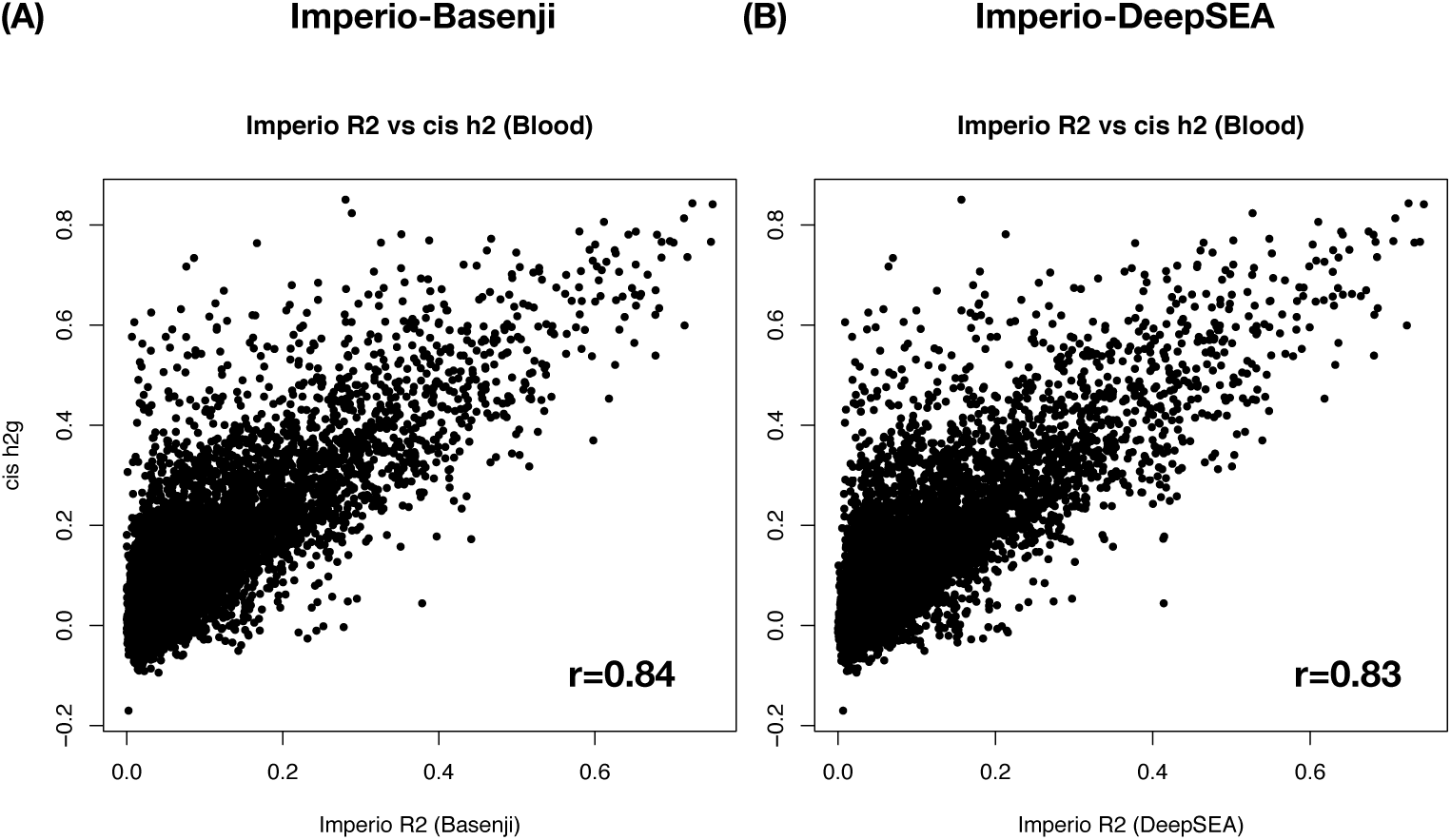
Comparison of Imperio prediction *r*^2^ for predictions of gene expression across individuals vs. cis-heritability. We plot the prediction *r*^2^ for the inter-individual model comprising of the Imperio predicted expression effects (see Methods) and the per-gene cis-heritability in Whole blood as estimated from trancriptiome wide association studies^52^ for two deep learning models - DeepSEA (Panel A) and Basenji (Panel B).

**Figure S16.**
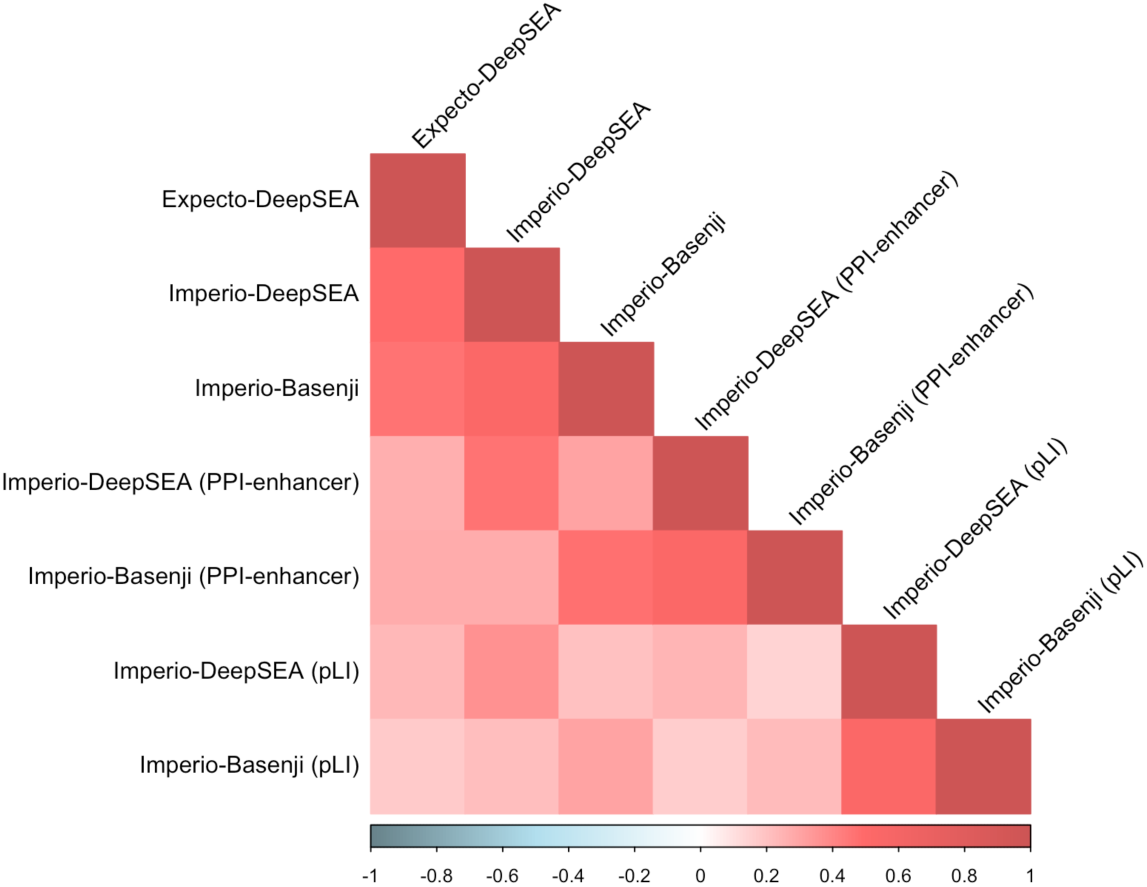
Correlation between all-genes ExPecto, all-genes Imperio, and gene set-specific Imperio annotations. Correlation matrix of Whole blood MaxCPP^27^, Whole blood ExPecto^27^, and 6 Imperio annotations corresponding to 2 deep learning models (DeepSEA and Basenji) and three sets of genes (all genes, pLI genes and PPI-enhancer genes). The correlations range from slightly positive to medium high positive values.

